# Selective identification of the herb *Picrorhiza kurroa* and probiotic *Lactobacillus fermentum* as a synbiotic with fermentation-enhanced physicochemical, biological, and metabolomic properties

**DOI:** 10.64898/2026.01.17.700043

**Authors:** Amit Kumar, Bhawna Diwan, Mohd Adil Khan, Ankita Awasthi, Ekta Bala, Praveen Kumar Vema, Rohit Sharma

## Abstract

**Purpose:** Although secondary metabolites of medicinal plants can modulate the gut microbiota, their prebiotic and synbiotic potential is not fully explored. This study aimed to identify novel medicinal plant-probiotic combination(s) that show prebiotic and synbiotic effects through mutual synergism and reciprocal interactions.

**Methods:** Ten Himalayan region medicinal plants were screened for their selective prebiotic efficacy by studying in vitro proliferation of three species of lactic acid bacteria (LAB) and concurrent suppression of *E. coli*. The top-performing plants were evaluated for synergism with LAB through fermentation of pure extracts, followed by physicochemical, biological, and metabolomic profiling.

**Results:** Among the ten plant species, *Picrorhiza kurroa* (PK) and *Adhatoda vasica* (AV) exhibited promising prebiotic attributes with PAS scores of 0.50 and 0.44, respectively, while *Lactobacillus fermentum* (LF) emerged as the most compatible probiotic strain. Unlike AV, fermentation of PK with LF demonstrated superior probiotic growth (2.2-fold increase), elevated total flavonoids (21.6% increase), and improved antioxidant capacity (24.02% increase) compared to unfermented PK. Fermented PK significantly enhanced cytoprotective effects and mitigated oxidative (ROS levels/lipid peroxidation) and inflammatory damage (NO/IL-6 levels) in macrophages and muscle cells exposed to separate exogenous stressors (LPS and H₂O₂). Elemental (ICP-OES) and metabolomic analysis (LC-MS/UHPLC) revealed that LF metabolised major elements of PK (Ca/Fe/K/Na/Cu) and induced biotransformation of large secondary metabolites, including the characteristic iridoid glycosides of PK, into smaller, more abundant metabolites.

**Conclusions:** PK-LF combination is identified as a synbiotic, and its fermented product is a superior bioactive product that can be developed into innovative nutraceuticals or herbal formulations.

## 1. Introduction

Medicinal plants are rich sources of unique bioactive secondary metabolites. Traditionally, these secondary metabolites have been viewed as anti-microbial agents. However, emerging studies indicate that they can also exert species-specific differential effects on the growth of gut microorganisms [1,2]. Specifically, metabolites such as polyphenols can suppress the proliferation of pathogenic bacteria while demonstrating growth-stimulatory properties on probiotic bacteria [3,4]. Therefore, due to these selective effects, plant bioactive compounds are being advocated as a second generation of prebiotics [1,5]. In addition, probiotic bacteria mediated utilisation of specific plant secondary metabolites often results in the biotransformation of native plant metabolites into simpler, more abundant, and bioactive compounds, thereby improving the plant’s in vivo efficacy [6,7]. Thus, a reciprocal and mutually beneficial interaction is plausible between probiotic bacteria and secondary metabolite-rich plants, which is otherwise lacking in traditional oligosaccharide-based prebiotics. Together, this provides opportunities to design novel synbiotic functional foods or herbal formulations, extending beyond conventional carbohydrate-based prebiotics [6].

The Himalayas are a biodiversity hotspot, and this region is rich in underutilised medicinal plants. Modern scientific studies are now demonstrating that these plants often contain specific metabolites that can be bioactive and pharmacologically significant. More importantly, plant metabolites interact with the gut microbiota and may also be used to treat gut microbial imbalances induced by various disorders [8]. On ingestion, plant metabolites are first acted upon by the gut bacteria, which not only aid in digestion and absorption, but the systemic effects of bioactive plant metabolites often appear to be dependent on their gut microbial interactions[9]. Therefore, understanding these reciprocal interactions and the possible amalgamation between plant secondary metabolites and gut microbiota is of great significance in food and herbal drug development [10]. However, the current knowledge about medicinal plants for their synergy and fermentability with probiotic bacteria is still limited[11]. As a result, the prebiotic and synbiotic potential of medicinal plants and/or their metabolites remains largely untapped [6,11,12].

Considering the foregoing discussion, this study systematically screened ten Himalayan medicinal plant extracts for potential prebiotic and synbiotic properties. The selective prebiotic potential of plants was identified using three probiotic lactic acid bacteria (LAB) strains, and the identified plant-LAB combination was subject to fermentation to develop a novel fermented product that was comprehensively evaluated for various physicochemical, bioactive, and metabolomic attributes. Our results identify the Himalayan herb *Picrorhiza kurroa* (PK) as a potential prebiotic agent that can be combined with probiotic *Lactobacillus fermentum* (LF) to yield a novel synbiotic herbal formulation with improved physicochemical profile and enhanced biological efficacy.

## 2. Materials and Methods

### 2.1. Plant material collection and phytochemical extraction

Ten different medicinal plants of the Himalayan and sub-Himalayan regions were used in this study. *Picrorhiza kurroa* (PK), *Podophylluym hexandrum, Rhododendron arboreum, Aconitum heterophyllum, Trillium govanianum,* and *Valeriana Jatamansi* were procured from Chamba (alt. 2500 m) and Kinnaur (alt. 2290 m) regions of Himachal Pradesh, India, while *Tinospora cordifolia, Adhatoda vasica* (AV), *Callistemon lanceolatus,* and *Glycerrihza glabra* were obtained from Solan, Himachal Pradesh (alt. 1467 m). The specific plant part utilised in the study is listed in Table 1, which was selected based on traditional usage and knowledge. The plant materials were shade-dried, powdered, and subjected to hydroalcoholic maceration (ethanol: water, 80:20). Briefly, ten grams of each powder was extracted twice (24 h and 48 h, respectively) with stirring, filtered, and concentrated under a rotary evaporator [13]. Extracts were dried, stored at −80 °C, and freshly prepared if degradation was observed.

**Table 1.** List of plants and their parts used in the study.

| S.no | Plant | Part used |
| --- | --- | --- |
| 1 | *Picrorhiza kurroa* | Roots |
| 2 | *Podophylluym hexandrum* | Roots |
| 3 | *Rhododendron arboreum* | Flowers |
| 4 | *Aconitum heterophyllum* | Roots |
| 5 | *Tinospora cordifolia* | Stem |
| 6 | *Trillium govanianum* | Rhizomes |
| 7 | *Glycerrihza glabra* | Rhizomes |
| 8 | *Adhatoda vasica* | Leaves |
| 9 | *Callistemon lanceolatus* | Leaves |
| 10 | *Valeriana Jatamansi* | Roots |

### 2.2. Culture and maintenance of microorganisms

Three commercial LAB strains: *Lactobacillus rhamnosus* GG (LR) (ATCC 53103), *Lactobacillus casei* (LC) (ATCC 393), and *Lactobacillus fermentum* (LF) (ATCC 9338), and one *Escherichia coli* (*E. coli*) strain (ATCC 4157) were used in this study, which were procured from HiMedia Laboratory, Mumbai, India. LAB strains were each cultivated in de Man, Rogosa, and Sharpe (MRS) broth and activated to the exponential phase before experiments. The growth profile of each LAB strain was monitored and standardised to accurately determine the target bacterial levels required in experiments. Similarly, *E. coli* was cultured and activated using Luria-Bertani (LB) broth.

### 2.3. Prebiotic activity and PAS

To assess prebiotic activity, different dried plant extracts were directly inoculated into each bacterial culture medium (containing 5 × 10^6^ CFU/mL) at four different concentrations: 0.25, 0.5, 2.5, 5, and 10 mg/mL, which are concentrations of extracts expected to be found under physiological conditions, as reported previously [3,14]. Negative control was kept without any plant extract. After aerobic incubation at 37 °C for 18 h at a constant stirring of 120 rpm in a shaking incubator, samples were collected, and optical density (OD) was immediately measured at 600 nm using a Varioskan Lux Microplate Reader (Thermo Fisher Scientific, USA). For bacterial colonies enumeration, samples were serially diluted in sterile normal saline and plated on MRS agar medium (for probiotic bacteria) and LB medium for *E. coli,* followed by incubation at 37 °C for 48 h. The prebiotic activity score (PAS) of each plant extract was calculated to accurately determine the prebiotic activity of plant extracts using the following equation, as reported before [15].

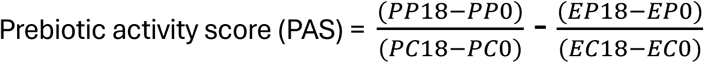

where, PP18 and PP0 are log_10_ CFU/mL of each LAB strain with plant extract obtained after 18 and 0 h of fermentation, respectively. PC18 and PC0 are log_10_ CFU/mL of each LAB strain without plant extract (control samples) obtained after 18 and 0 h of fermentation, respectively. Similarly, EP18 and EP0 are log_10_ CFU/mL of *E. coli* with plant extract obtained after 18 and 0 h of fermentation, respectively, while EC18 and EC0 are log_10_ CFU/mL of *E. coli* without plant extract (control samples) obtained after 18 and 0 h of fermentation, respectively.

### 2.4 Validation of prebiotic eLects using murine faecal microbiota

To further validate the prebiotic effects of identified plant extracts, an in vitro study on the growth of murine faecal LAB was conducted, as also reported previously [16]. Fresh faecal samples from male Swiss albino mice (4 months old; n=3), kept on a basal diet, were collected and homogenised in PBS (1:10 w/v) to make a slurry. Then, the slurry was inoculated (@ 0.1% v/v) into LAB-specific MRS medium with or without respective plant extracts (@5 mg/mL), followed by incubation at 37 °C for 18 h at a constant stirring rate of 120 rpm. Subsequently, bacterial CFUs were enumerated by plate count using MRS agar plates [16,17].

### 2.5 Plant extract fermentation

The plant extracts were freshly prepared (5 mg/mL), centrifuged to separate large insoluble particles, and the supernatant was subjected to filter sterilisation (0.22 µm) followed by inoculation with 1 × 10^8^ CFU/mL of the desired probiotic bacteria in a 500 mL Erlenmeyer flask, bringing the final volume to 200 mL. The flask was incubated at 37 °C for 24 h at 120 rpm in a shaking incubator. The plant extract without any added probiotic bacteria served as the negative control. Samples were recovered at 6, 9, 12, 24, and 48 h, aliquoted, and stored at -80°C for assessing various physicochemical and biological parameters as detailed below.

### 2.6 pH measurement and bacterial viability count

All samples were subject to pH measurement immediately after acquisition using Eutech pH 700 (Eutech Instruments Ltd., Singapore). Viable bacterial CFUs were enumerated using serial dilution and plating on MRS agar medium, followed by incubation at 37 °C for 48 h. All experiments were conducted in triplicate.

### 2.7 Estimation of total phenolic content (TPC)

TPC was determined in samples using the Folin-Ciocalteu method as reported previously[18]. A standard calibration curve was prepared using gallic acid, and the TPC of the samples is expressed as micrograms of gallic acid equivalents per mL.

### 2.8 Estimation of total flavonoid content (TFC)

TFC was estimated using the aluminium chloride colourimetric method as reported before, with slight modifications, using EGCG as a standard [19]. TFC is expressed as micrograms of EGCG equivalents per mL.

### 2.9 Estimation of total polysaccharides

Levels of total polysaccharides in samples were determined by the phenol–sulfuric acid method using anhydrous glucose as the standard [20]. The polysaccharide levels are expressed as µg/mL of glucose equivalent.

### 2.10 Estimation of total antioxidant capacity (TAC)

The total antioxidant capacity of the samples was assessed using the Ferric Reducing Antioxidant Power (FRAP) assay, using ascorbic acid as a standard[20]. Results are expressed as ascorbic acid equivalent (µg/mL).

### 2.11 Cell lines culture and treatment

The cytoprotective attributes of the fermented product were evaluated using two cell lines and two separate stressors. RAW264.7 murine macrophage and C2C12 murine myoblast cell lines were cultivated and maintained in DMEM supplemented with 10% FBS and 100 µg/mL of penicillin-streptomycin (Invitrogen, USA) at 37 °C in a 5% CO_2_ incubator. RAW264.7 cells were stimulated with LPS (1 µg/mL) for 24 h, while C2C12 cells were stimulated with H_2_O_2_ (200 µM) for 3 h to establish models of oxidative-inflammatory stress-induced cellular damage. Initial dose standardisation was performed, and the protective effects of test products were evaluated by co-incubation with the stressor for respective incubation periods.

### 2.12 Estimation of intracellular reactive oxygen species (ROS)

The levels of intracellular ROS were determined by fluorescence microscopy using 2′,7′-dichlorofluorescin diacetate (DCFH-DA) (Merck, D6883) redox probe as described previously [21]. Fluorescence quantification of captured images was performed using CellProfiler image analysis software (version 4.2.8). For flow cytometry, DCFH-DA-stained cells were subject to acquisition in an Attune NxT flow cytometer (Thermo Fisher Scientific, USA). At least 50,000 events per sample were captured and analysed as a histogram under the BL-1 channel. Relevant unstained and stained controls were used to establish the fluorescence threshold, and results were quantified using FlowJo software.

### 2.13 Estimation of nitric oxide (NO) production

NO levels in the culture supernatants were quantified and reported using a Griess reagent-assay kit (Cat.#G2930, Promega, Madison, WI, USA) as described previously [22].

### 2.14 Lipid peroxidation assay

The extent of stress-induced lipid peroxidation in cells was analysed using the Image-iT™ Lipid peroxidation kit (C10445, Thermo Fisher Scientific, USA) as per the manufacturer’s instructions. Fluorescence quantification of captured images on EVOS M5000 Imaging System (Thermo Fisher Scientific, USA) was performed using the CellProfiler^TM^ software (v 4.2.8). Results are expressed as the ratio of red/green integrated fluorescence intensities per image.

### 2.15 IL-6 estimation by ELISA

The levels of secretory IL-6 in the culture supernatants were determined through a sandwich ELISA kit (Cat no. 88-7064-88; Thermo Fisher Scientific, USA) as per the manufacturer’s instructions. The results for IL-6 levels are reported as per µg total protein.

### 2.16 Elemental content evaluation using ICP-OES

Levels of calcium (Ca), magnesium (Mg), potassium (K), phosphorus (P), copper (Cu), iron (Fe), sodium (Na), manganese (Mn) and zinc (Zn) were analysed through Inductively Coupled Plasma Optical Emission Spectrometry (ICP-OES) (Thermo Fisher, iCAP 7200 Duo series) and a microwave digester (Anton Paar 24 HVT50) at the Food Testing Laboratory of Shoolini Life Science Private Limited, Solan (India). One mL of liquid sample was mixed with 6 mL of concentrated nitric acid (HNO_3_), 0.5 mL of deionised water, and 0.5 mL of hydrogen peroxide (H_2_O_2_) and placed in the microwave digestor. The digestion program was set to heat the samples to 185°C, followed by holding at this temperature for 20-30 minutes before cooling. After digestion, the solutions were diluted with deionized water, and finally, the ICP-OES was calibrated with appropriate standards before quantifying the samples for elemental content.

### 2.17 Liquid chromatography–mass spectrometry (LC-MS)

LC-MS was performed using a Waters SYNAPT-XS HDMS system (Model: DBA064, Waters, UK) equipped with a UPLC ACQUITY H-Class separation module and controlled by MassLynx software (Version 4.2) at the Sophisticated Analytical Instrumentation Facility, Panjab University, Chandigarh (India). Chromatographic separation was achieved on an Acquity BEH C18 column (2.1 × 100 mm, 1.7 µm; Waters) with a mobile phase consisting of solvent A (0.1% formic acid in LC-MS grade water) and solvent B (0.1% formic acid in acetonitrile). The gradient program was set as follows: 0–5 min, 95% A; 5–30 min, linear gradient to 10% A; 30–35 min, 10% A; 35–36 min, return to 95% A; and 36–45 min, re-equilibration at 95% A. The flow rate was maintained at 0.2 mL/min, the injection volume was 5 µL, and the total run time was 45 min. Mass spectrometric detection was carried out in positive electrospray ionisation (ESI) mode under multiple reaction monitoring (MRM) with unit resolution. The source parameters were optimised as follows: capillary voltage 3.22 kV, cone voltage 50 V, source offset 80 V, source temperature 120 °C, desolvation temperature 550 °C, desolvation gas flow 950 L/h, cone gas flow 50 L/h, and collision energy 4 eV.

### 2.18 UHPLC analysis

UHPLC analysis was performed at the Food Testing Laboratory of Shoolini Life Science Private Limited, Solan (India), using an UltiMate 3000 UHPLC system (Thermo Fisher Scientific, USA) and analysed through Chromeleon chromatography data system software. Briefly, the extracts (1 mL) were filtered through a 0.2 µm filter, and 20 µL of each filtered extract was injected using a Dionex UltiMate 3000 autosampler. The solvent system used for gradient elution was a mixture of orthophosphoric acid (pH 3.5) (85%) and acetonitrile (15%). The acetonitrile concentration was increased to 30% for the first 5 min and to 50% over the next 10 min, then increased to 15% for the next minute and maintained for the next 20 min (total run time, 20 min). An analytical column (4.6 x 250 mm) with a packing material of 5 µm particle size at a flow rate of 1 mL/min at room temperature was used. During each run, the absorbance was recorded at 280 nm. Pure standards of gallic acid, catechin, epicatechin, quercetin, p-coumaric acid, caffeic acid, and resveratrol in 100% methanol were used to calibrate the standard curves and retention times.

### 2.19 Statistical analysis

Data are expressed as mean ± S.D. (n = 3). Significant differences among the groups were determined using one-way ANOVA followed by Tukey’s post-hoc test for multiple comparison corrections, as performed using GraphPad Prism statistical software (version 10.2). Differences between means were considered statistically significant at p ≤ 0.05. Principal component analysis (PCA) was performed and analysed using OriginPro statistical software (v 10.1.0.178).

## 3. Results

### 3.1 Prebiotic activity of herbal plant extracts and PAS

The influence of various plant extracts on the growth of probiotic bacteria is presented in Figure 1. It was observed that different plant extracts exerted species-dependent effects on bacterial growth. A close relationship between OD values and CFU counts was also noticeable. *Adhatoda vasica* (AV) plant extract appeared to be most favourable for promoting the growth of LC, while *Callistemon lanceolatus* and *Glycerrihza glabra* were most inhibitory (Figure 1A-B). On the other hand, a majority of plant extracts induced the growth of LF, which was particularly noticeable for *Adhatoda vasica*, *Picrorhiza kurroa*, *Valeriana jatamansi*, and *Trillium govanianum* (Figure 1C-D). LR growth was significantly increased in the presence of *Adhatoda vasica*, *Picrorhiza kurroa, Glycerrihza glabra*, and *Rhododendron arboreum* (Figure 1E-F). Among all the tested LAB strains, the growth of LC appeared to be the least affected by plant extracts, while LF appeared to be the most favoured. On *E. coli*, *Adhatoda vasica*, *Picrorhiza kurroa, Podophyllum hexandrum,* and *Glycerrihza glabra* exerted significant growth inhibitory effects, while *Valeriana jatamansi*, *Tinospora cordifolia*, and *Callistemon lanceolatus* were stimulatory (Figure 2A-B). The prebiotic activity of different plant extracts, as well as the suitability of various LAB as fermenting agents, was objectively assessed through the PAS. A PAS value greater than zero indicates that the examined compound selectively favours the growth of probiotic bacteria, while a negative PAS value is suggestive of increased pathogenic growth[15]. As shown in Figure 2C-E, LF categorically appeared to be the most effective strain since 7 out of 10 tested plants fermented with LF showed a positive PAS value. Further, the degree of a positive PAS value was significantly higher for LF, while the degree of a negative PAS value was lower compared to the remaining LAB strains. The highest PAS of 0.9 was observed for *Picrorhiza kurroa* in LF, indicating its strong prebiotic activity. The combined PAS of all LAB strains unambiguously indicated that *Picrorhiza kurroa* (PK) and *Adhatoda vasica* (AV) were the candidate prebiotics with an average PAS value of 0.50 and 0.44, respectively (Figure 2F). In vitro faecal fermentation of these plants also showed a strong and significant increase in total faecal LAB population compared to their respective controls (Figure 2G). Collectively, based on these results, PK and AV plant extracts were chosen as prebiotics, and LF was selected as the candidate compatible probiotic strain for further investigation.

**Figure 1.**
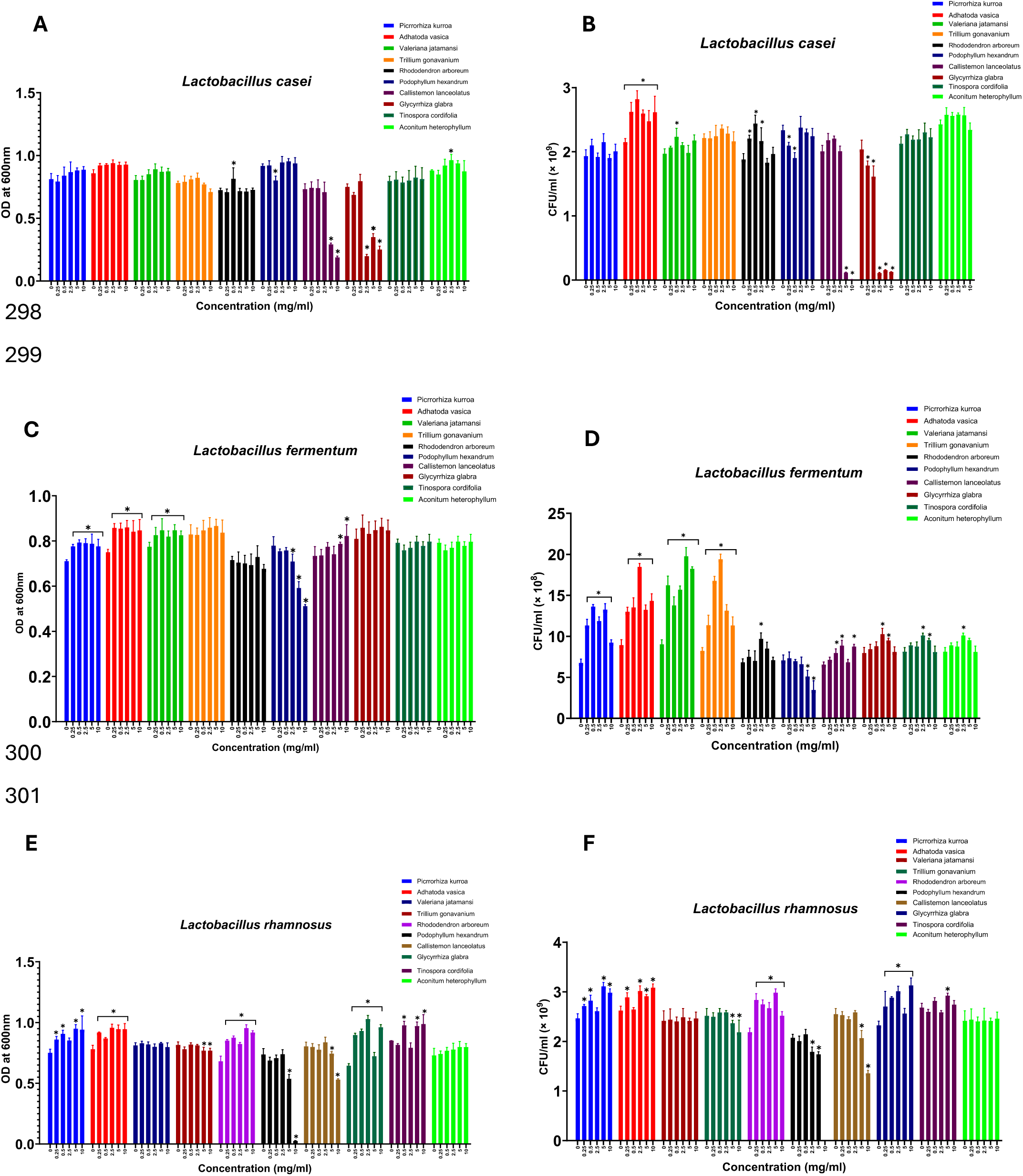
Effect of different plant extracts on the proliferation of **A-B** *Lactobacillus casei.* **C-D** *Lactobacillus fermentum.* **E-F** *Lactobacillus rhamnosus.* *represents a significant difference at p<0.05 compared to the control.

**Figure 2.**
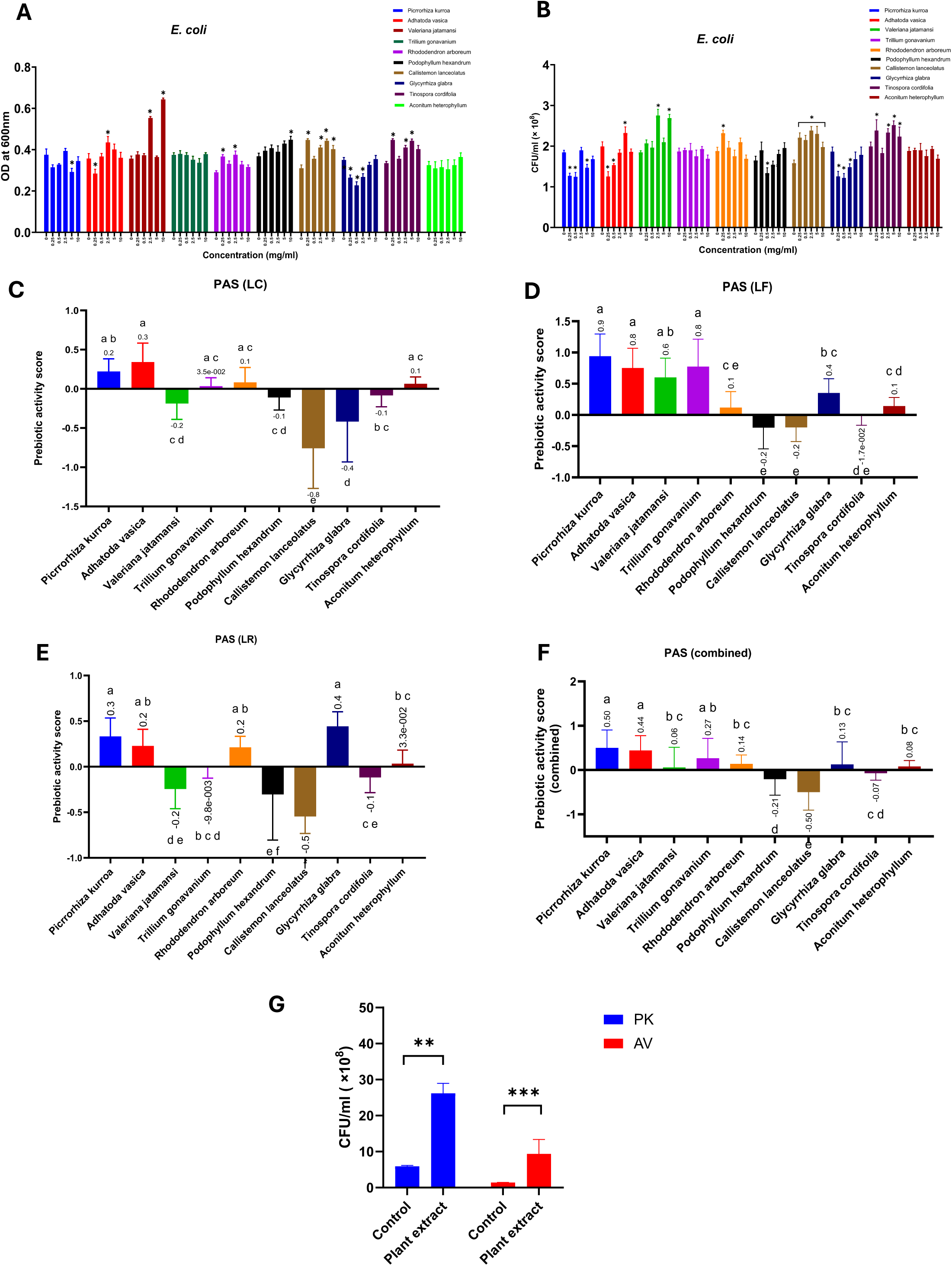
Effect of different plant extracts on the proliferation of **A-B** *Escherichia coli.* Prebiotic activity score (PAS) of plant extracts using **C** *Lactobacillus casei.* **D** *Lactobacillus fermentum.* **E** *Lactobacillus rhamnosus.* **F** Combined PAS using all LAB. **G** Effect of PK and AV on the proliferation of murine faecal LAB. *represents a significant difference compared to the control. *p<0.05, **p<0.01, ***p<0.001. Means that do not share a common letter indicate a significant difference at p<0.05.

### 3.2 Physicochemical analyses and correlation evaluation

*Trillium govanianum*, *Rhododendron arboreum*, and *Callistemon lanceolatus* recorded the highest TPC levels among all tested plant extracts (Figure 3A), while *Aconitum heterophyllum*, *Tinospora cordifolia*, and *Callistemon lanceolatus* had the highest TFC levels (Figure 3B). *Trillium govanianum, Valeriana jatamansi,* and *Aconitum heterophyllum* exhibited the highest total polysaccharides (Figure 3C), while *Tinospora cordifolia* and *Rhododendron arboreum* recorded the highest antioxidant capacity (Figure 3D). Interestingly, in all these parameters, PK and AV always exhibited moderate activity and were never among the highest or lowest recorded values. As noted in Table 2, Spearman correlation analysis (n = 10) revealed that TPC was strongly and significantly associated with antioxidant capacity (r=0.76), although it showed no clear association with PAS. Instead, total sugar levels demonstrated a moderate but non-significant positive correlation with PAS (r = 0.49). Similarly, TAC in plants was also weakly correlated with PAS, albeit non-significantly. TFC showed a weak negative correlation (r=-0.224) with PAS, suggesting the anti-microbial effects of flavonoids.

**Figure 3.**
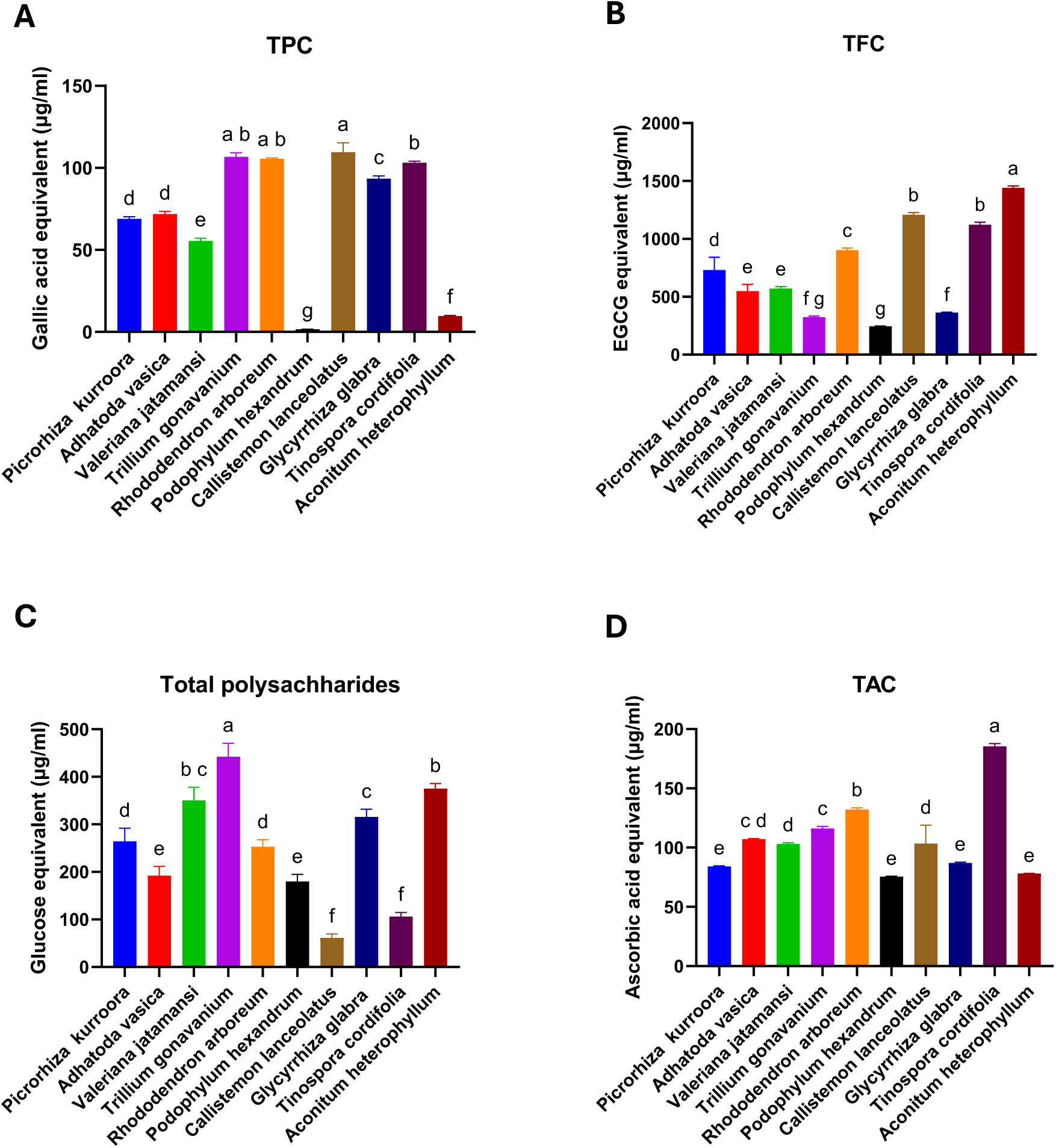
Analysis of physicochemical parameters of plant extracts. **A** Total phenolic content (TPC). **B** Total flavonoid content (TFC). **C** Total polysaccharide content**. D.** Total antioxidant capacity (TAC). Means that do not share a common letter indicate a significant difference at p<0.05.

**Table 2:** Spearman correlation analysis between plant physicochemical attributes and PAS.

| *r* | *TPC* | *TFC* | *TAC* | *Total Sugars* | *PAS* |
| --- | --- | --- | --- | --- | --- |
| *TPC* | 1 | 0.187 | 0.757* | -0.212 | 0.018 |
| *TFC* | 0.187 | 1 | 0.175 | -0.200 | -0.224 |
| *TAC* | 0.757* | 0.175 | 1 | -0.200 | 0.115 |
| *Total Sugars* | -0.212 | -0.200 | -0.200 | 1 | 0.490 |
| *PAS* | 0.018 | -0.224 | 0.115 | 0.490 | 1 |

*r*=Spearman correlation
\**p*<0.05

### 3.3 Identification of the synbiotic combination through fermentability evaluation

#### 3.3.1 Changes in pH and bacterial CFU

Pilot runs were conducted in which fermentation was performed for up to 48 h, and the viable bacterial count was measured at 0, 24, and 48 h. A strong suppression in viable bacterial count was observed after a 24-48 h fermentation period (data not shown). Thus, a smaller fermentation timeline (< 24 h) was determined optimal. The effects of LF-mediated fermentation on PK and AV are presented in Figure 4. It was observed that the pH of both PK and AV fermented extracts decreased rapidly within the first 6 h of fermentation, and then slowed, eventually reaching a near plateau after 9 h (Figure 4A). However, the degree of pH decrease was more noticeable in PK at all tested timepoints. A similar trend was evident in viable LF count, which significantly increased by over 2.2-fold after 6 h fermentation in both PK and AV compared to the control, but then showed a gradual decline to almost a non-viable state after 24 h (Figure 4B). Notably, LF viability was higher in PK than in AV at all tested time points.

**Figure 4.**
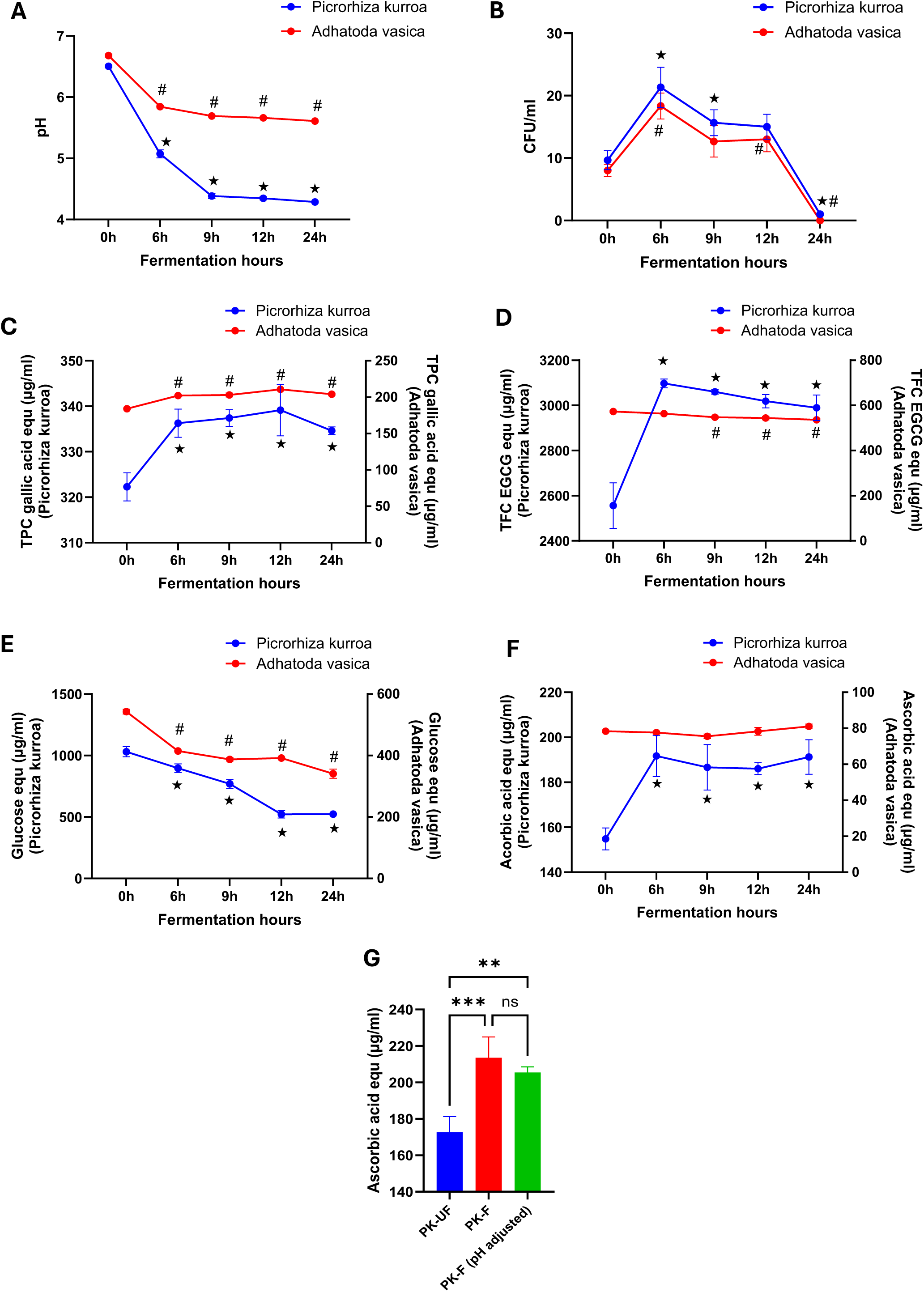
Characterisation of the fermentation of PK and AV with LF. Changes in **A** pH. **B** Bacterial proliferation. **C** Total phenolic content (TPC). **D** Total flavonoid content (TFC). **E** Total polysaccharide content**. F** Total antioxidant capacity (TAC). **G.** Effect of pH neutralisation on TAC of fermented PK. * and # represent a significant difference (p<0.05) compared to the control in PK and AV, respectively. *p<0.05, **p<0.01, ***p<0.001.

#### 3.3.2 Changes in total phenolics and total flavonoids

TPC significantly increased after fermentation in both PK and AV and recorded higher levels than control at all measured time-points (Figure 4C). Specifically, in fermented PK, TPC recorded the highest significant increase of 5.27% at 12 h of fermentation, while AV recorded the highest increase of 14.1% in TPC after 12 h of fermentation. However, TFC levels significantly increased only in fermented PK, while a significant decline in TFC was recorded in AV fermentation (Figure 4D). Specifically, a 21.16% increase in TFC was observed after 6h-fermentation in PK, and a 6.45% decrease in TFC was recorded in AV fermentation after 24 h.

#### 3.3.3 Changes in total polysaccharides

Both PK and AV fermentation resulted in a time-dependent significant decrease in polysaccharide content (Figure 4E). A 49.36% decrease in polysaccharide levels of PK was recorded after 24 h of fermentation, while a 37.26% decrease was observed during AV fermentation after 24 h.

#### 3.3.4 Changes in total antioxidant capacity (TAC)

TAC significantly increased in fermented PK at all tested time-points. Conversely, no significant changes in TAC were observed during AV fermentation (Figure 4F). Particularly, a 24.02% increase in TAC was observed in PK after fermentation for 6 h, which was also the highest among all tested timepoints. Interestingly, there was no significant difference in TAC after neutralisation of the acidity of the fermented PK (Figure 4G).

#### 3.3.5 Principal component analysis (PCA)

As noted in Figure 5A, PC1 and PC2 captured most (85.6%) of the total variations based on key parameters. A distinct clustering of fermented and unfermented PK and AV samples was evident, driven primarily by differences in TAC, TPC, and TFC (Figure 5A). PK-fermented samples clustered on the positive PC1 axis, linked to elevated TPC, TFC, TAC, sugars, and microbial viability, indicating enhanced bioactive release and growth. Conversely, AV fermented samples grouped on the negative PC1 axis, correlating with higher pH levels, suggesting slower acidification and bacterial proliferation. Correlation analysis confirmed strong positive associations among TPC, TFC, and TAC (r = 0.987–0.997), moderate correlations with sugars (r = 0.649–0.697), and negative correlations with pH (r = –0.663 to –0.706) (Figure 5B). Overall, PK fermentation by LF was superior to AV, with the 6 h PK–LF condition showing the best bacterial viability and physicochemical attributes. This combination was selected for subsequent synbiotic analysis.

**Figure 5.**
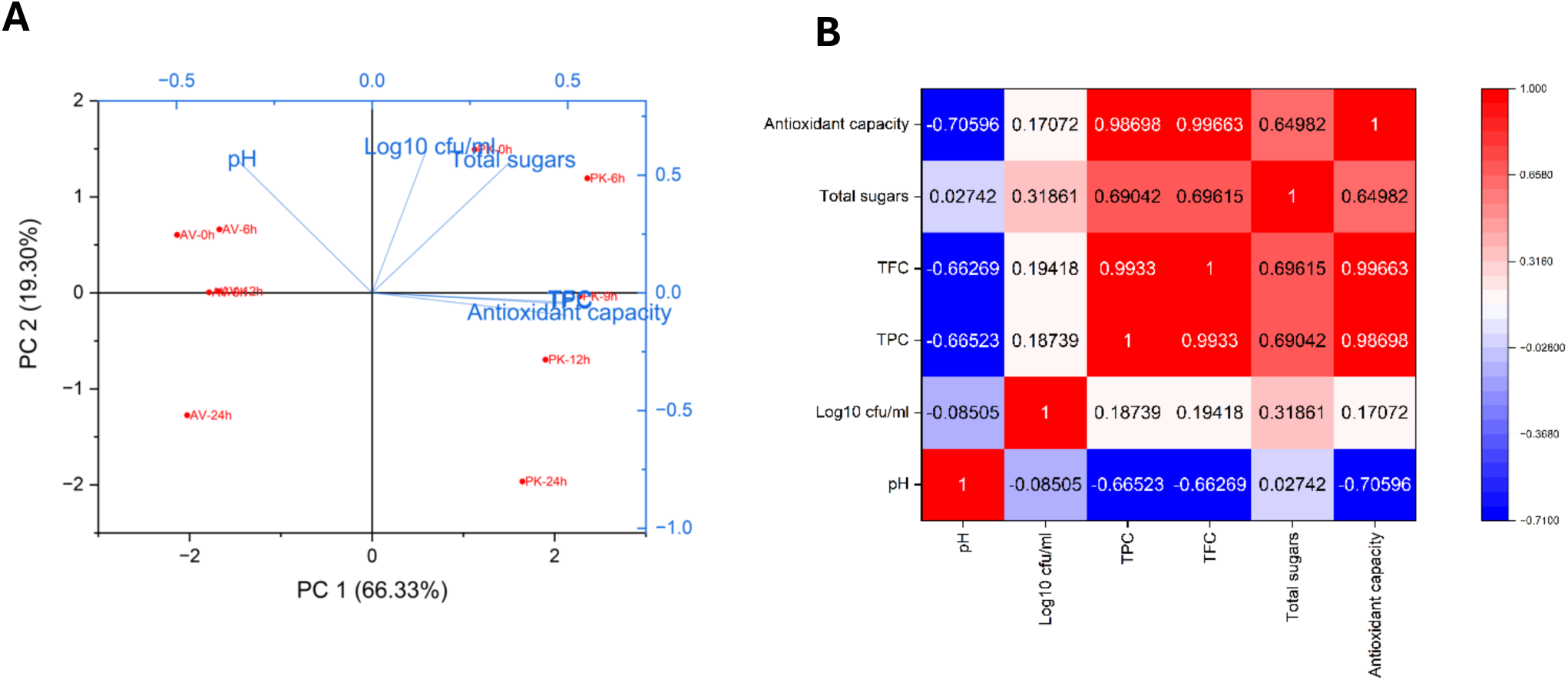
**A** Principal component analysis (PCA) of various fermentation parameters. **B.** Correlation matrix.

### 3.4 Biological activities evaluation

#### 3.4.1 NO production in LPS-induced macrophages

RAW264.7 macrophages were co-exposed to LPS and fermented (PK-F) and unfermented PK (PK-UF), respectively, for the analysis of anti-inflammatory and antioxidant attributes. During standardisation, plant extracts induced cytotoxicity at an exposure of ≥40% (v/v), and thus a dosage of 10% and 20% (v/v) was selected for further assessment. It was observed that NO production significantly increased by over 2-fold in LPS-treated macrophages, which was attenuated by the exposure to PK-UF and PK-F at both the tested concentrations (@10 and 20% v/v) (Figure 6A). However, PK-F clearly induced a more significant effect, with a 21.7% decrease in NO levels compared to PK-UF (20% v/v) (Figure 6A). Furthermore, the level of NO was comparable to the control only after treatment with PK-F (20% v/v).

**Figure 6.**
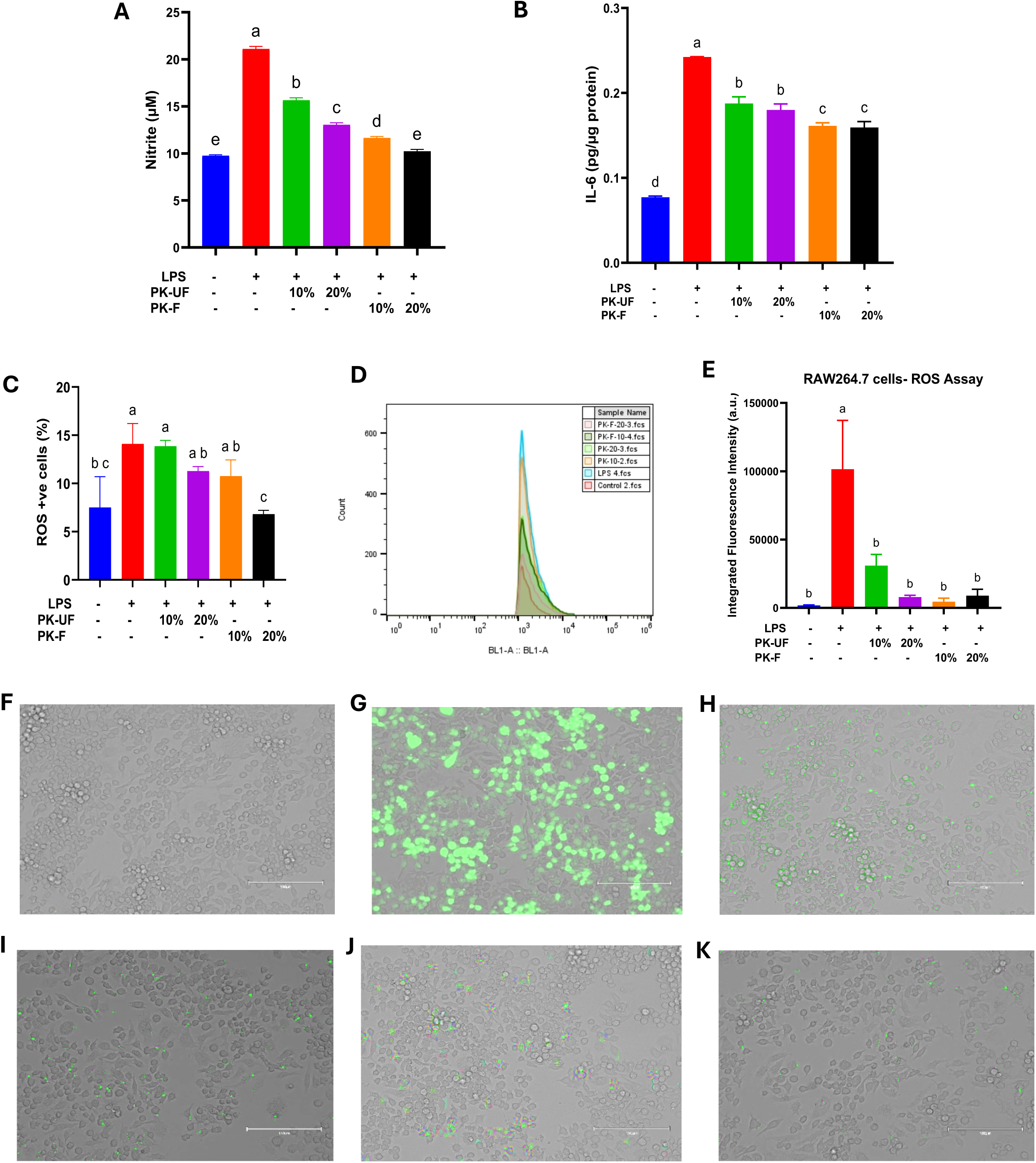
Antioxidant and anti-inflammatory effects of fermented PK (PK-F) in LPS-induced macrophages compared to unfermented PK (PK-UF). Levels of **A** NO. **B** IL-6. **C** No. of ROS+ cells determined by flow cytometry. **D.** Histogram plot. **E.** Integrated fluorescence intensity determined from fluorescence images. Representative fluorescent images of ROS+ cells in **F** Control. **G** LPS-treated cells. **H** LPS+PK-UF (10% v/v). **I** LPS+PK-UF (20% v/v). **J** LPS+PK-F (10% v/v). **K** LPS+PK-F (20% v/v). Means that do not share a common letter indicate a significant difference at p<0.05.

#### 3.4.2. IL-6 levels in LPS-induced macrophages

The levels of IL-6 in LPS-treated macrophages increased by over 2.4-fold compared to the control, which were significantly decreased by treatment with both PK-UF and PK-F (Figure 6B). However, a significant 16.6% more suppression in IL-6 levels was evident in cells treated with PK-F (@20% v/v) compared to PK-UF (@20% v/v) (Figure 6B). None of the tested treatments, however, matched the control.

#### 3.4.3. ELect on intracellular ROS levels in LPS-induced macrophages

Almost a 2-fold significant increase in cells positive for ROS was observed in LPS-treated cells through flow cytometery (Figure 6C). Although all tested extracts induced a decrease in the number of ROS+ cells, only the treatment with PK-F resulted in a significant decline of 51.4% in ROS+ cells compared to LPS-treated cells, which was also comparable to the control (Figure 6C). A representative histogram of various treatments is presented in Figure 6D. These observations were also corroborated by fluorescence microscopy analysis, which revealed a 55-fold significant increase in integrated fluorescence intensity in LPS-treated cells, which was significantly attenuated by treatment with both PK-F and PK-UF, although PK-F appeared to be more effective even at lower concentrations (Figure 6E). Representative fluorescent images are presented in Figure 6F-K.

#### 3.4.4. ROS production under oxidative stress in C2C12 cells

As noted in Figure 7, treatment with H_2_O_2_ induced a robust near 4-fold increase in ROS+ fluorescence intensity compared to the control. This was ostensibly attenuated by treatment with both PK-F and PK-UF. However, the effect was significantly more pronounced in PK-F, which appeared to be more effective at a lower concentration (@10%) (Figure 7B). Representative images are presented in Figure 7A.

**Figure 7.**
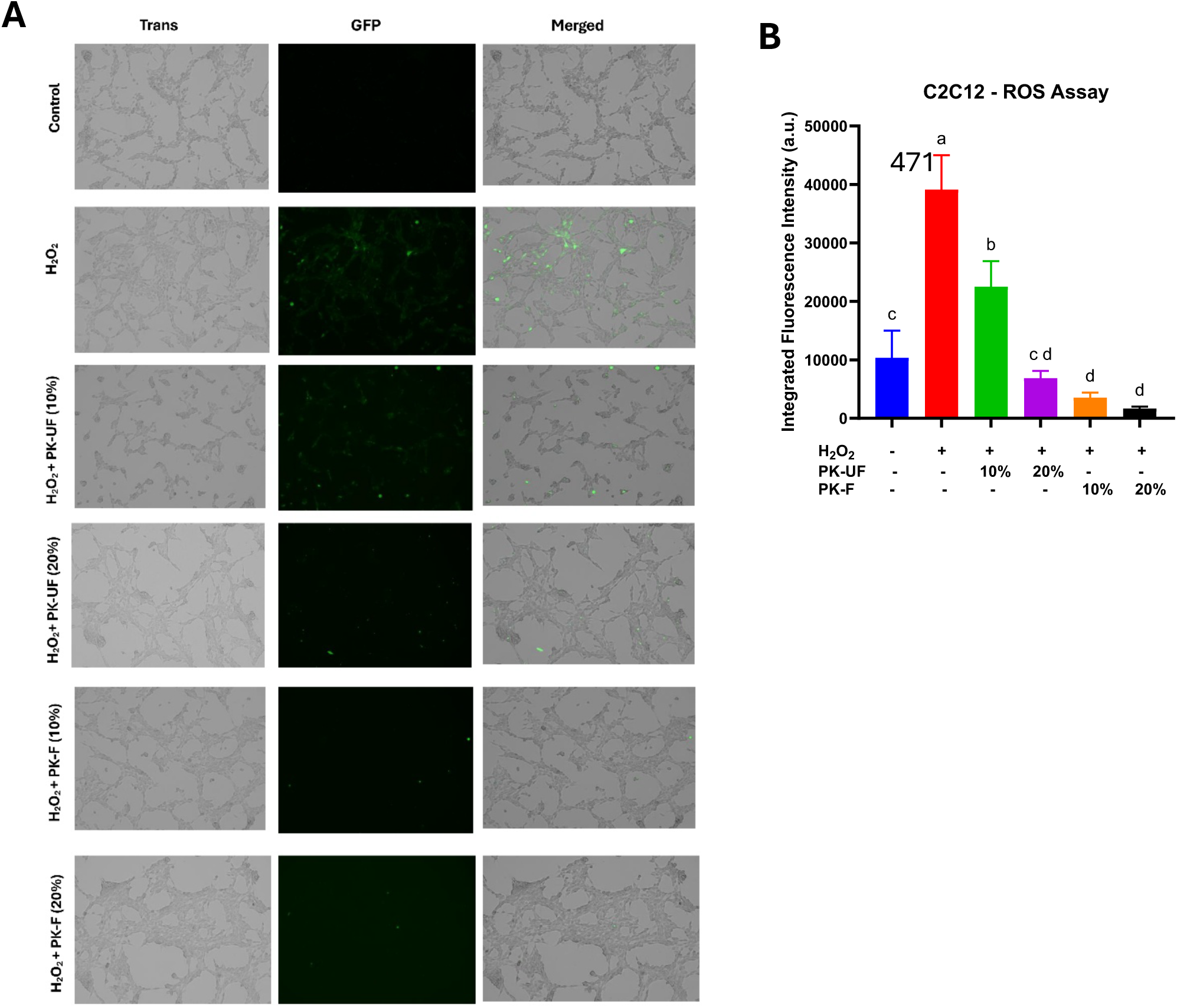
Influence of fermented PK (PK-F) in mitigating H_2_O_2_-induced ROS production in C2C12 cells compared to unfermented PK (PK-UF). **A** Representative fluorescence images **B.** Integrated fluorescence intensity. Means that do not share a common letter indicate a significant difference at p<0.05.

#### 3.4.5. Lipid peroxidation in C2C12 cells

As shown in Figure 8, the ratio of reduced/oxidised lipids significantly decreased by about 6-fold in H_2_O_2_-treated cells, indicating prevalent oxidation of cellular lipids (Figure 8B). On the other hand, treatment with PK-F appeared to moderately protect against lipid peroxidation, especially at a 10% (v/v) concentration. Representative images are presented in Figure 8A.

**Figure 8.**
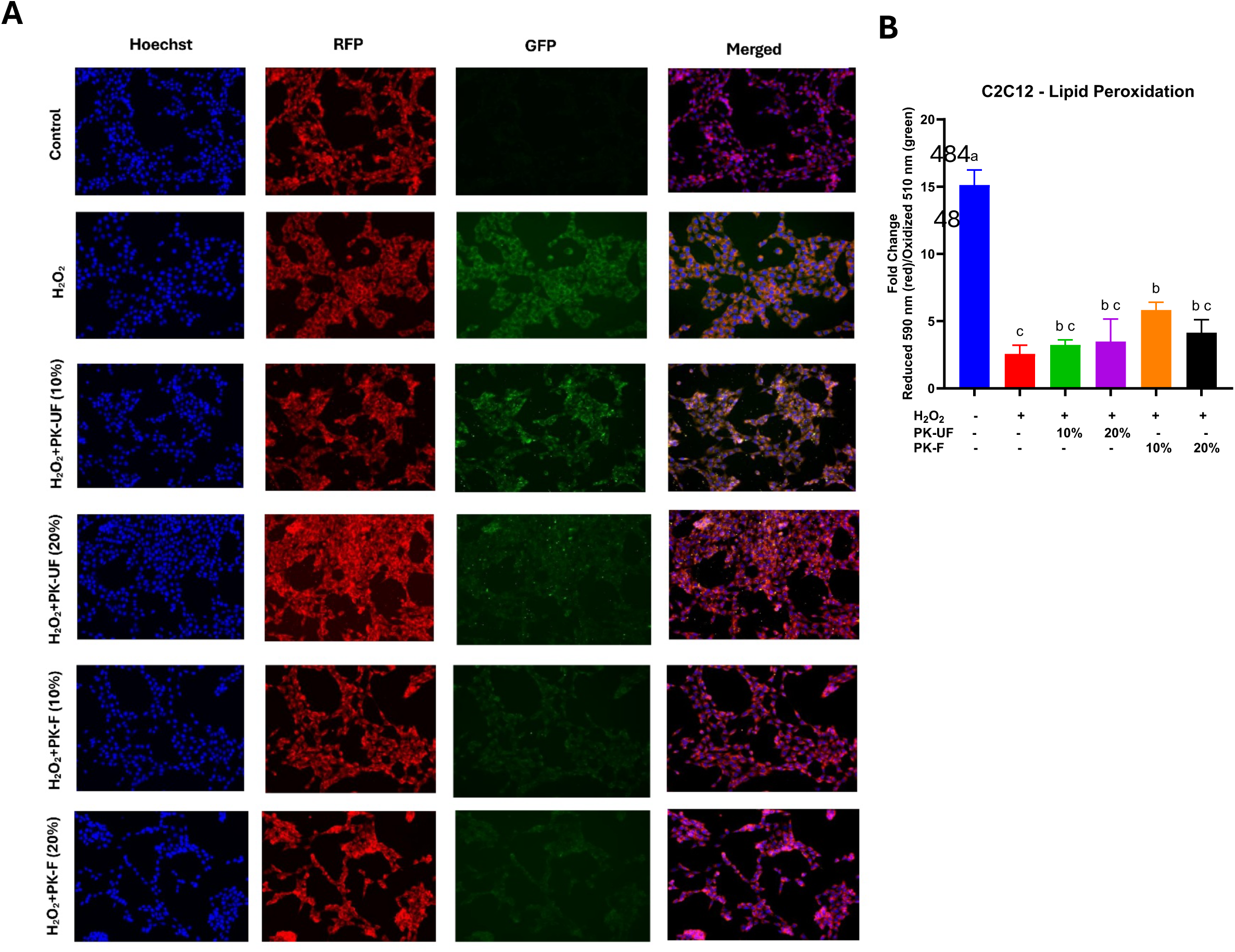
Effect of fermented PK (PK-F) in mitigating H_2_O_2_-induced lipid peroxidation in C2C12 cells compared to unfermented PK (PK-UF). **A** Representative fluorescence images **B.** Integrated fluorescence intensity. Means that do not share a common letter indicate a significant difference at p<0.05.

### 3.5 Elemental analysis

The levels of almost all tested elements decreased in the fermented supernatant (Table 3). Specifically, calcium, iron, and potassium in the fermented supernatant were reduced the most by 25.19%, 37.23%, and 46.05%, respectively. Similarly, sodium and copper decreased by 9.9% and 17.24%, respectively, while manganese levels slightly increased (Table 3).

**Table 3:**
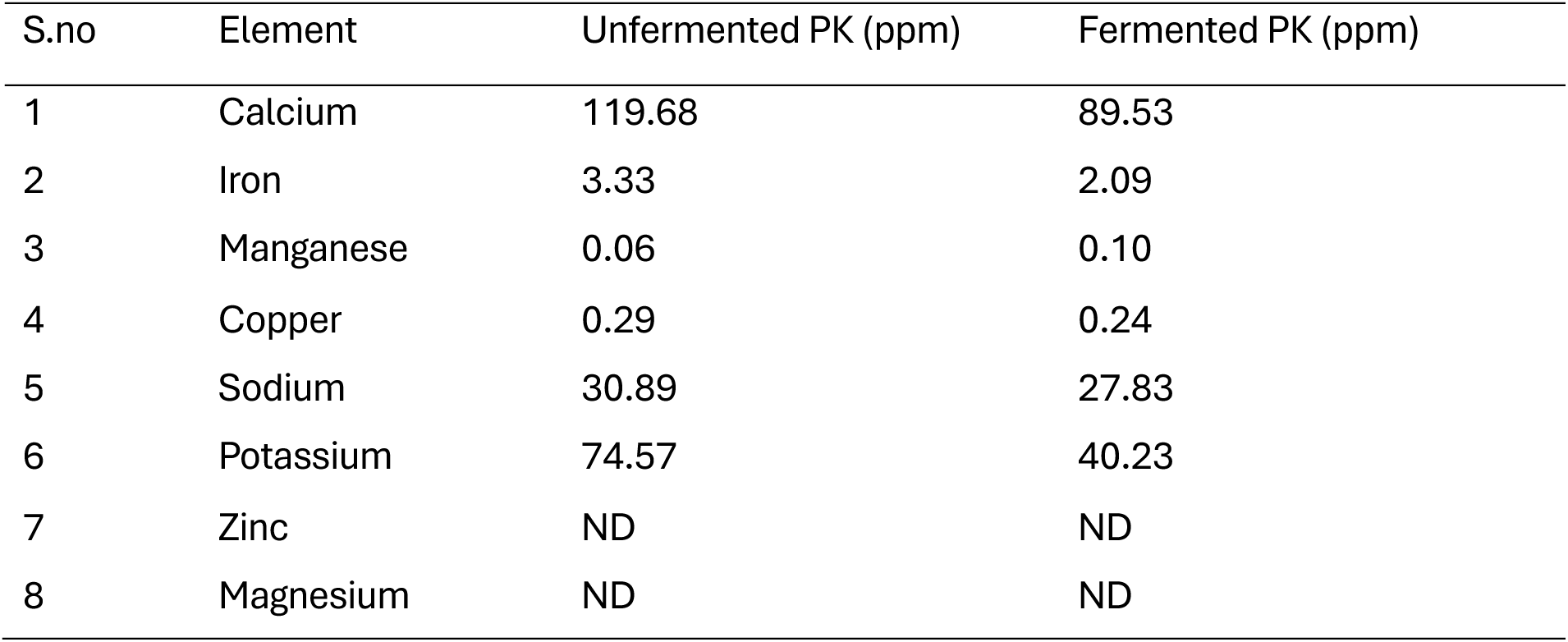
EUect of fermentation on major elemental composition of PK extracts.

### 3.6 LC-MS metabolite analysis

LC-MS chromatograms of PK-UF and PK-F are presented in Figure 9. In general, fermentation induced notable shifts in the metabolite profile of PK, with a decrease in higher mass compounds, such as m/z 535.2527, 663.3329, 515.2566, and 731.5596, and increases in dominant peaks at relatively lower m/z values, such as 355.1421. In addition, a new metabolite at m/z 275.0541 (RT: 15.97) was also prominent in the fermented sample, likely a fermentation by-product, reflecting overall catabolic activity and biotransformation. The characteristic metabolites of PK were identified in the chromatograms, and their relative concentrations (as interpreted through % of area under the respective peaks) were analysed, which appeared to decrease after fermentation. Specifically, the levels of Picroside I, Kutkoside, Picroside II, Picroside III, and Catalpol decreased from 11.5% to 10.06%, 3.05% to 2.24%, 1.96% to 1.59%, 9.05% to 8.92%, and 5.89% to 3.31%, respectively, after fermentation. The complete chromatograms, along with a list of specifically identified metabolites, are presented in the supplementary Table 1 and supplementary Figures 1-2.

**Figure 9.**
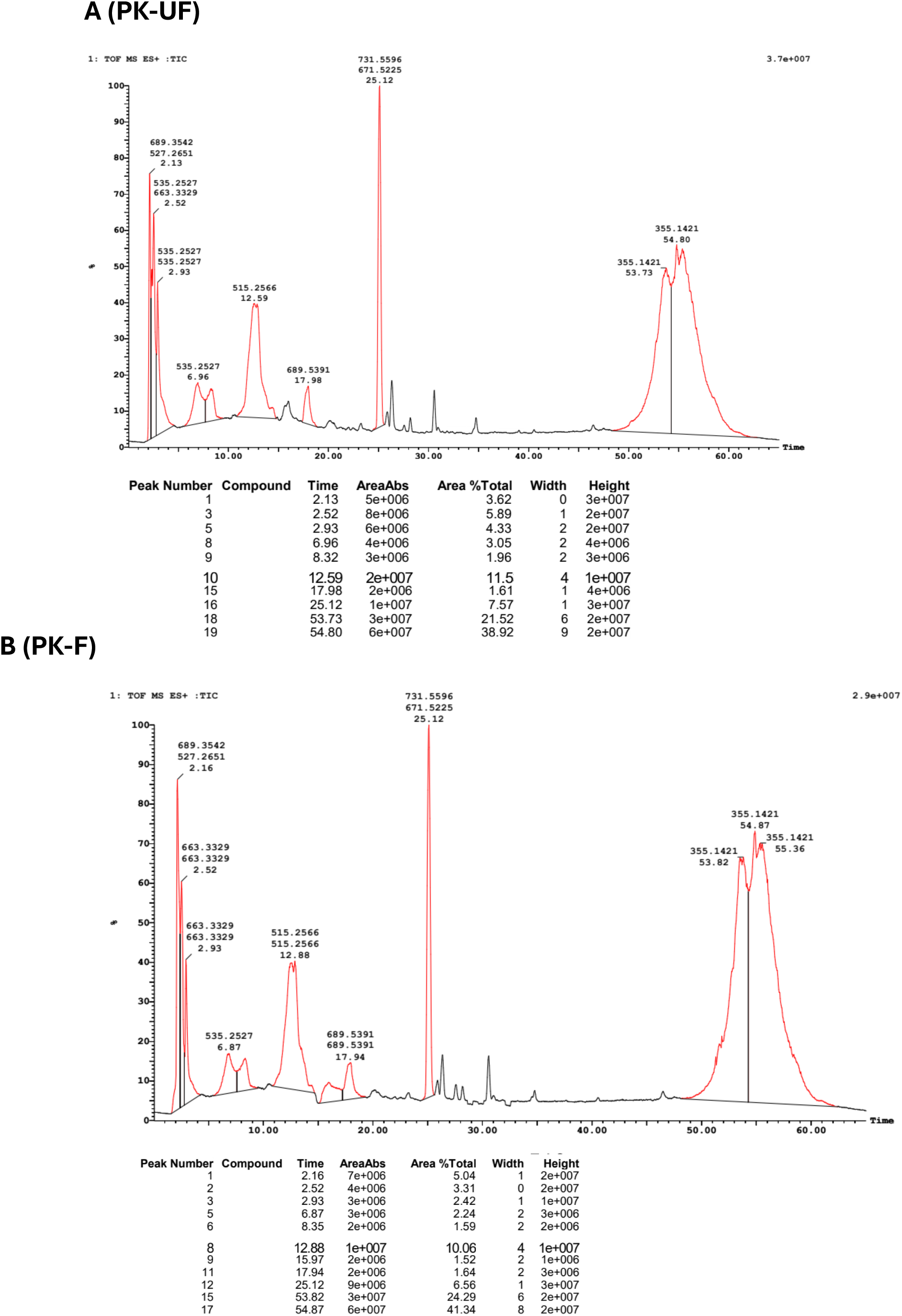
LC-MS chromatograms of **A** Unfermented PK (PK-UF). **B.** Fermented PK (PK-F).

### 3.7 UHPLC analysis

It was observed that fermentation strongly influenced the levels of all tested metabolites. As shown in Figure 10A, the abundance of chlorogenic acid and rosmarinic acid increased by 15.38% and 20.79%, respectively, in PK-F compared to PK-UF. Conversely, caffeic acid, rutin, and ferulic acid levels decreased in PK-F by 92.8%, 76.7%, and 26%, respectively. Representative HPLC chromatograms are presented in Figure 10B-C.

**Figure 10.**
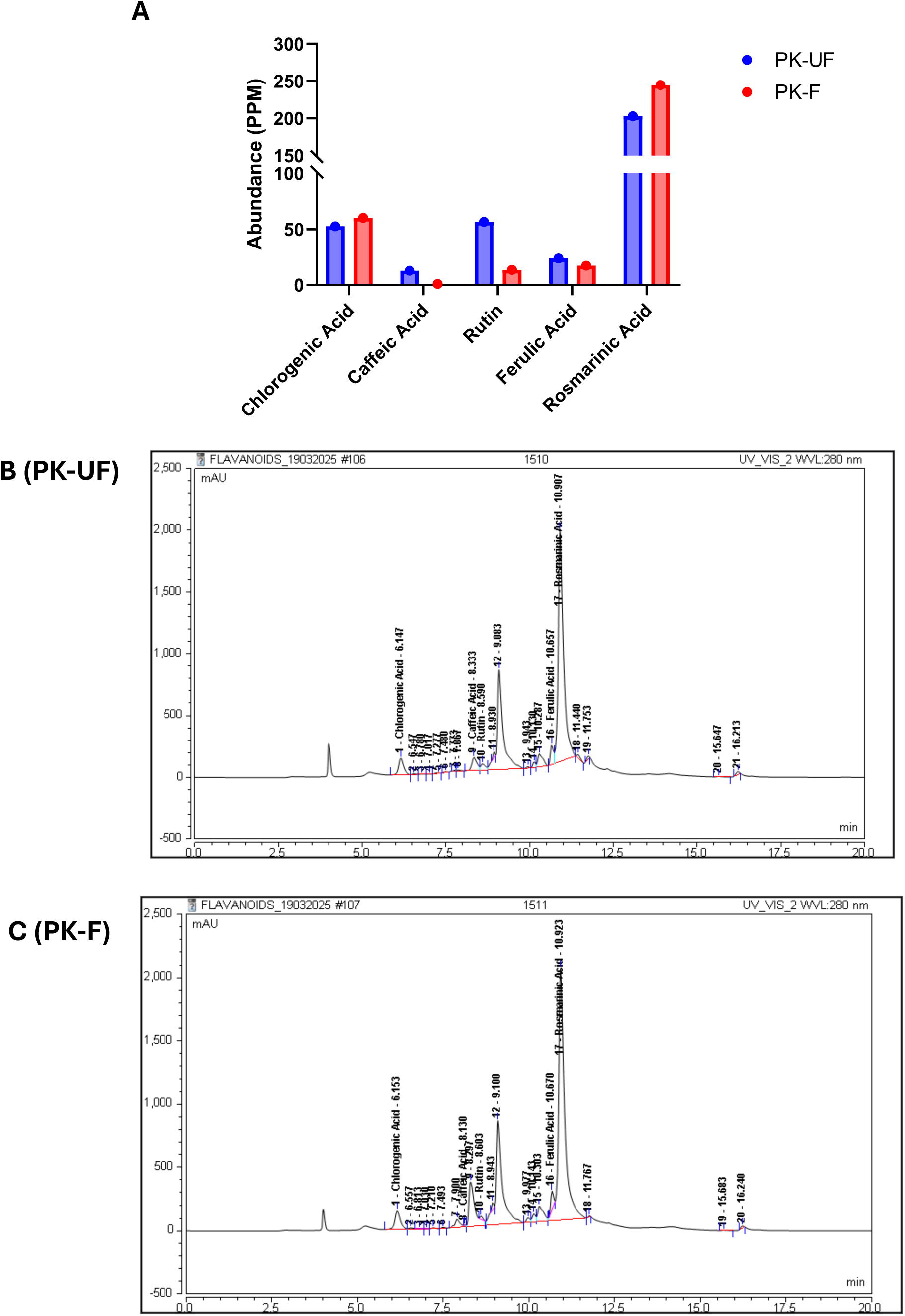
UHPLC characterisation of the fermentation of PK. **A** Relative abundance of metabolites after fermentation. UHPLC chromatograms with retention times of **B** Unfermented PK (PK-UF). **C** Fermented PK (PK-F).

## 4. Discussion

The present study tested several medicinal plants, but not all of them showed prebiotic activity. Correlation analysis revealed that none of the factors, such as TPC, TFC, TAC, or polysaccharide content in plants, could directly predict their prebiotic efficacy, indicating that plant prebiotic potential is not governed solely by bulk levels of sugars, phenolics, or antioxidants, but rather by a complex interplay of specific metabolite classes. PK is rich in sugar-conjugated iridoid glycosides, such as picrosides, which can be metabolised by probiotic bacteria through the action of specific enzymes. Similarly, AV is rich in pyrroloquinazoline alkaloids, saponins, tannins, and phenolic glycosides, which may act as mild prebiotic substrates enabling probiotic growth. In previous studies, species-specific effects on bacterial growth, ultimately favouring probiotic proliferation, were observed in polyphenol and flavonoid-rich foods such as green tea [4,23], blueberry [24], grapes [25], pomegranate extract [26], as well as in various medicinal plants [27,28]. Our study, however, provides the first evidence that PK and AV can be potential prebiotic agents.

The observed probiotic LAB tolerance and growth in the presence of secondary metabolite-rich PK and AV extracts may be attributed to the diverse and specialised polyphenol-metabolising enzymes in the genomes of LAB compared to opportunistic pathogens such as *E. coli* [29,30]. However, like plant extracts, not all tested LAB were equally effective in this study. Specifically, LF appeared as the major probiotic proliferating in the presence of most tested plant extracts. LF possesses several polyphenol-metabolising enzymes, including feruloyl esterases, phenolic acid reductases, β-glycosidases, and α-L-rhamnosidases, which may enable the bacterium to detoxify and metabolise complex phytochemicals [31]. Due to this, LF has been previously utilised in non-dairy-based fermentation processes involving plant extracts [32–34]. Given that LF showed a positive PAS value in 7 out of 10 tested plant extracts in our work, it suggests its suitability for designing prebiotic activity studies, as also noted before [35–38].

Apart from preservation, the ancient process of fermentation is currently emerging as a tool for the enhancement of nutritional properties and the development of novel functional foods, especially using various fruits [39,40]. However, systematic studies on the fermentation of crude medicinal plants remain relatively scarce. In our study, although both PK and AV appeared strongly prebiotic, fermentation of only PK was ostensibly favoured by LF. Notably, fermentation enhanced TFC and TAC, which are some of the desirable characteristics of a successful fermentation, only during the fermentation of PK, but not AV. The apparent disparity could be attributed to various reasons, including the different and complex plant metabolite profiles and LF fermentability characteristics. It is plausible that PK metabolites are better substrates for LF-mediated fermentation, and although AV acts as a prebiotic, its fermentation resulted in different end products that were less biotransformed into flavonoids, resulting in a minimal increase in TFC and TAC despite fermentation. Supporting our observations, it was reported that fermentation of *Cabernet sauvignon* grapes with *Aspergillus carbonarius* increased the antioxidant activity, but did not affect the antioxidant activity when fermented with *Moscato italico* grapes, thus demonstrating that fruit variety has an impact on fermentation outcomes [41]. Similarly, fermentation of Avocado leaf extracts demonstrated strain-dependent effects of LAB on antioxidant capacity [42]. Taken together, our observations imply that a positive prebiotic activity of a medicinal plant may not necessarily be considered amenable to desirable fermentation or synergism. Nevertheless, due to its observed prebiotic attributes, AV should be considered a viable candidate for addressing gut microbial imbalance.

Interestingly, a significant increase in TFC (21%) and TAC (24%) was observed within just 6 h of fermentation of PK with LF. Previously, similar or even lesser magnitudes of improvement have often been reported in other plant matrices, and they typically require a longer (12–72 h) fermentation period [20,43–45]. Therefore, unlike most studies, our data show that PK fermentation achieved significant increases in TFC and TAC within a shorter fermentation period (6 h), underscoring the rapidity and efficiency of this system. To delineate the cause of improved TAC, the pH difference between fermented and unfermented PK was eliminated, which demonstrated no significant decrease in the TAC. This indicated that the increase in TAC could primarily be due to observed changes in TPC/TFC rather than organic acids production by LF. Further, the improved physicochemical profile of fermented PK also translated into superior cytoprotective effects during in vitro evaluation. Previous studies have reported that fermentation of plant parts as well as specific plant metabolites can enhance their biological efficiency, as observed in the fermentation of okara [46], hempseed [47], horse gram extract [48], stevia leaf extract [49], and theabrownins[50]. Thus, LF-mediated fermentation enhanced the medicinal value of PK.

A general decrease in the levels of various elements, including calcium, sodium, and potassium, was noticed in fermented PK, which were likely assimilated during rapid bacterial proliferation. Previously, an increase in mineral profile after fermentation is often observed [51,52], although a decrease in mineral content after LAB fermentation was also reported [53]. LC-MS metabolite analysis revealed decomposition of larger PK metabolites, indicating that LF may have induced biotransformation of PK metabolites into simpler and newer low molecular weight metabolites, which ultimately increased the apparent TPC and TFC, and therefore TAC. This is of particular significance since PK metabolites, especially the bioactive iridoid glycosides, have poor oral bioavailability, which limits their translational potential[54]. To circumvent this, enhancing the bioavailability of polyphenols and other secondary metabolites through fermentation, as used in the present study, is considered a viable approach, which may also have contributed to improvements in the bioefficacy of fermented PK [6,39,55].

To conclude, the present study identified that PK can specifically induce LAB growth and that its metabolites can be used as a sole substrate by LF, resulting in a fermented product characterised by enhanced LF proliferation, metabolite biotransformation, and cytoprotective attributes. The identified plant, *Picrorhiza kurroa* (PK), is ethnomedicinally acclaimed, which grows between an altitude of 3000-4500 m and typically reaches a height of 10–20 cm. It bears basal alternate leaves measuring 5–15 cm in length, a rhizome extending 2.5–12.0 cm, and greyish-brown roots [56]. The plant is often consumed as a powder mixed with vehicles like warm water or honey as a decoction [57]; however, the plant extract can also be developed as a functional food or nutraceutical in specific food matrices, such as milk[58]. Overall, we identify a novel synbiotic PK-LF combination and demonstrate that underutilised Himalayan plants can serve as valuable resources for designing novel synbiotics. Future studies should be aimed at developing the PK-LF combination either as a synergistic synbiotic (using dried components) or as a fermented functional ingredient through fortification of food matrices.

## List of Abbreviations

AV: Adhatoda vasica
E. coli: Escherichia coli
LAB: lactic acid bacteria
LC: Lactobacillus casei
LF: Lactobacillus fermentum
LR: Lactobacillus rhamnosus GG
PAS: prebiotic activity score
OD: optical density
PCA: principal component analysis
PK: Picrorhiza kurroa
PK-UF: unfermented Picrorhiza kurroa
PK-F: fermented Picrorhiza kurroa
ROS: reactive oxygen species
TAC: total antioxidant capacity
TFC: total flavonoid content;
TPC: total phenolic content.

## Declarations

### Ethics approval and consent to participate

Not applicable

### Consent for publication

Not applicable.

### Availability of data and material

All data are presented in the manuscript.

### CRediT authorship contribution statement

Amit Kumar: Investigation, Data curation. Bhawna Diwan: Investigation, Data curation. Mohd Adil Khan: Investigation, Data curation. Ankita Awasthi: Investigation, Data curation. Ekta Bala: Investigation, Data curation. Parveen Kumar Verma: Methodology, Writing and Reviewing. Rohit Sharma: Conceptualisation, Funding acquisition, Supervision, Methodology, Reviewing, Writing original draft.

## Acknowledgements

Authors acknowledge infrastructure support from Shoolini University, Solan.

