## Supplementary Table 1 for "Selective identification of the herb *Picrorhiza kurroa* and probiotic *Lactobacillus fermentum* as a synbiotic with fermentation-enhanced physicochemical, biological, and metabolomic properties"

**Supplementary Table 1.** Identification of characteristic secondary metabolites in *Picrorhiza kurroa* through LC-MS.

| **S. No.** | **Name of Compound** | **RT** | **Actual MW** | **Observed**  **Mass** | **Mass Fragmentation** | **Ref.** |
| --- | --- | --- | --- | --- | --- | --- |
| 1 | Catalpol | 2.52 | 362.3310 | [M + Na] = 385.1900 | 385, 183, 177 | 1 |
| 2 | Picroside I | 12.59 | 492.1632 | [M + Na] = 515.2566 | 515, 293, 177, 131 | 1 |
| 3^*^ | Kutkoside | 6.96 | 512.1530 | [M + Na] = 535.2527 | 535, 333, 151 | 2 |
| 4^*^ | Picroside II | 8.32 | 512.1530 | [M + Na] = 535.2527 | 535, 497, 333, 151 | 1 |
| 5 | Picroside III | 12.70 | 538.1686 | [M + Na] = 561.2748 | 561, 523, 183, 177, 131 | 3 |

* The retention time of identified compounds might interchange due to similar molecular weight.

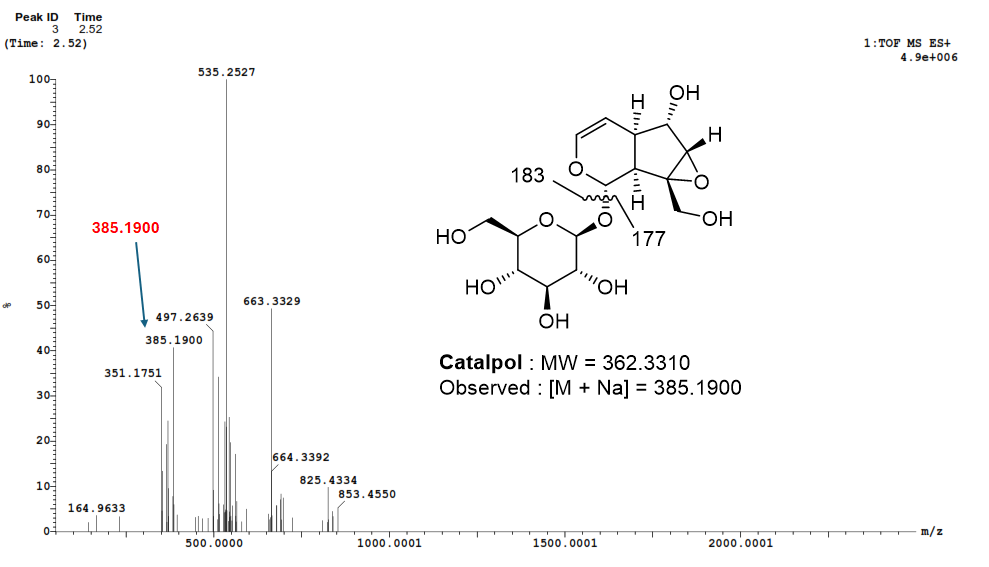
**Identified characteristic metabolites of PK:**

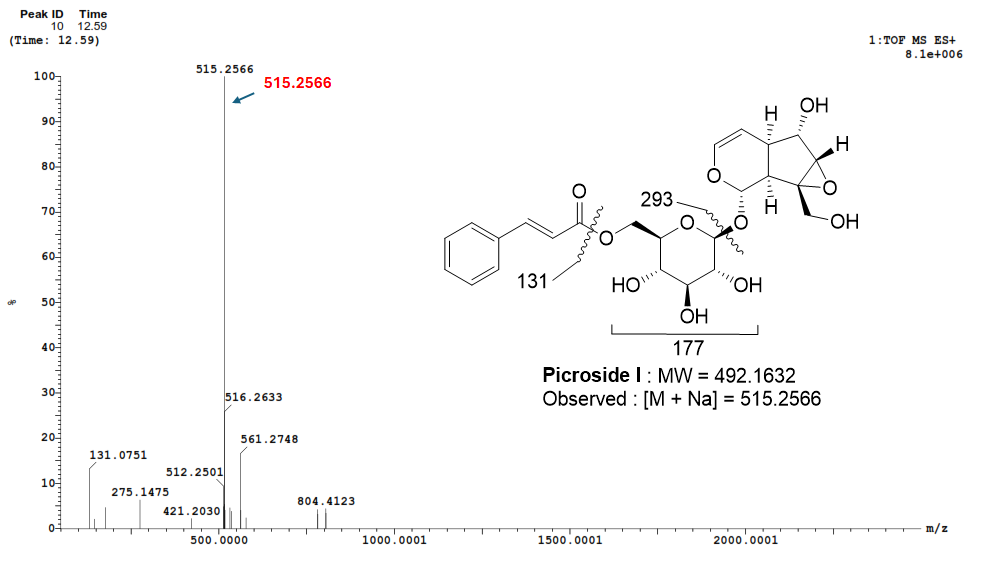

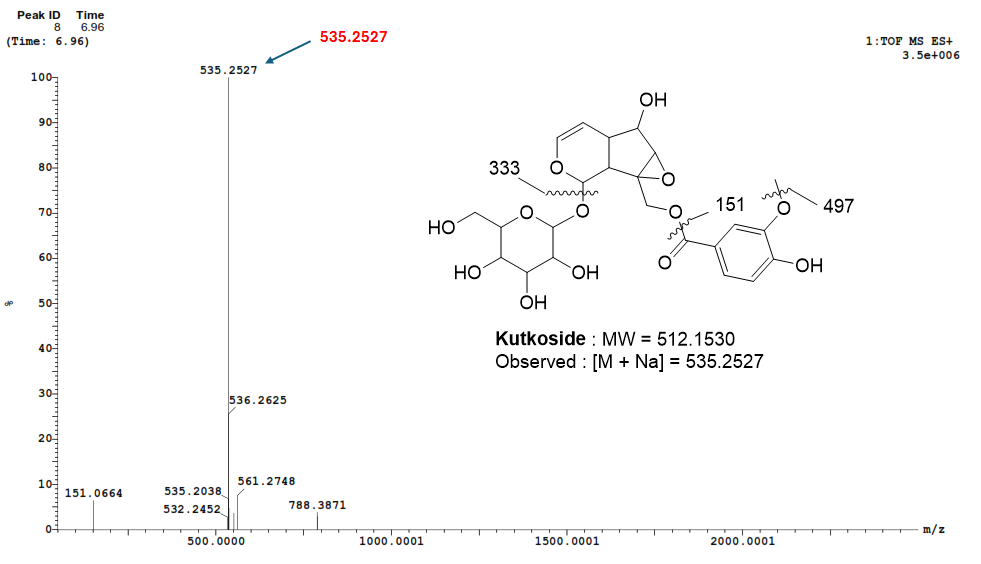

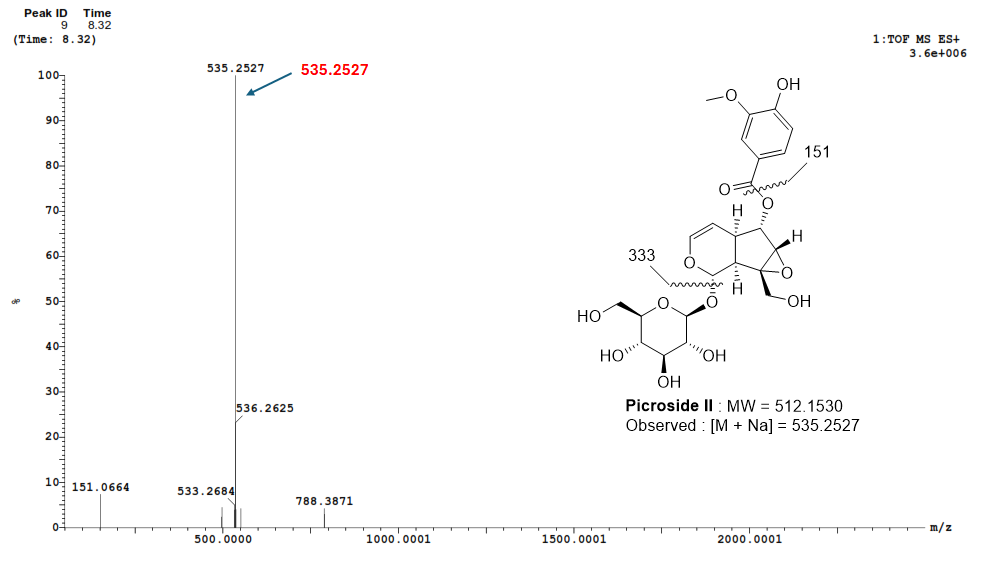

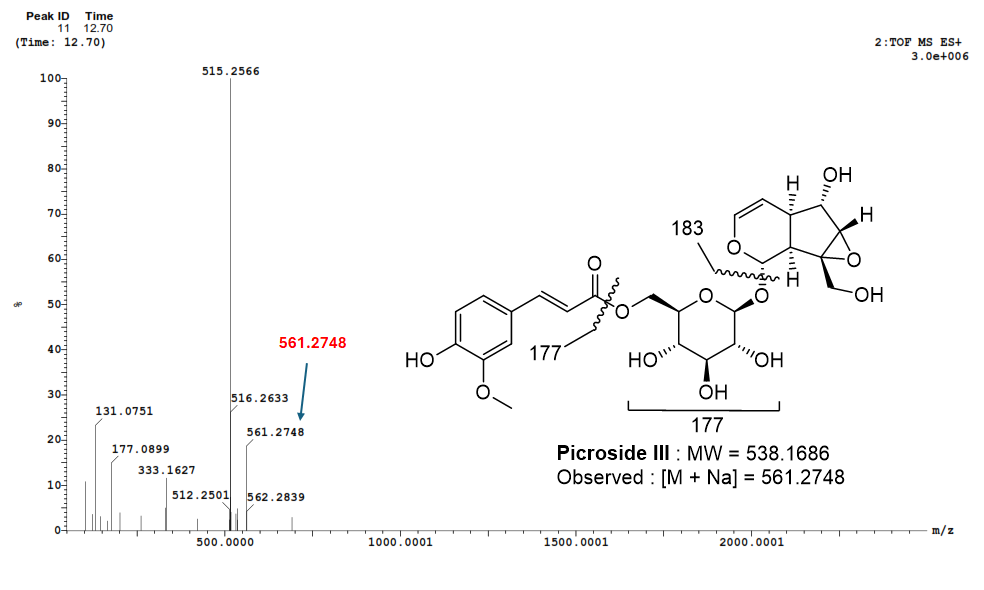
