## Supplementary Figures 1-2 for "Selective identification of the herb *Picrorhiza kurroa* and probiotic *Lactobacillus fermentum* as a synbiotic with fermentation-enhanced physicochemical, biological, and metabolomic properties"

**Openlynx Report SAIF,PANJAB UNIVERSITY CHANDIGARH** 

Sample: 32 Vial:1:C,1 ID:
File:AMIT_SAMPLE_A Date:16-Aug-2025 Time:04:06:06
Description:

Printed: Mon Aug 18 14:20:06 2025

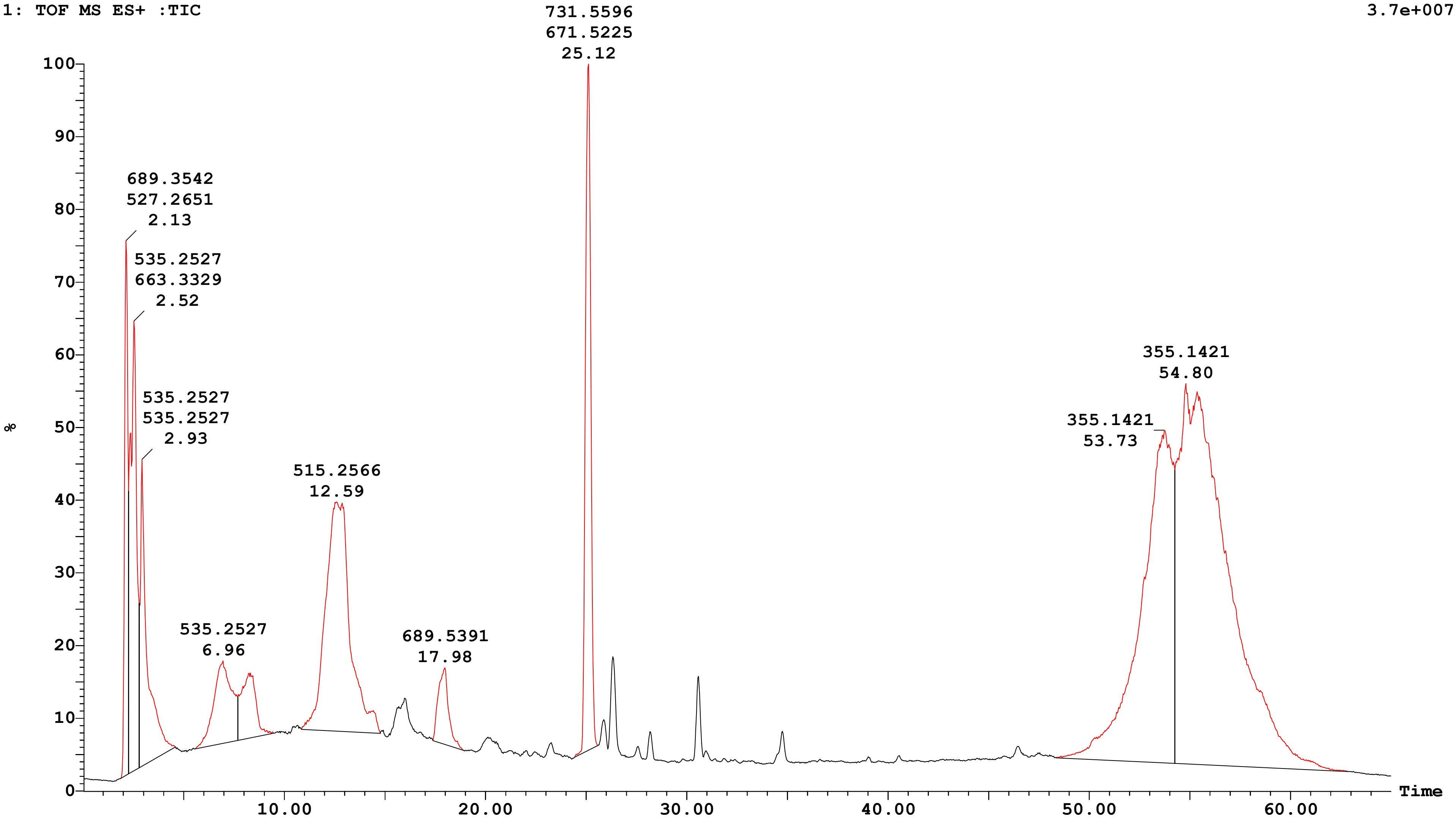


| Peak Number | Compound | Time | AreaAbs | Area %Total | Width | Height | Mass Found |
|---|---|---|---|---|---|---|---|
| 1 | | 2.13 | 5e+006 | 3.62 | 0 | 3e+007 | |
| 3 | | 2.52 | 8e+006 | 5.89 | 1 | 2e+007 | |
| 5 | | 2.93 | 6e+006 | 4.33 | 2 | 2e+007 | |
| 8 | | 6.96 | 4e+006 | 3.05 | 2 | 4e+006 | |
| 9 | | 8.32 | 3e+006 | 1.96 | 2 | 3e+006 | |

Sample: 32 Vial:1:C,1 ID:
File:AMIT_SAMPLE_A Date:16-Aug-2025 Time:04:06:06
Description:

Printed: Mon Aug 18 14:20:06 2025

| | | | | | |
|---|---|---|---|---|---|
| 10 | 12.59 | 2e+007 | 11.5 | 4 | 1e+007 |
| 15 | 17.98 | 2e+006 | 1.61 | 1 | 4e+006 |
| 16 | 25.12 | 1e+007 | 7.57 | 1 | 3e+007 |
| 18 | 53.73 | 3e+007 | 21.52 | 6 | 2e+007 |
| 19 | 54.80 | 6e+007 | 38.92 | 9 | 2e+007 |

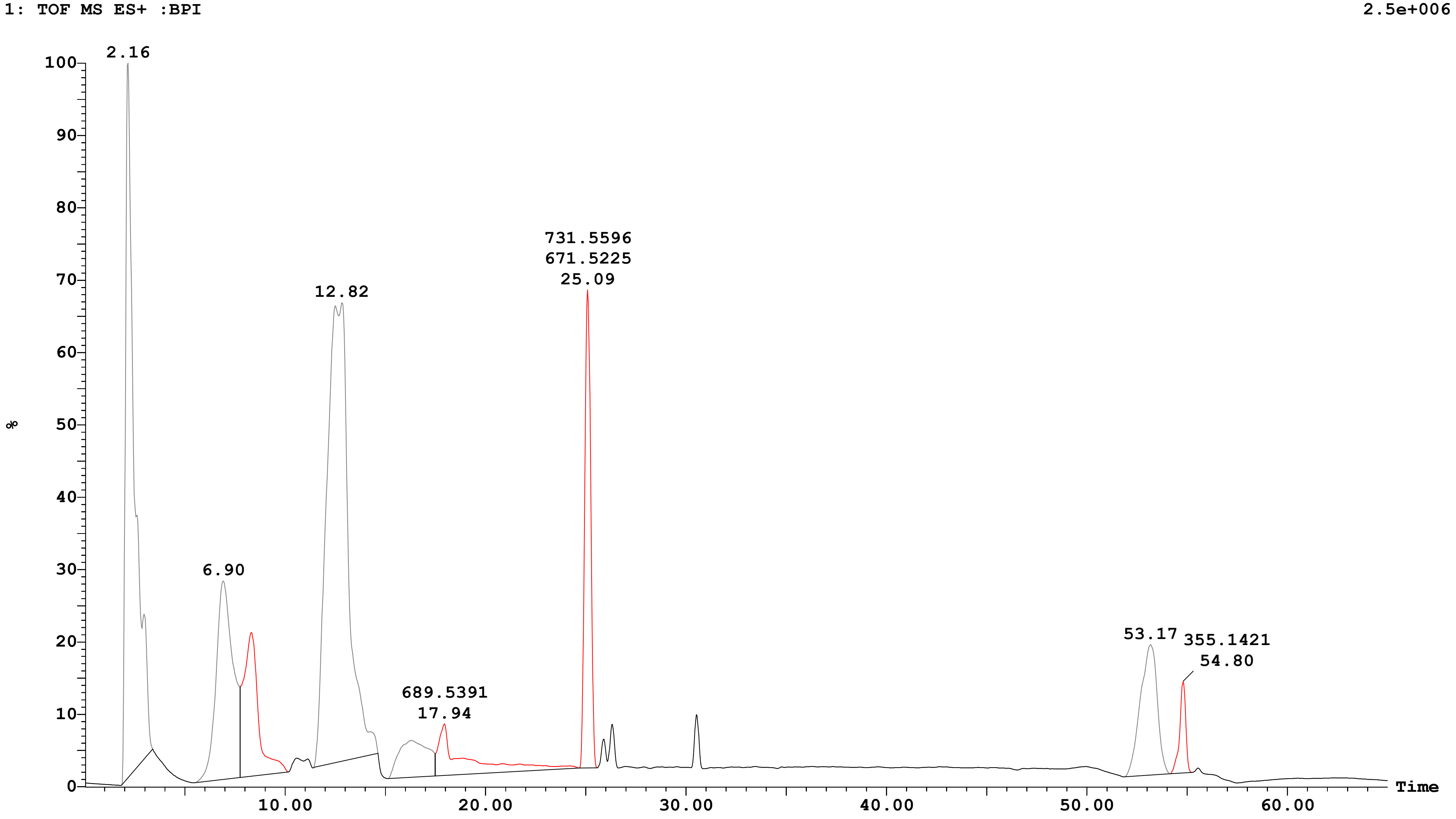

**Openlynx Report SAIF,PANJAB UNIVERSITY CHANDIGARH**

Sample: 32 | Vial:1:C,1 | ID:
File:AMIT_SAMPLE_A | Date:16-Aug-2025 | Time:04:06:06
Description:

Printed: Mon Aug 18 14:20:06 2025

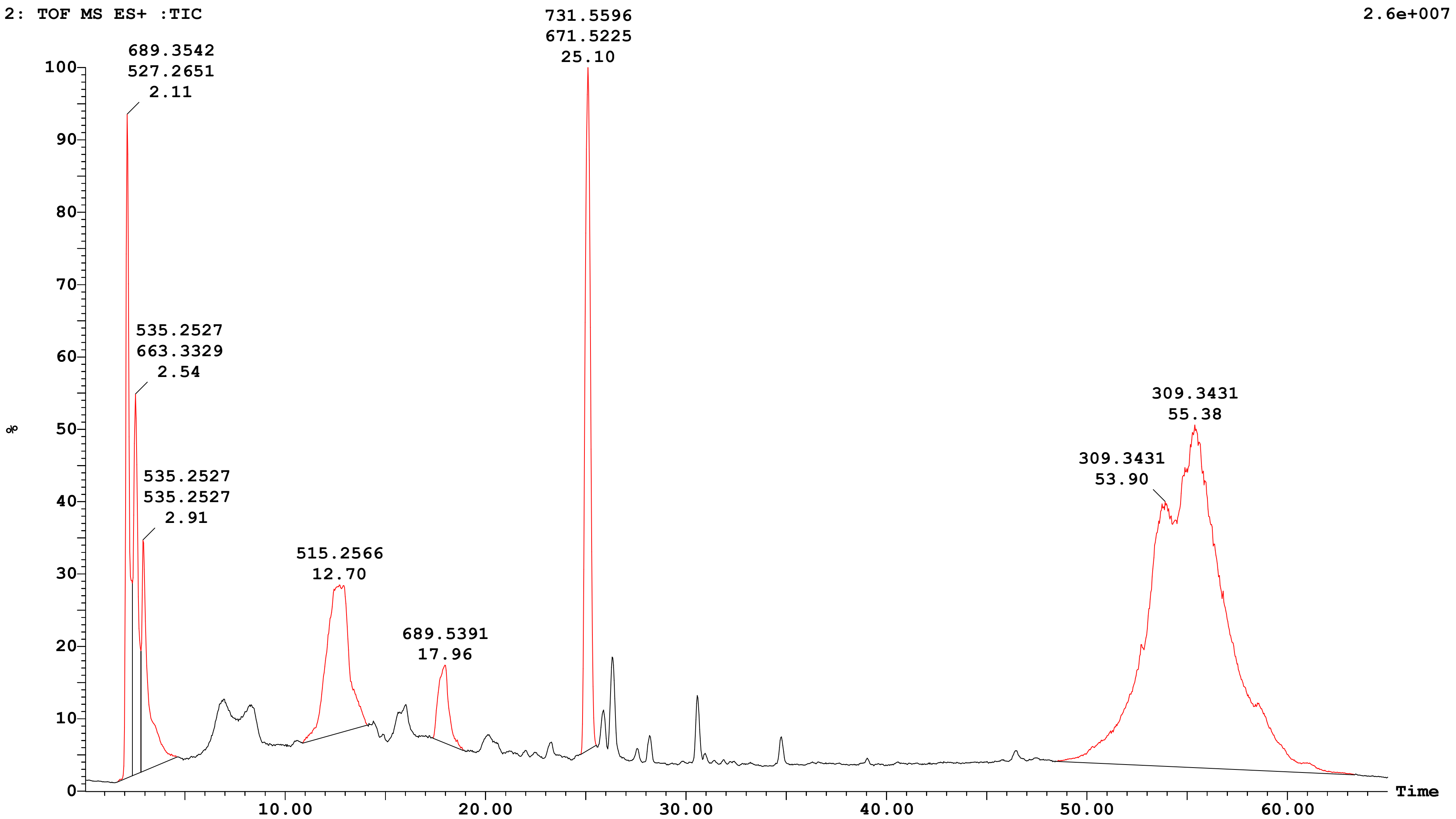


| Peak Number | Compound | Time | AreaAbs | Area %Total | Width | Height | Mass Found |
|---|---|---|---|---|---|---|---|
| 1 | | 2.11 | 5e+006 | 6.25 | 1 | 2e+007 | |
| 3 | | 2.54 | 4e+006 | 4.72 | 0 | 1e+007 | |
| 5 | | 2.91 | 3e+006 | 4.09 | 2 | 8e+006 | |
| 11 | | 12.70 | 7e+006 | 9.05 | 3 | 5e+006 | |
| 14 | | 17.96 | 2e+006 | 2.17 | 2 | 3e+006 | |

Sample: 32 Vial:1:C,1 ID:
File:AMIT_SAMPLE_A Date:16-Aug-2025 Time:04:06:06
Description:

Printed: Mon Aug 18 14:20:06 2025

| | | | | | |
|---|---|---|---|---|---|
| 16 | 25.10 | 8e+006 | 9.66 | 1 | 2e+007 |
| 20 | 55.38 | 5e+007 | 64.06 | 15 | 1e+007 |

2: TOF MS ES+ :BPI 1.3e+006

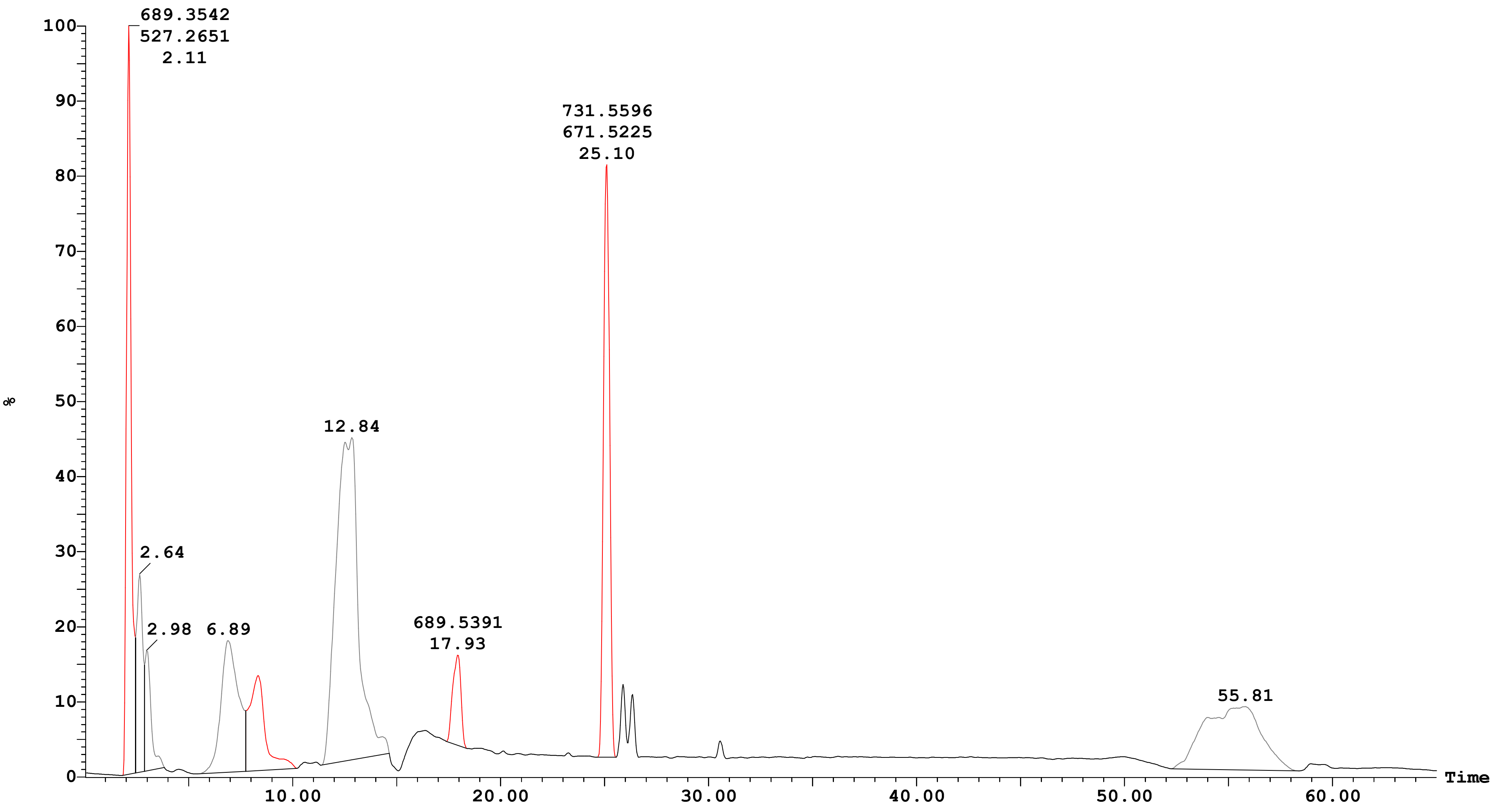

**Openlynx Report SAIF,PANJAB UNIVERSITY CHANDIGARH**

Sample: 32 | Vial:1:C,1 | ID:
File:AMIT_SAMPLE_A | Date:16-Aug-2025 | Time:04:06:06
Description:

Printed: Mon Aug 18 14:20:06 2025

| Peak ID | Time |
|---|---|
| 1 | 2.13 |

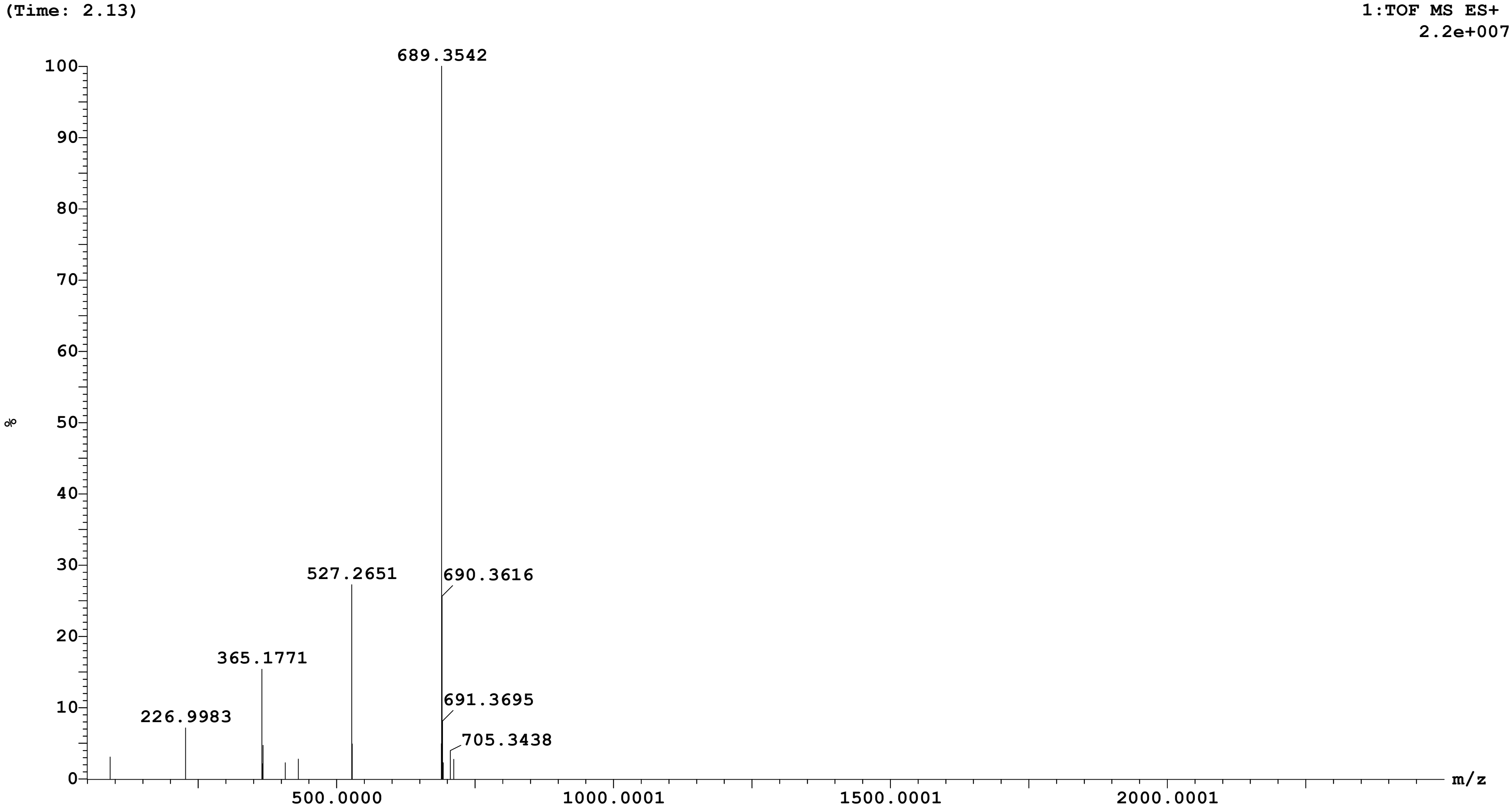

Sample: 32 Vial:1:C,1 ID:
File:AMIT_SAMPLE_A Date:16-Aug-2025 Time:04:06:06
Description:

Printed: Mon Aug 18 14:20:06 2025

| **Peak ID** | **Time** |
|---|---|
| 3 | 2.52 |

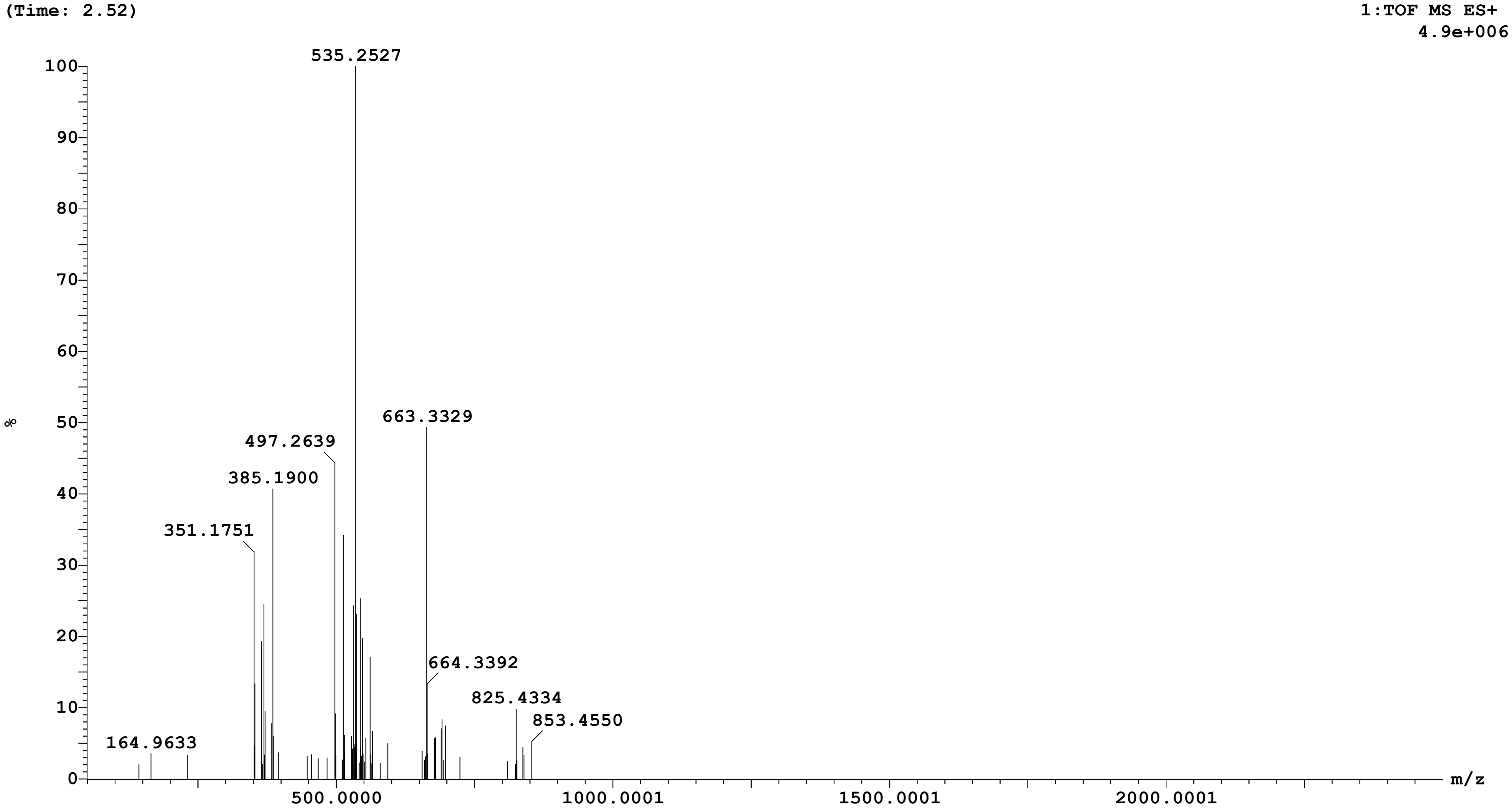

Sample: 32 Vial:1:C,1 ID:
File:AMIT_SAMPLE_A Date:16-Aug-2025 Time:04:06:06
Description:

Printed: Mon Aug 18 14:20:06 2025

| Peak ID | Time |
| --- | --- |
| 5 | 2.93 |

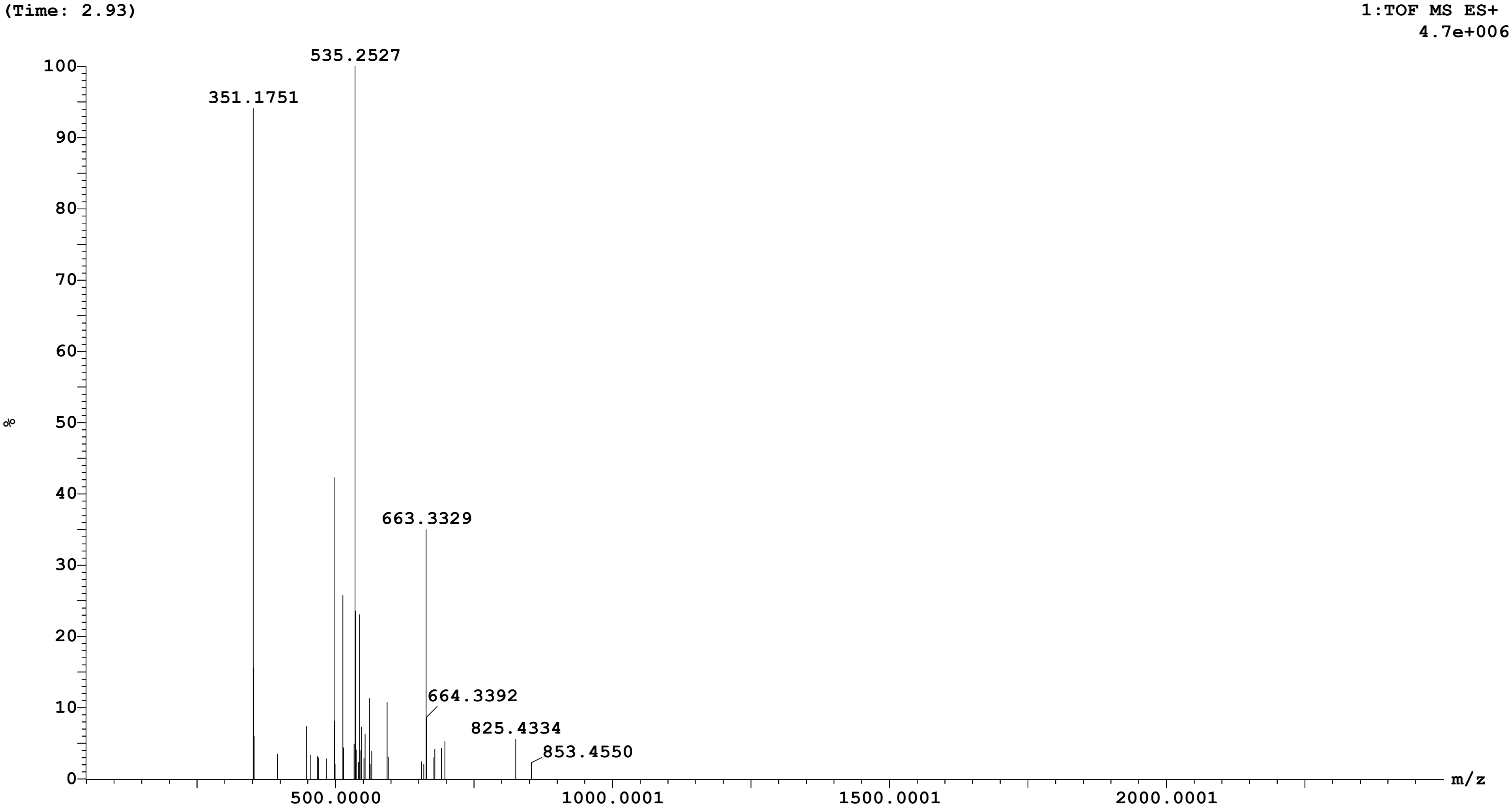

Sample: 32 Vial:1:C,1 ID:
File:AMIT_SAMPLE_A Date:16-Aug-2025 Time:04:06:06
Description:

Printed: Mon Aug 18 14:20:06 2025

| Peak ID | Time |
|---|---|
| 8 | 6.96 |

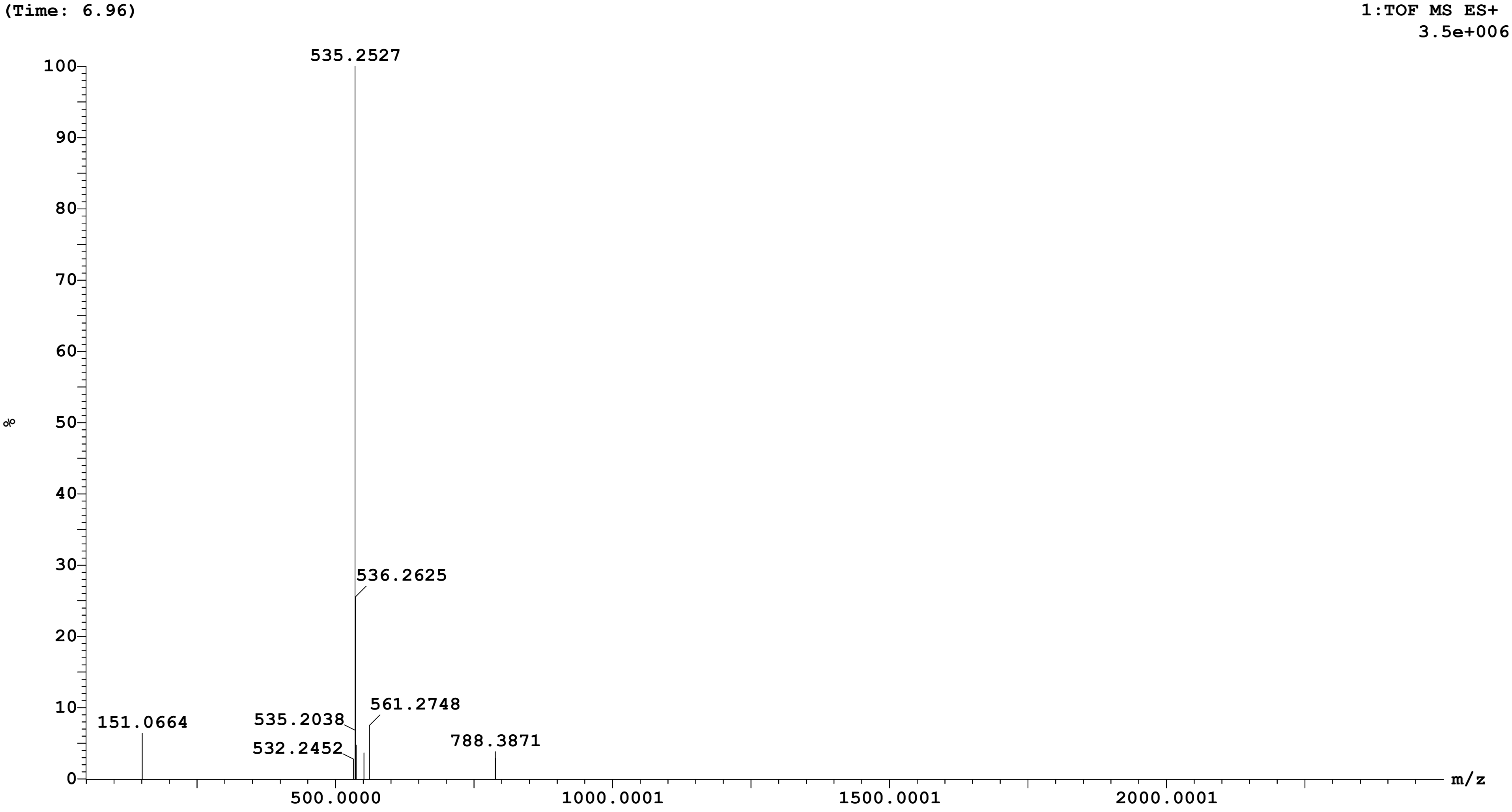

Sample: 32 Vial:1:C,1 ID:
File:AMIT_SAMPLE_A Date:16-Aug-2025 Time:04:06:06
Description:

Printed: Mon Aug 18 14:20:06 2025

| Peak ID | Time |
|---|---|
| 9 | 8.32 |

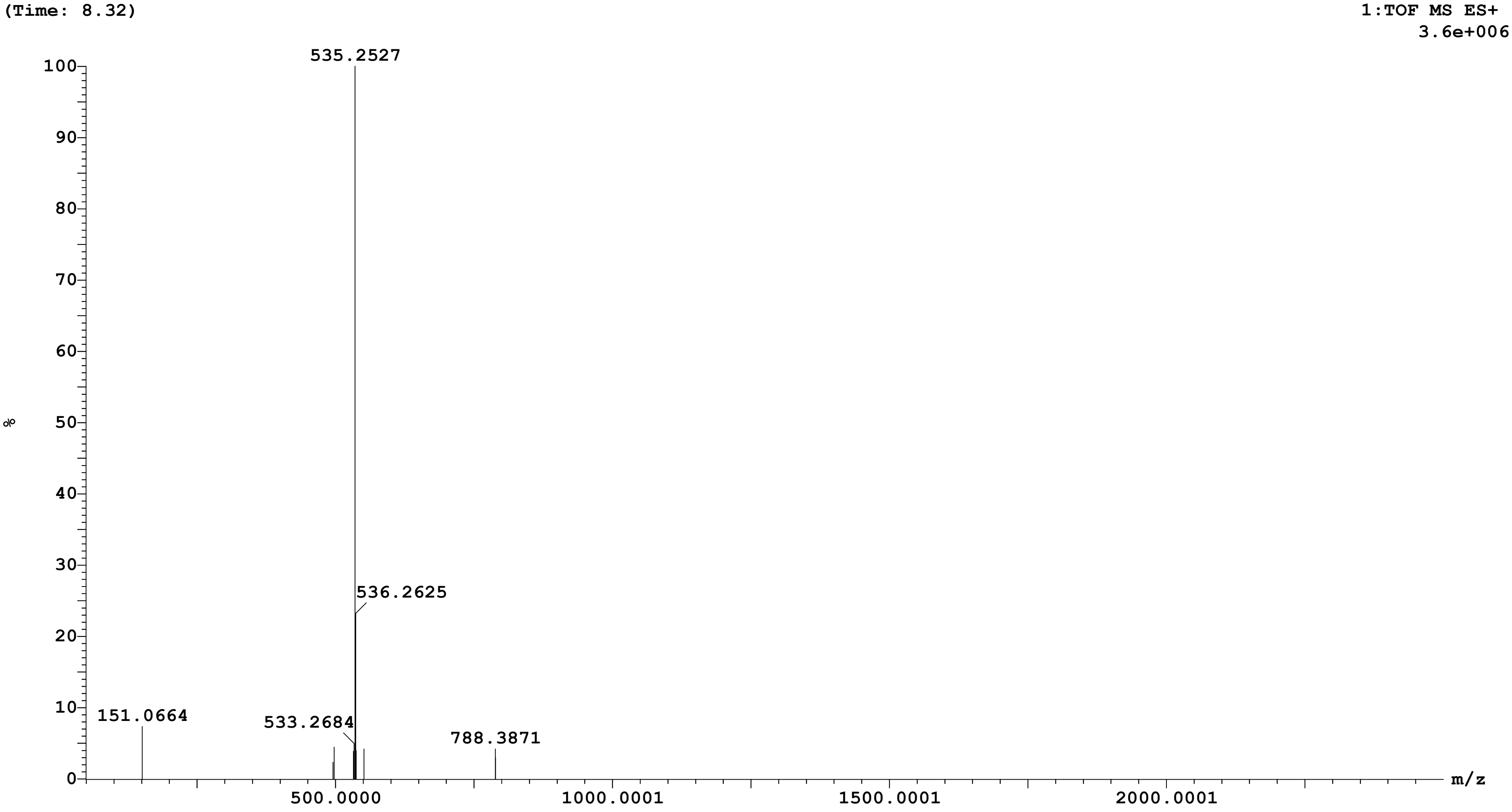

**Openlynx Report SAIF,PANJAB UNIVERSITY CHANDIGARH**

Sample: 32 | Vial:1:C,1 | ID:
File:AMIT_SAMPLE_A | Date:16-Aug-2025 | Time:04:06:06
Description:

Printed: Mon Aug 18 14:20:06 2025

| Peak ID | Time |
|---|---|
| 10 | 12.59 |

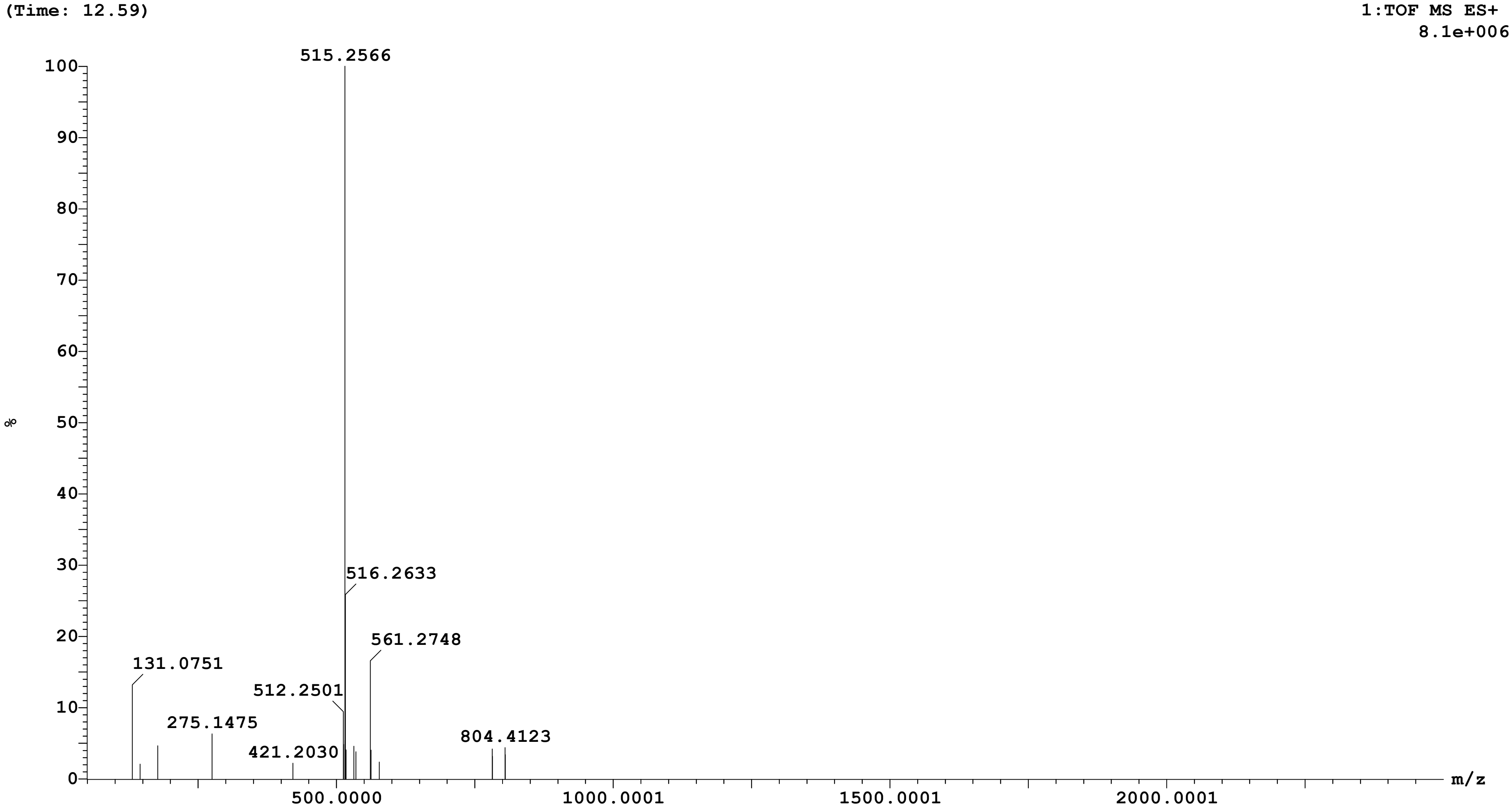

Sample: 32 Vial:1:C,1 ID:
File:AMIT_SAMPLE_A Date:16-Aug-2025 Time:04:06:06
Description:

Printed: Mon Aug 18 14:20:06 2025

| **Peak ID** | **Time** |
|---|---|
| 15 | 17.98 |

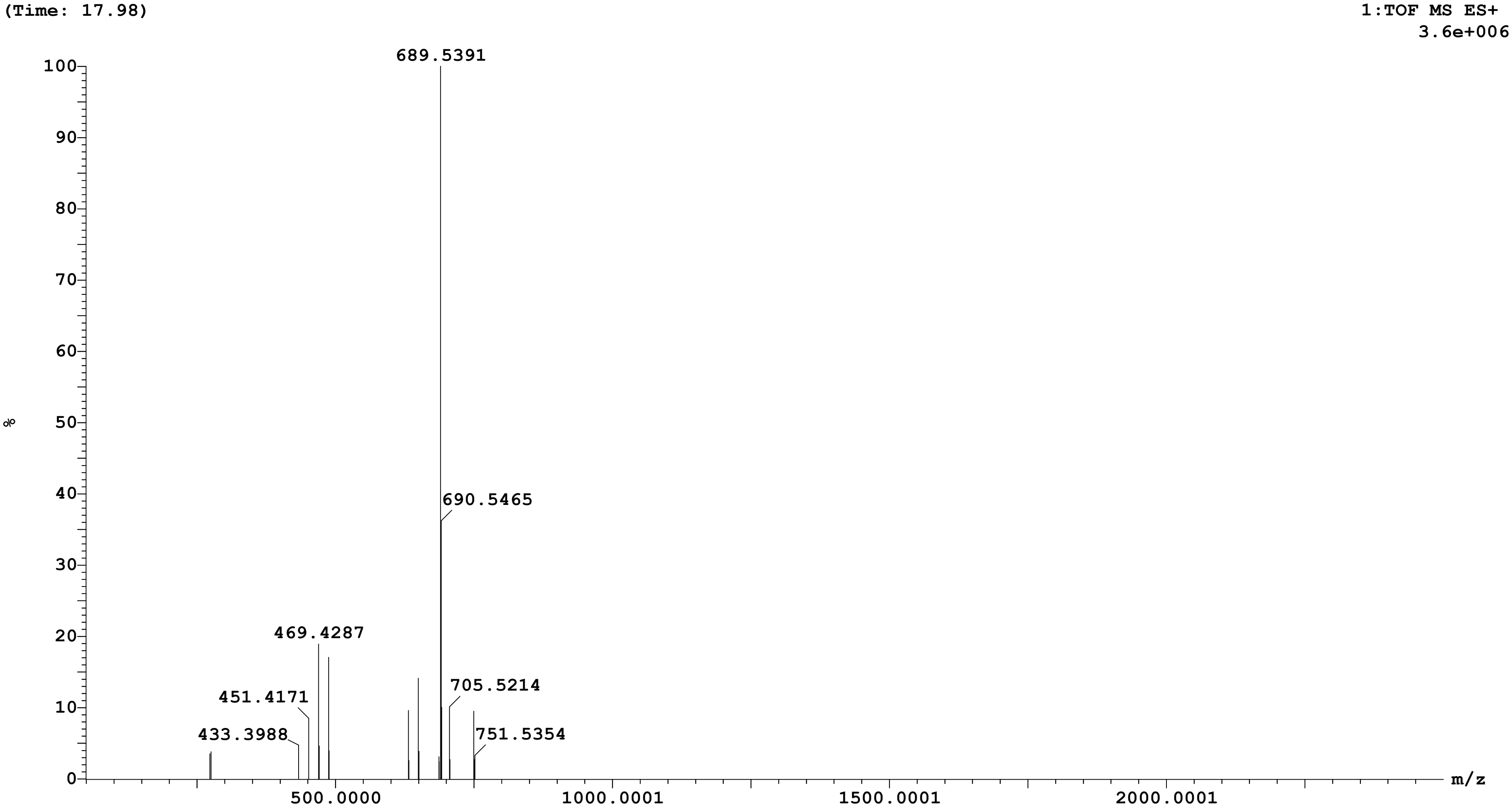

Sample: 32 Vial:1:C,1 ID:
File:AMIT_SAMPLE_A Date:16-Aug-2025 Time:04:06:06
Description:

Printed: Mon Aug 18 14:20:06 2025

| **Peak ID** | **Time** |
|---|---|
| 16 | 25.12 |

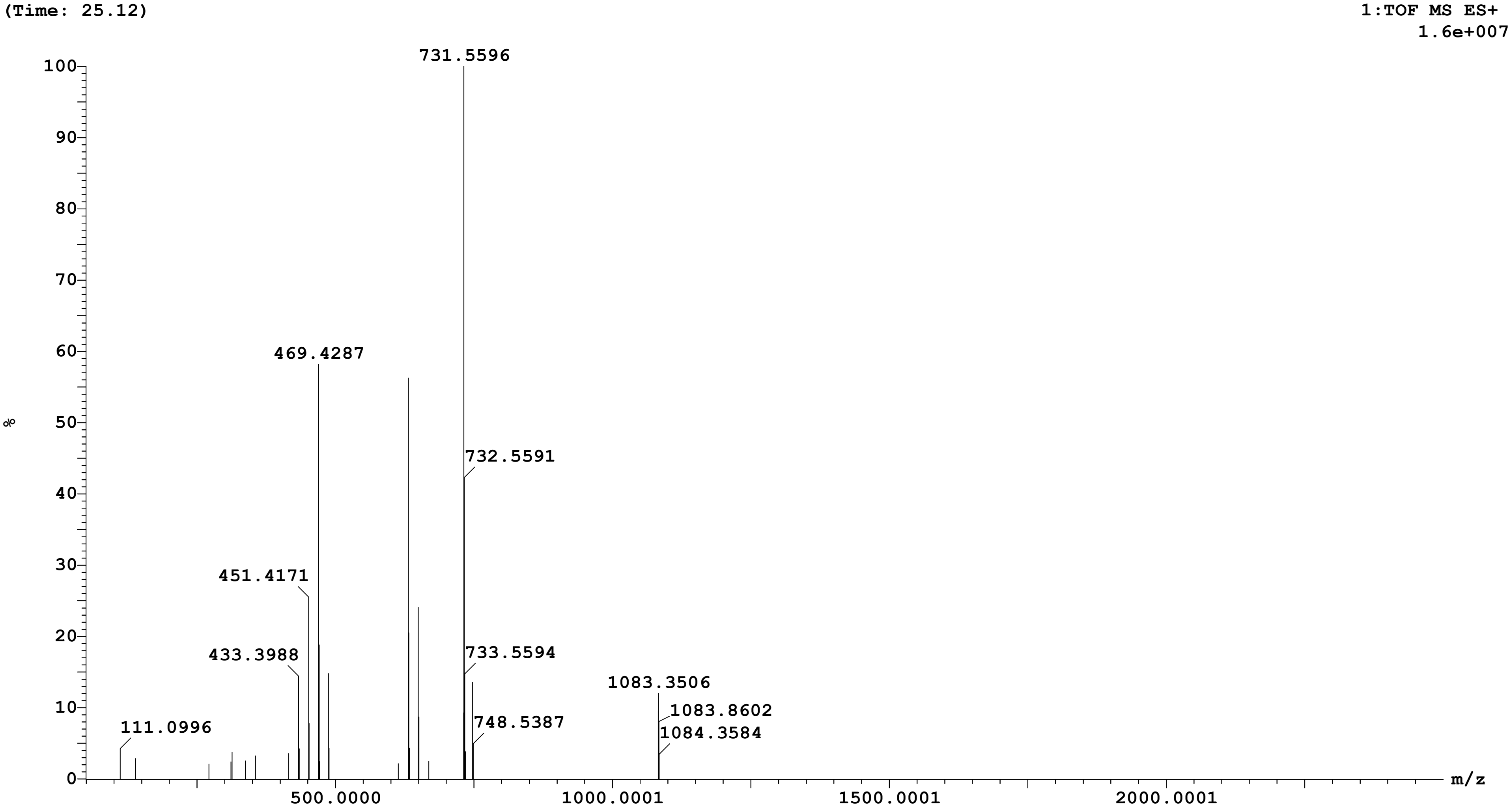

Sample: 32 Vial:1:C,1 ID:
File:AMIT_SAMPLE_A Date:16-Aug-2025 Time:04:06:06
Description:

Printed: Mon Aug 18 14:20:06 2025

| Peak ID | Time |
|---|---|
| 18 | 53.73 |

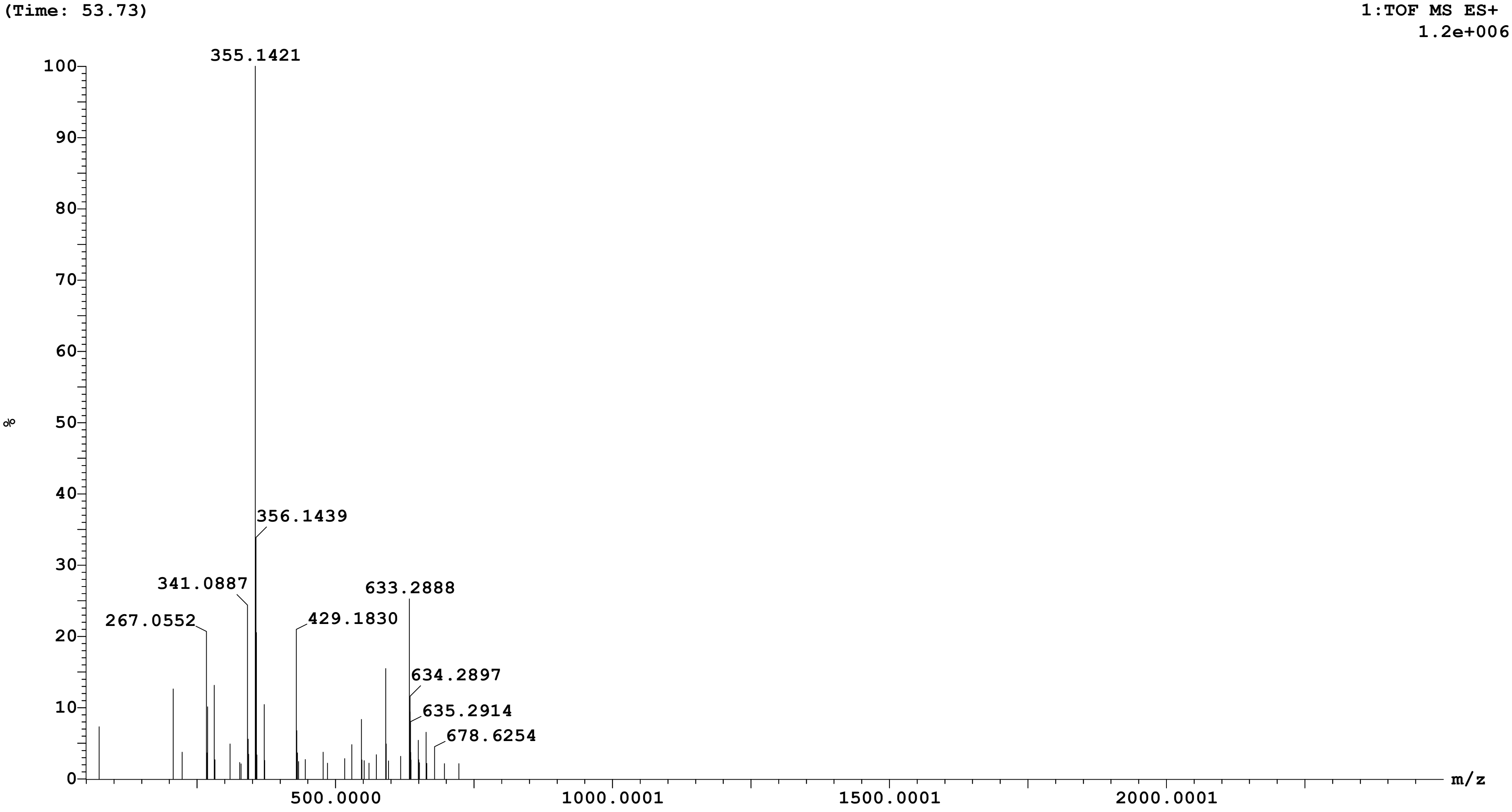

Sample: 32 | Vial:1:C,1 | ID:
File:AMIT_SAMPLE_A | Date:16-Aug-2025 | Time:04:06:06
Description:

Printed: Mon Aug 18 14:20:06 2025

| Peak ID | Time |
|---|---|
| 19 | 54.80 |

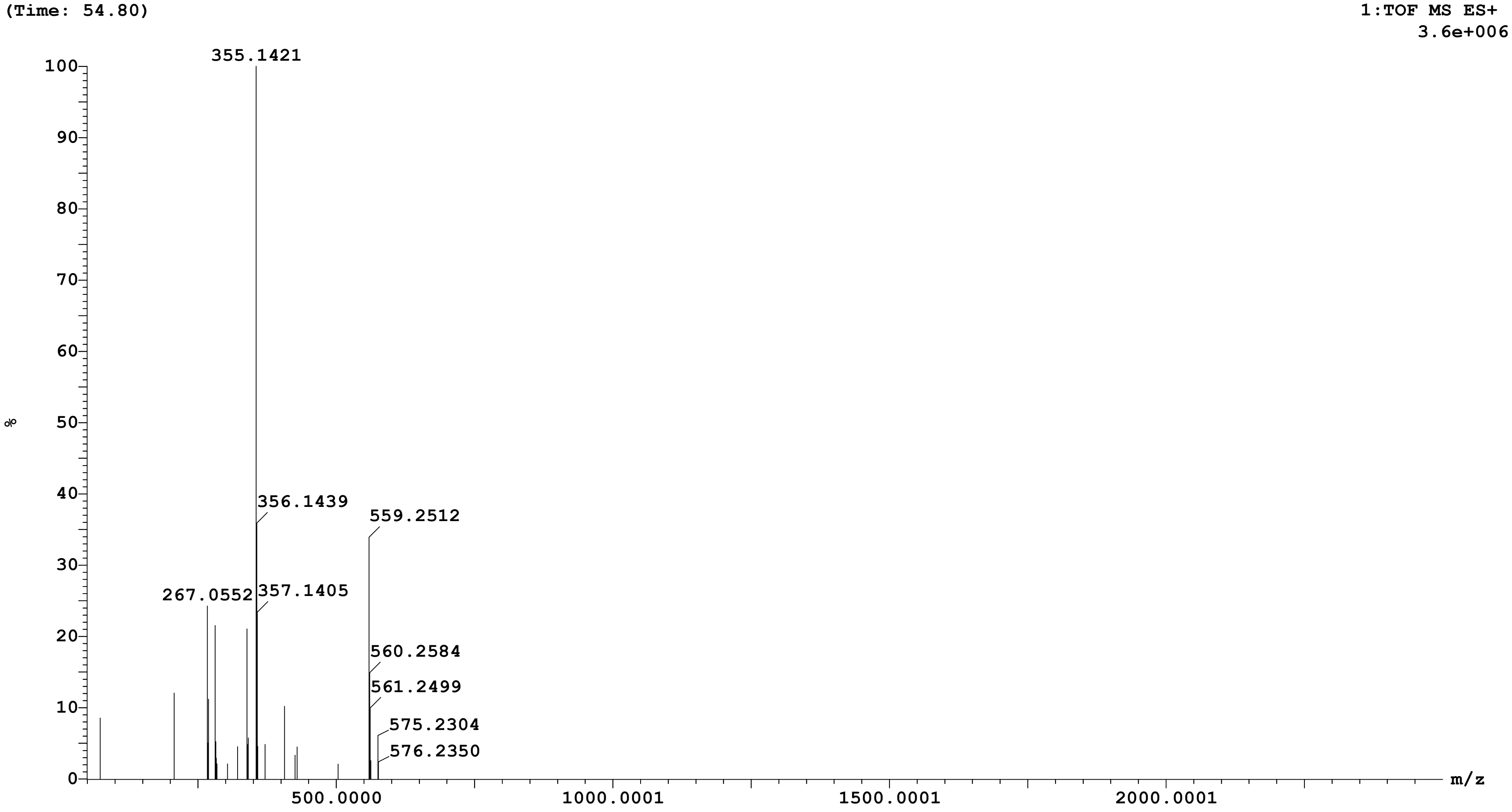

Sample: 32 Vial:1:C,1 ID:
File:AMIT_SAMPLE_A Date:16-Aug-2025 Time:04:06:06
Description:

Printed: Mon Aug 18 14:20:06 2025

| Peak ID | Time |
|---|---|
| 1 | 2.13 |

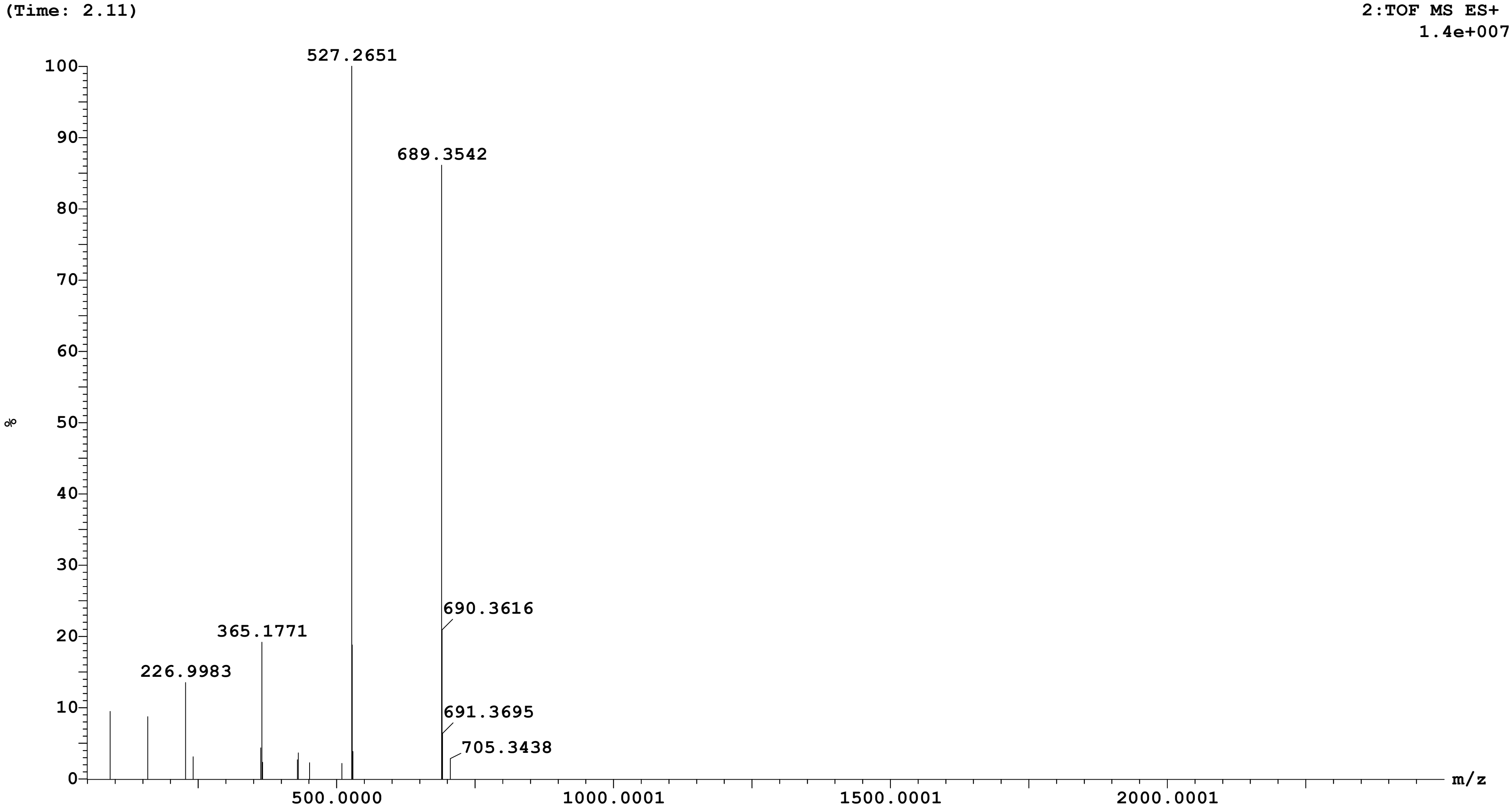

Sample: 32 Vial:1:C,1 ID:
File:AMIT_SAMPLE_A Date:16-Aug-2025 Time:04:06:06
Description:

Printed: Mon Aug 18 14:20:06 2025

| Peak ID | Time |
|---|---|
| 3 | 2.52 |

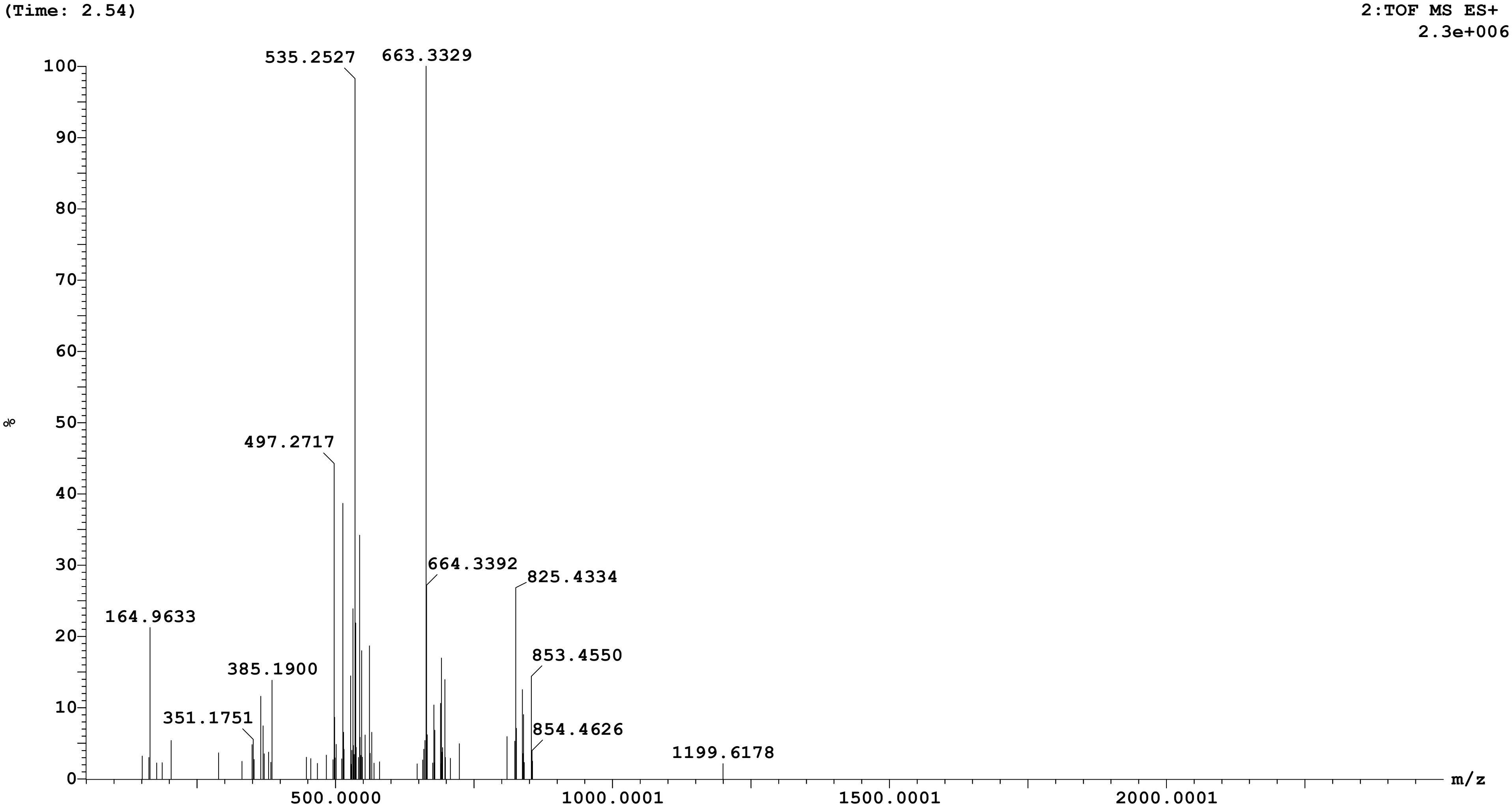

Sample: 32 Vial:1:C,1 ID:
File:AMIT_SAMPLE_A Date:16-Aug-2025 Time:04:06:06
Description:

Printed: Mon Aug 18 14:20:06 2025

| Peak ID | Time |
| --- | --- |
| 5 | 2.93 |

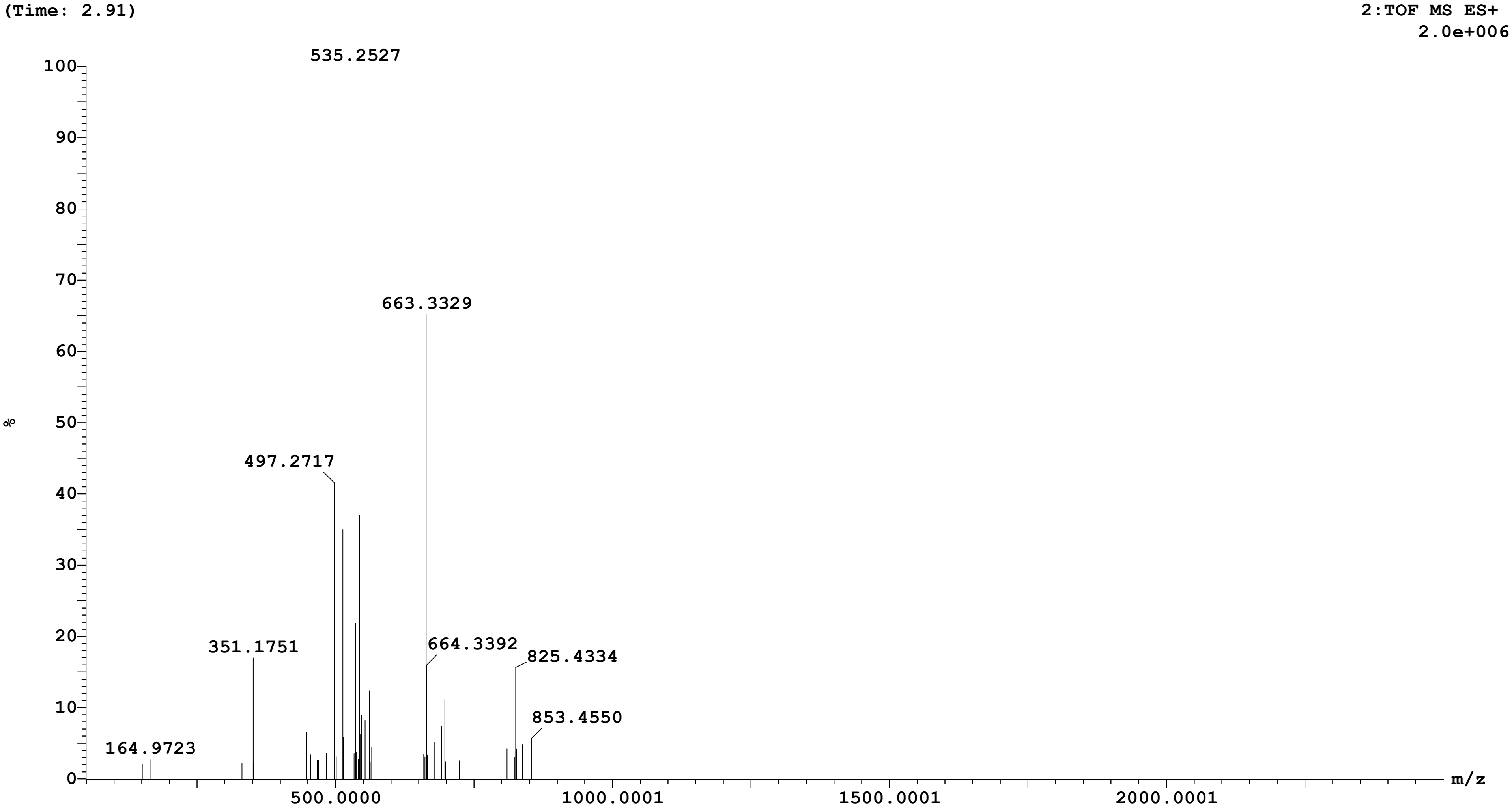

Sample: 32 Vial:1:C,1 ID:
File:AMIT_SAMPLE_A Date:16-Aug-2025 Time:04:06:06
Description:

Printed: Mon Aug 18 14:20:06 2025

| **Peak ID** | **Time** |
|---|---|
| 11 | 12.70 |

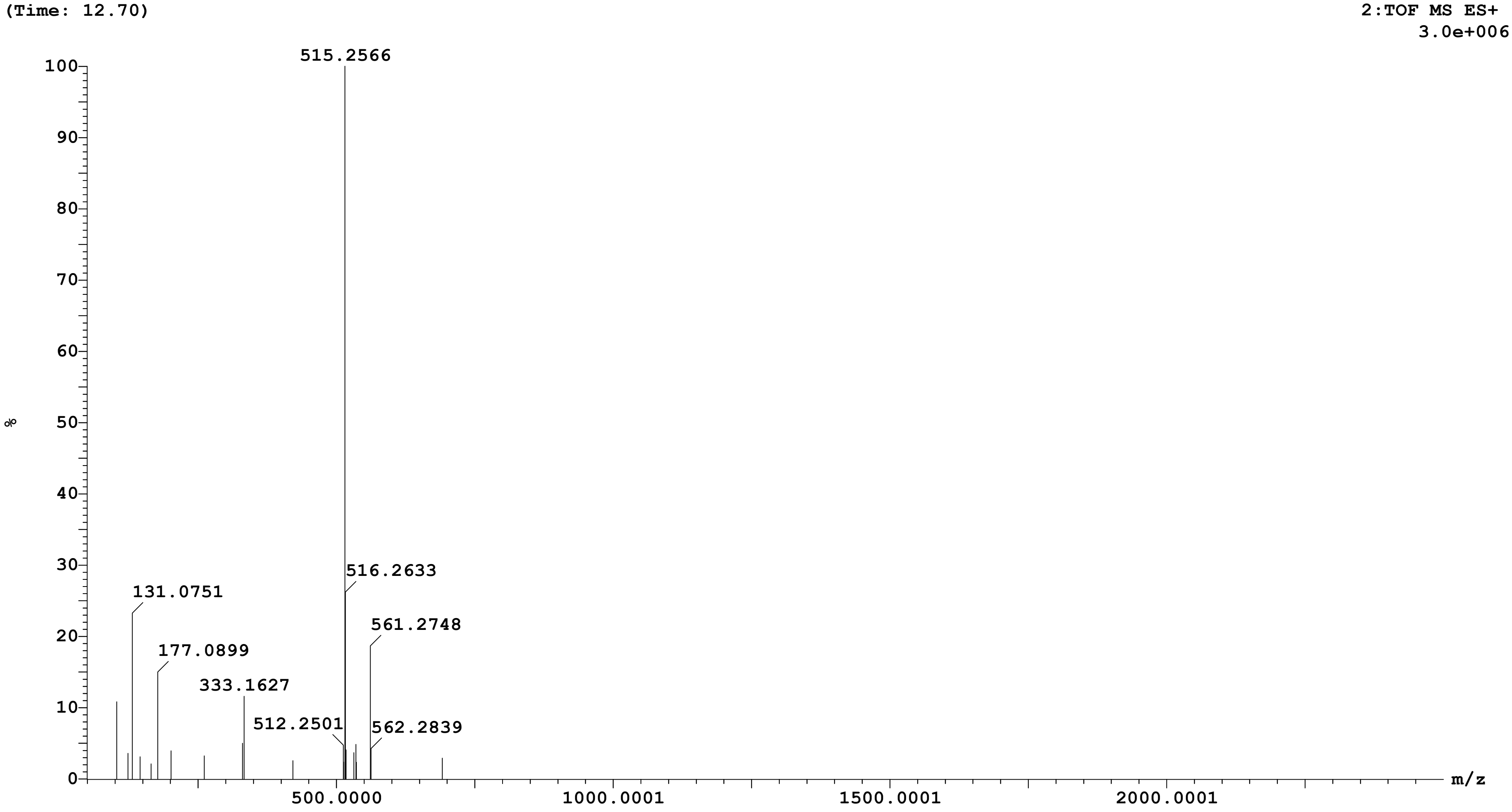

Sample: 32 Vial:1:C,1 ID:
File:AMIT_SAMPLE_A Date:16-Aug-2025 Time:04:06:06
Description:

Printed: Mon Aug 18 14:20:06 2025

| Peak ID | Time |
|---|---|
| 14 | 17.94 |

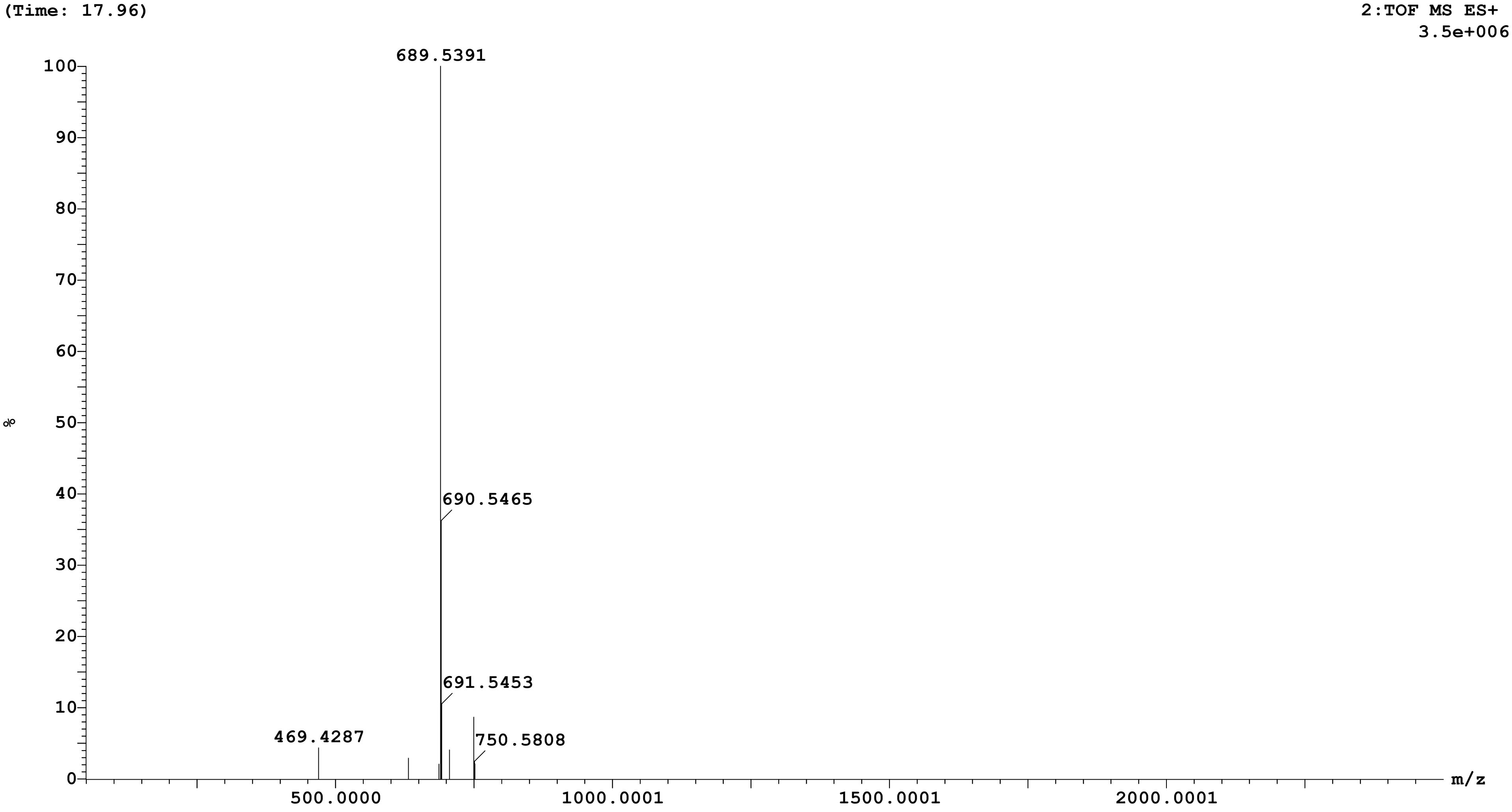

Sample: 32 Vial:1:C,1 ID:
File:AMIT_SAMPLE_A Date:16-Aug-2025 Time:04:06:06
Description:

Printed: Mon Aug 18 14:20:06 2025

| Peak ID | Time |
|---|---|
| 16 | 25.12 |

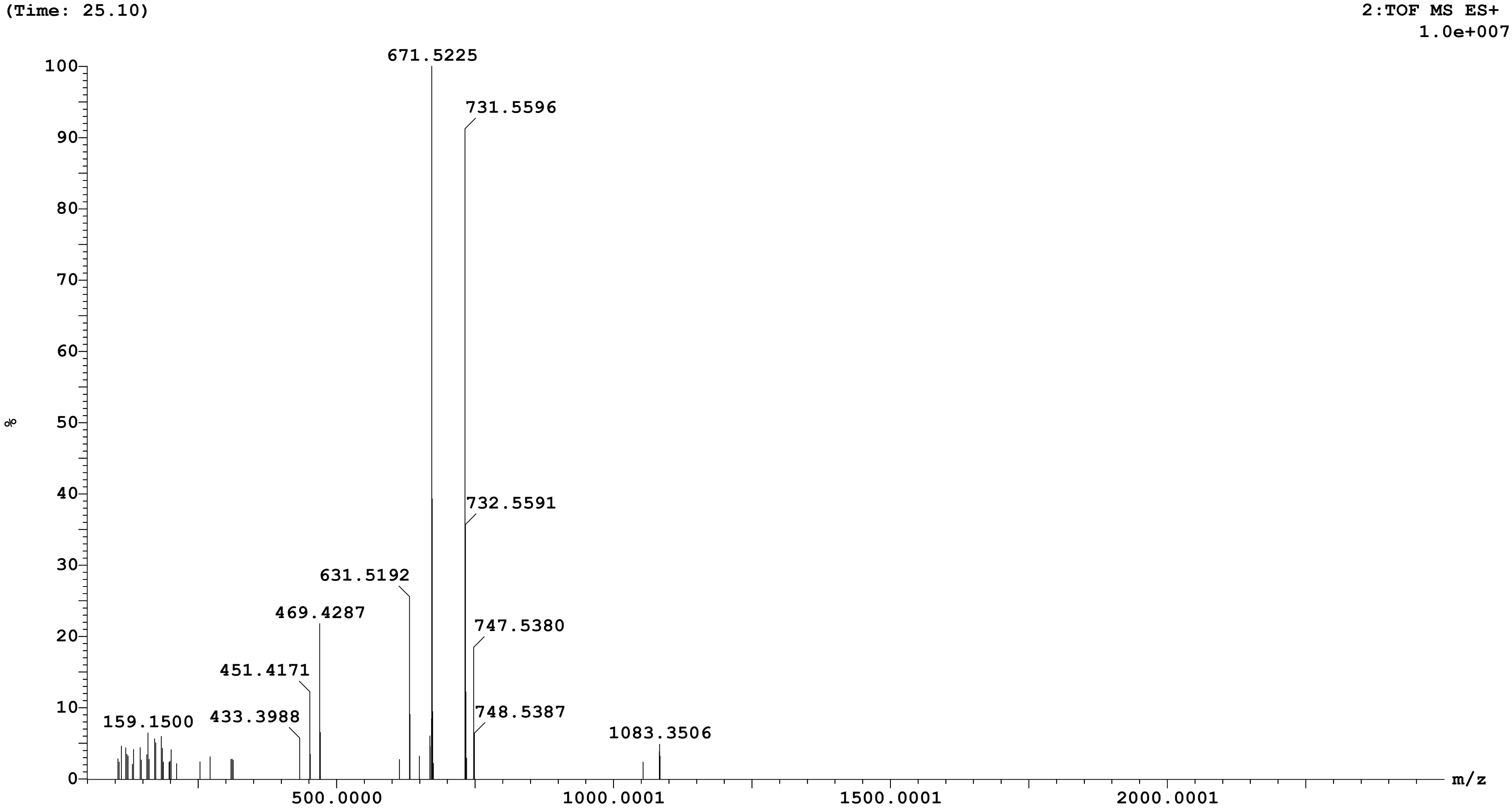

Sample: 32 Vial:1:C,1 ID:
File:AMIT_SAMPLE_A Date:16-Aug-2025 Time:04:06:06
Description:

Printed: Mon Aug 18 14:20:06 2025

| **Peak ID** | **Time** |
|---|---|
| 20 | 55.38 |

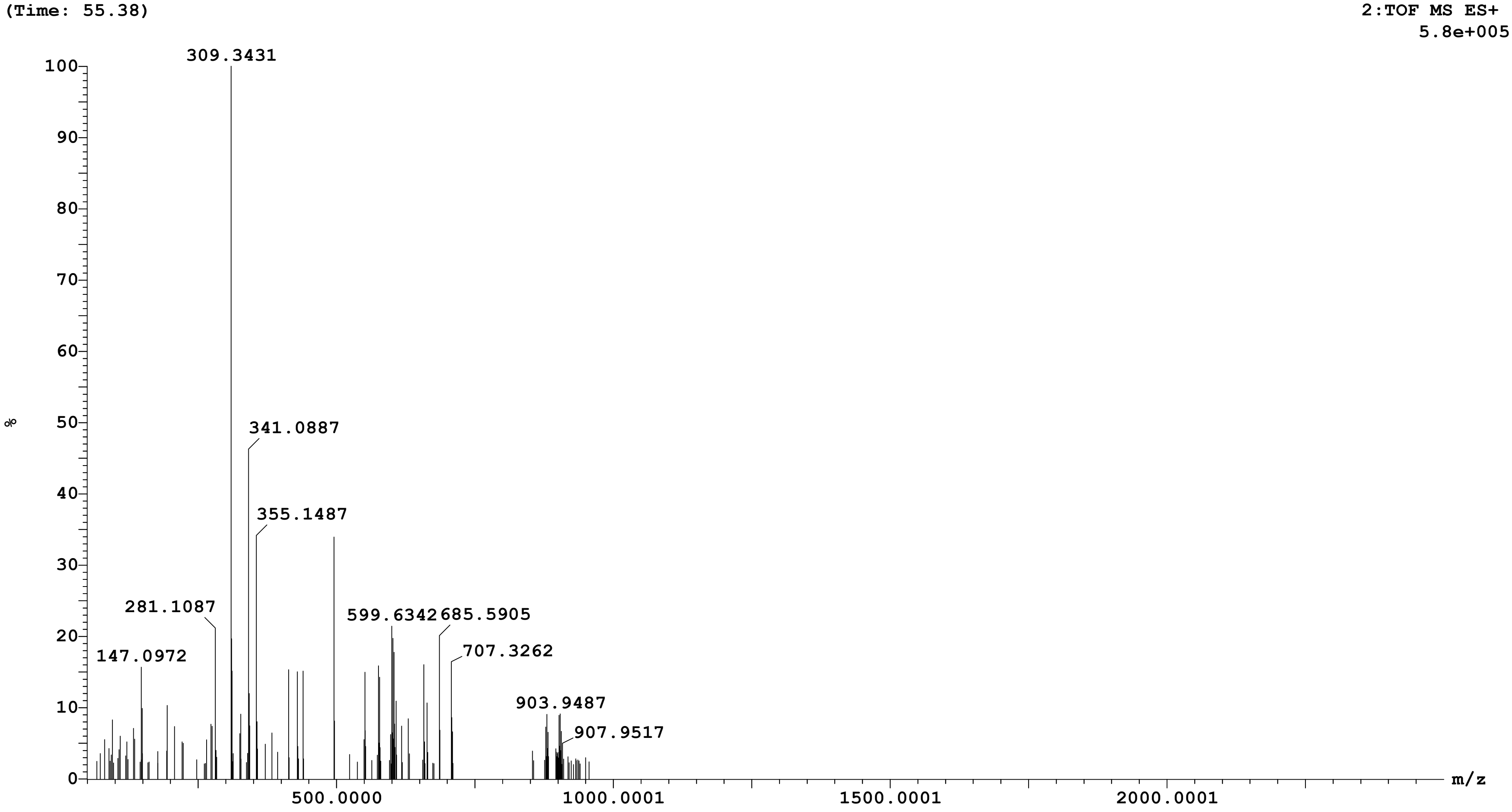
