## Supplementary Figures 1-2 for "Selective identification of the herb *Picrorhiza kurroa* and probiotic *Lactobacillus fermentum* as a synbiotic with fermentation-enhanced physicochemical, biological, and metabolomic properties"

**Openlynx Report SAIF,PANJAB UNIVERSITY CHANDIGARH**



Sample: 34
Vial:1:C,2
ID:
File:AMIT_SAMPLE_B
Date:16-Aug-2025
Time:06:18:36
Description:

Printed: Mon Aug 18 14:21:03 2025

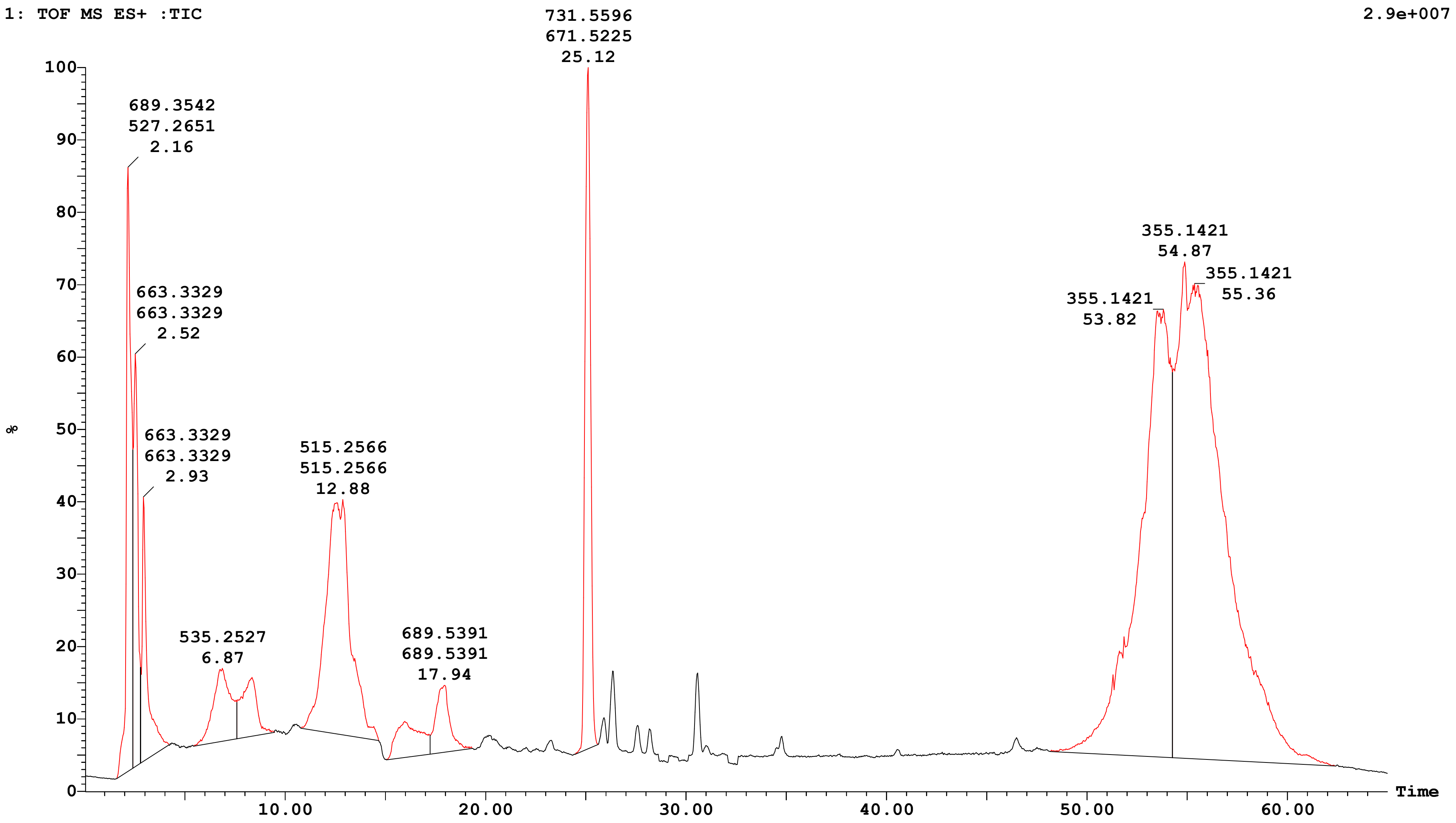


| Peak Number | Compound | Time | AreaAbs | Area %Total | Width | Height | Mass Found |
|---|---|---|---|---|---|---|---|
| 1 | | 2.16 | 7e+006 | 5.04 | 1 | 2e+007 | |
| 2 | | 2.52 | 4e+006 | 3.31 | 0 | 2e+007 | |
| 3 | | 2.93 | 3e+006 | 2.42 | 1 | 1e+007 | |
| 5 | | 6.87 | 3e+006 | 2.24 | 2 | 3e+006 | |
| 6 | | 8.35 | 2e+006 | 1.59 | 2 | 2e+006 | |

Sample: 34 Vial:1:C,2 ID:
File:AMIT_SAMPLE_B Date:16-Aug-2025 Time:06:18:36
Description:

Printed: Mon Aug 18 14:21:03 2025

| | | | | | |
|---|---|---|---|---|---|
| 8 | 12.88 | 1e+007 | 10.06 | 4 | 1e+007 |
| 9 | 15.97 | 2e+006 | 1.52 | 2 | 1e+006 |
| 11 | 17.94 | 2e+006 | 1.64 | 2 | 3e+006 |
| 12 | 25.12 | 9e+006 | 6.56 | 1 | 3e+007 |
| 15 | 53.82 | 3e+007 | 24.29 | 6 | 2e+007 |
| 17 | 54.87 | 6e+007 | 41.34 | 8 | 2e+007 |

**Openlynx Report SAIF,PANJAB UNIVERSITY CHANDIGARH**

Sample: 34 Vial:1:C,2 ID:
File:AMIT_SAMPLE_B Date:16-Aug-2025 Time:06:18:36
Description:

Printed: Mon Aug 18 14:21:03 2025

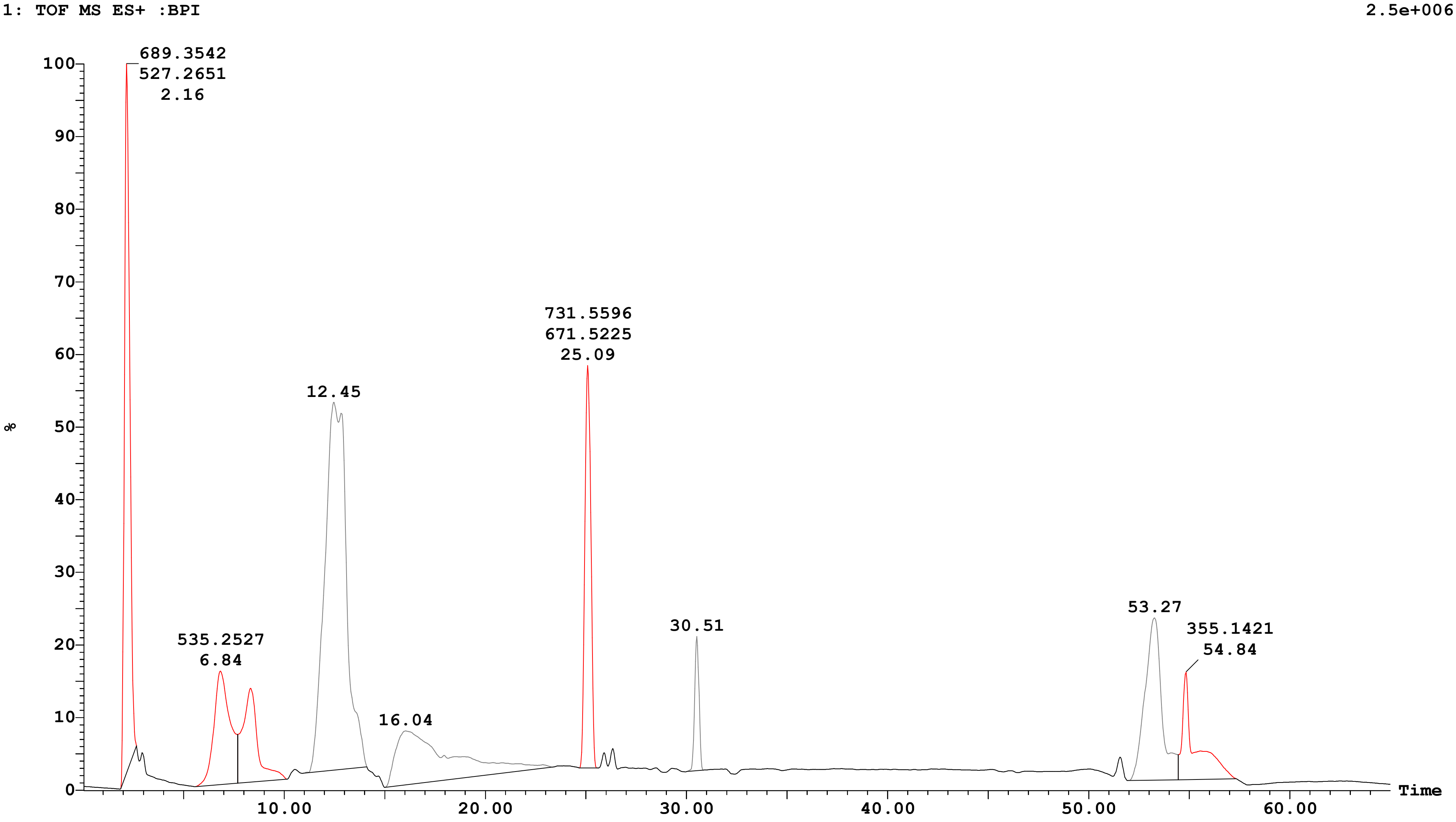

**Openlynx Report SAIF,PANJAB UNIVERSITY CHANDIGARH**

Sample: 34 Vial:1:C,2 ID:
File:AMIT_SAMPLE_B Date:16-Aug-2025 Time:06:18:36
Description:

Printed: Mon Aug 18 14:21:03 2025

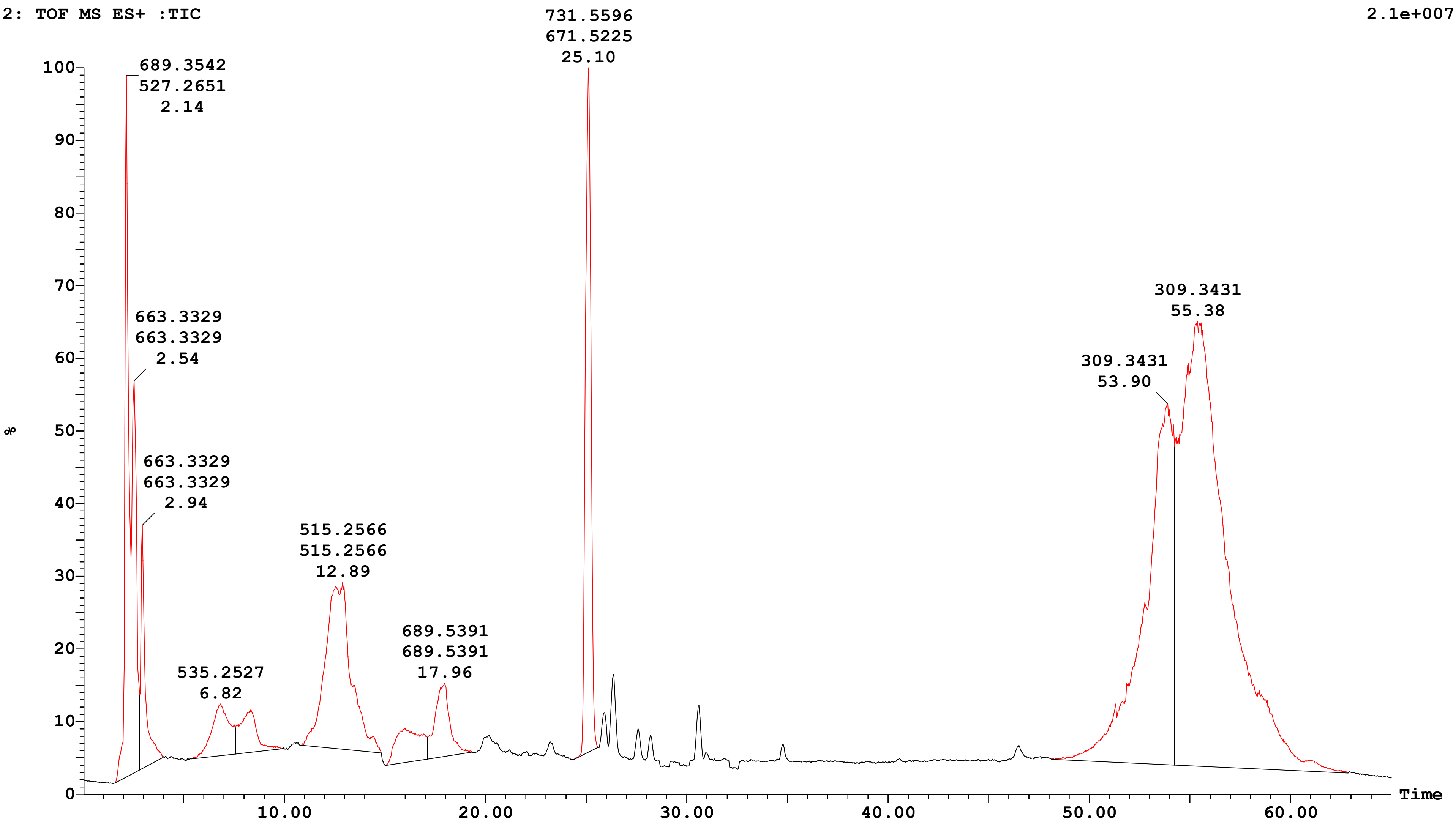


| Peak Number | Compound | Time | AreaAbs | Area %Total | Width | Height | Mass Found |
|---|---|---|---|---|---|---|---|
| 1 | | 2.14 | 4e+006 | 5.37 | 1 | 2e+007 | |
| 2 | | 2.54 | 3e+006 | 3.78 | 0 | 1e+007 | |
| 3 | | 2.94 | 2e+006 | 2.25 | 1 | 7e+006 | |
| 4 | | 6.82 | 2e+006 | 1.90 | 2 | 1e+006 | |
| 6 | | 8.33 | 1e+006 | 1.48 | 2 | 1e+006 | |

Sample: 34 Vial:1:C,2 ID:
File:AMIT_SAMPLE_B Date:16-Aug-2025 Time:06:18:36
Description:

Printed: Mon Aug 18 14:21:03 2025

| | | | | | |
|---|---|---|---|---|---|
| 8 | 12.89 | 7e+006 | 8.92 | 4 | 5e+006 |
| 9 | 15.99 | 1e+006 | 1.78 | 2 | 1e+006 |
| 11 | 17.96 | 2e+006 | 2.26 | 2 | 2e+006 |
| 12 | 25.10 | 6e+006 | 7.72 | 1 | 2e+007 |
| 16 | 53.90 | 2e+007 | 21.20 | 6 | 1e+007 |
| 18 | 55.38 | 4e+007 | 43.33 | 9 | 1e+007 |

Sample: 34 Vial:1:C,2 ID:
File:AMIT_SAMPLE_B Date:16-Aug-2025 Time:06:18:36
Description:

Printed: Mon Aug 18 14:21:03 2025

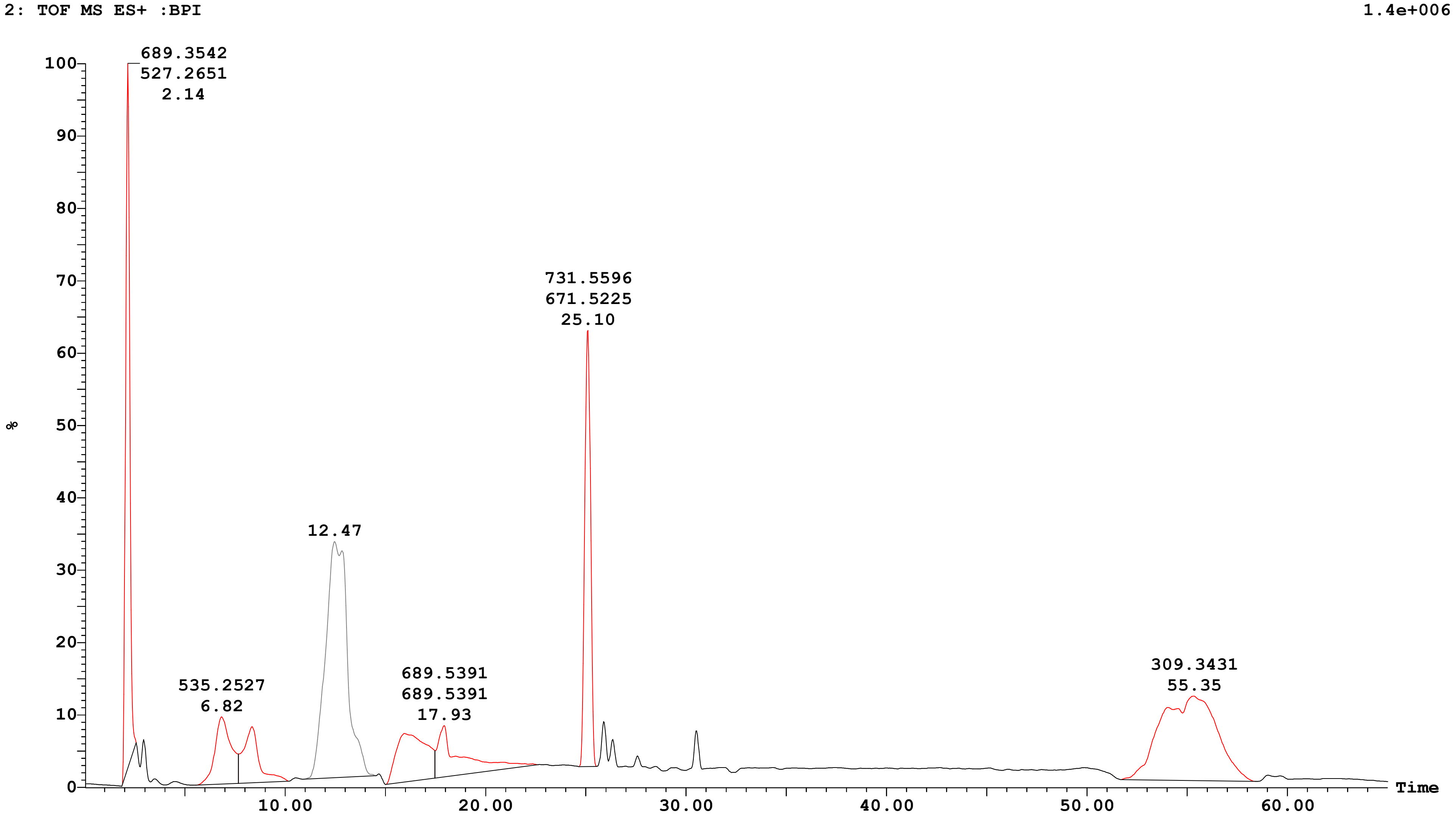

Sample: 34 Vial:1:C,2 ID:
File:AMIT_SAMPLE_B Date:16-Aug-2025 Time:06:18:36
Description:

Printed: Mon Aug 18 14:21:03 2025

| Peak ID | Time |
|---|---|
| 1 | 2.16 |

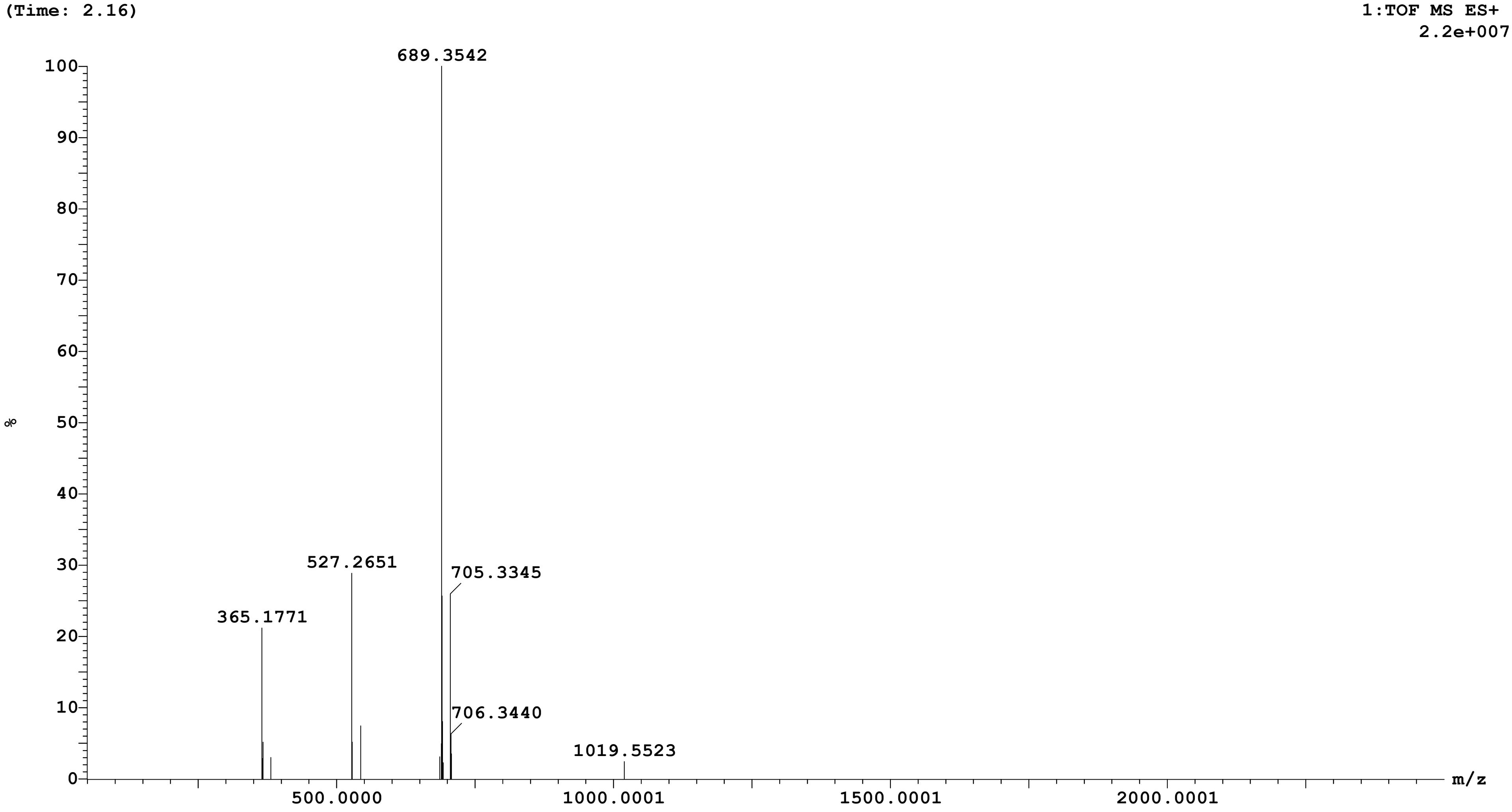

**Openlynx Report SAIF,PANJAB UNIVERSITY CHANDIGARH**

Sample: 34 | Vial:1:C,2 | ID:
File:AMIT_SAMPLE_B | Date:16-Aug-2025 | Time:06:18:36
Description:

Printed: Mon Aug 18 14:21:03 2025

| Peak ID | Time |
|---|---|
| 2 | 2.52 |

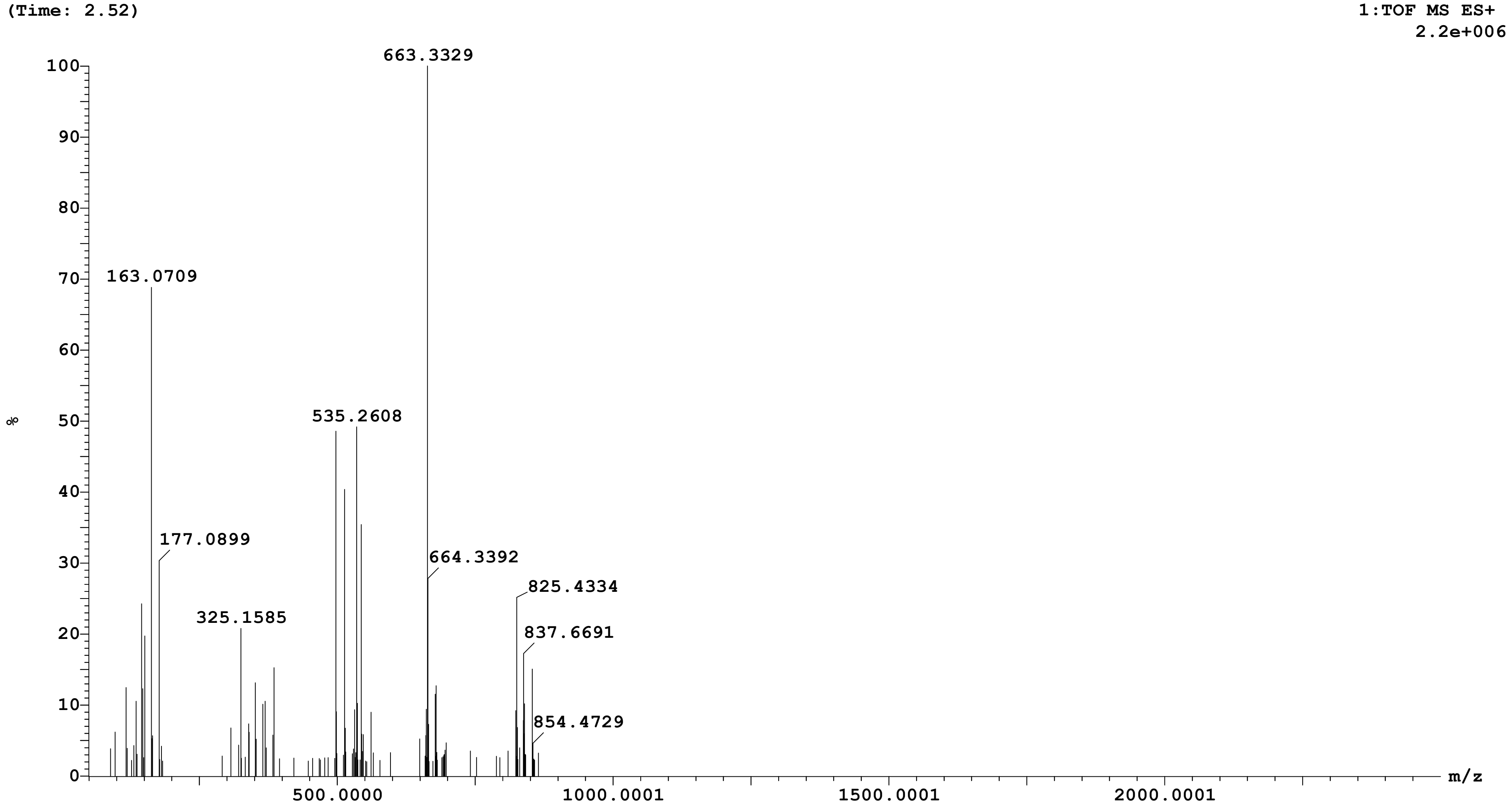

Sample: 34 Vial:1:C,2 ID:
File:AMIT_SAMPLE_B Date:16-Aug-2025 Time:06:18:36
Description:

Printed: Mon Aug 18 14:21:03 2025

| Peak ID | Time |
|---|---|
| 3 | 2.93 |

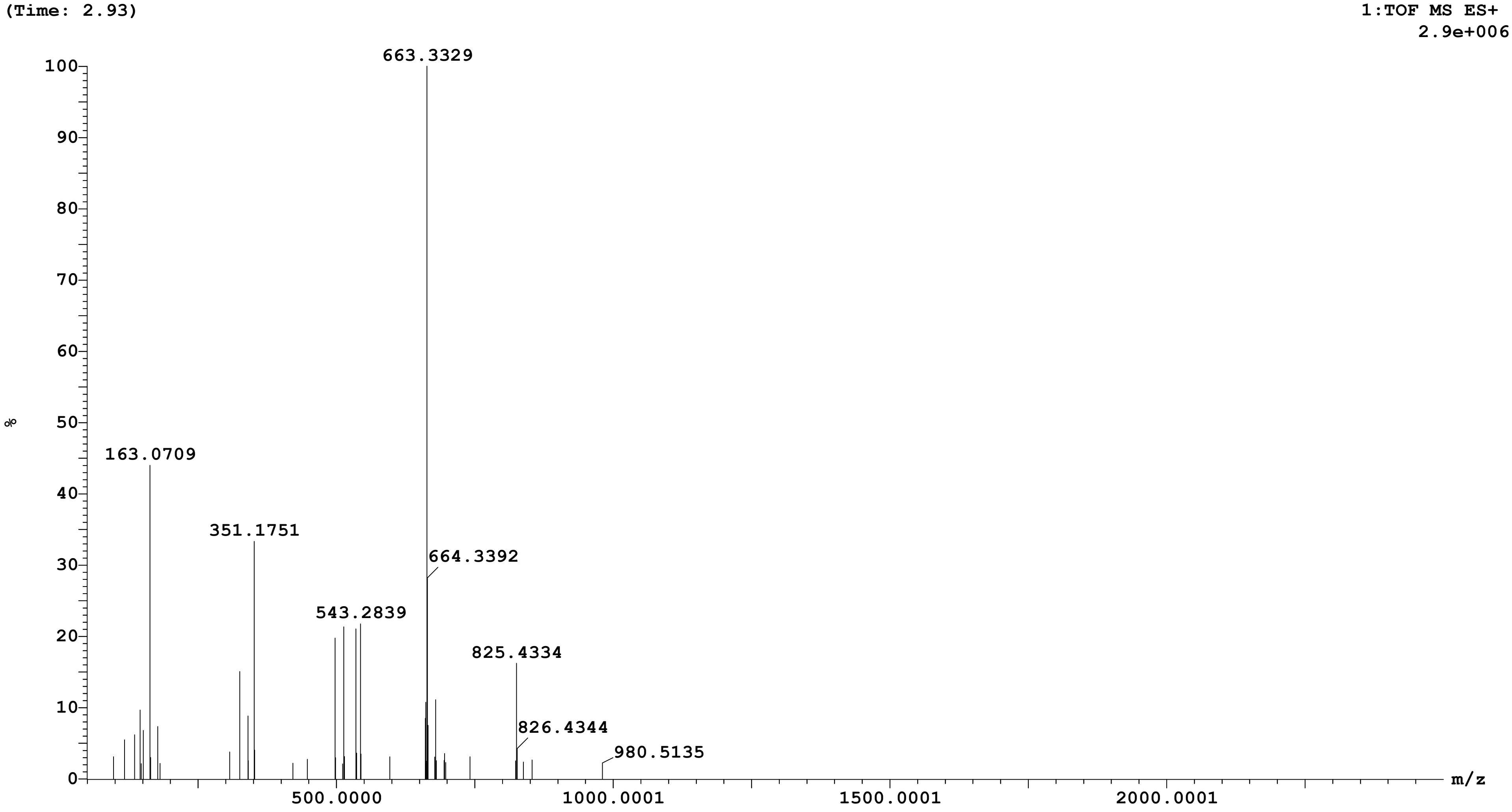

Sample: 34 Vial:1:C,2 ID:
File:AMIT_SAMPLE_B Date:16-Aug-2025 Time:06:18:36
Description:

Printed: Mon Aug 18 14:21:03 2025

| **Peak ID** | **Time** |
|---|---|
| 5 | 6.87 |

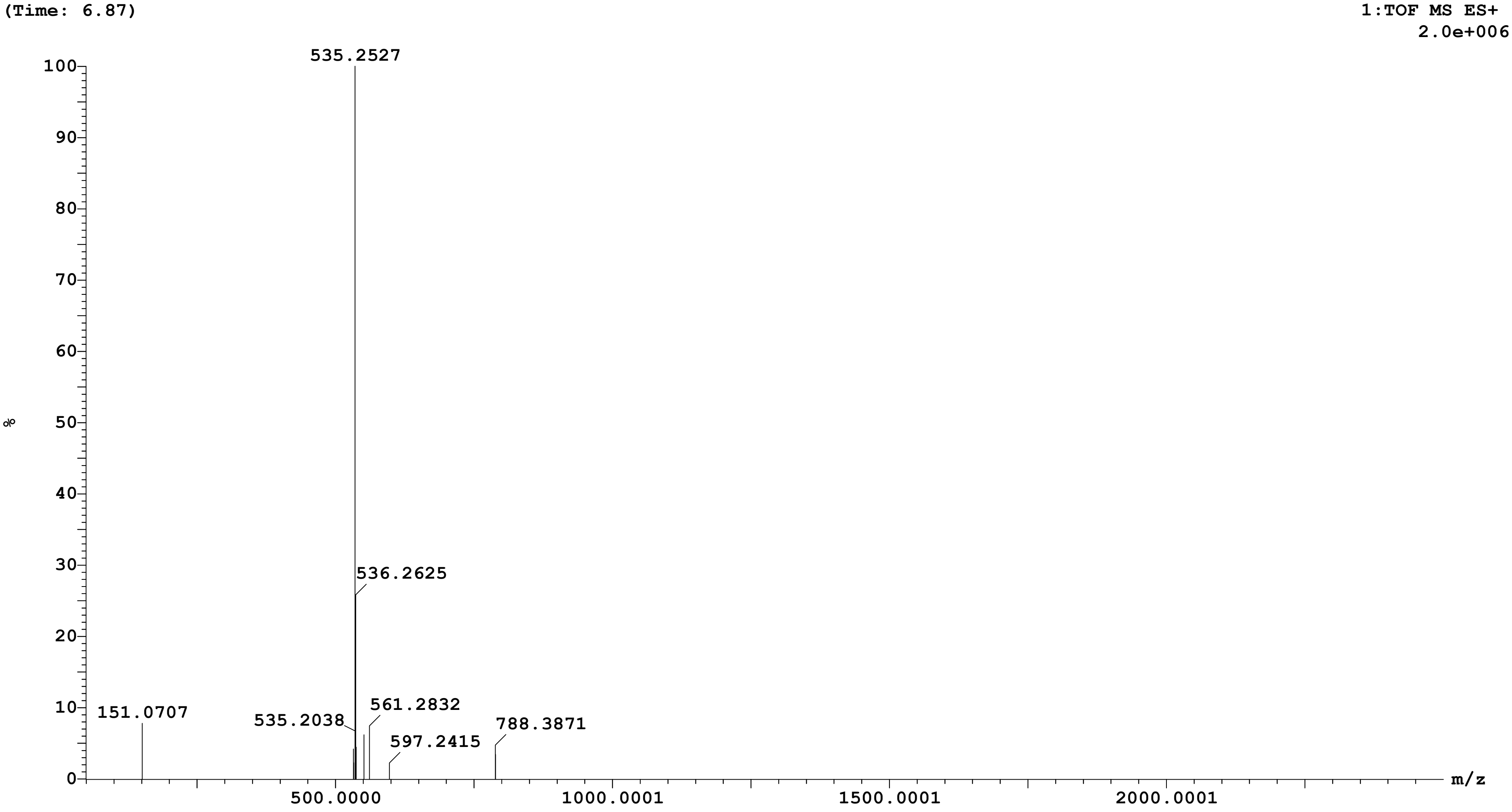

Sample: 34 Vial:1:C,2 ID:
File:AMIT_SAMPLE_B Date:16-Aug-2025 Time:06:18:36
Description:

Printed: Mon Aug 18 14:21:03 2025

| **Peak ID** | **Time** |
|---|---|
| 6 | 8.35 |

(Time: 8.35)

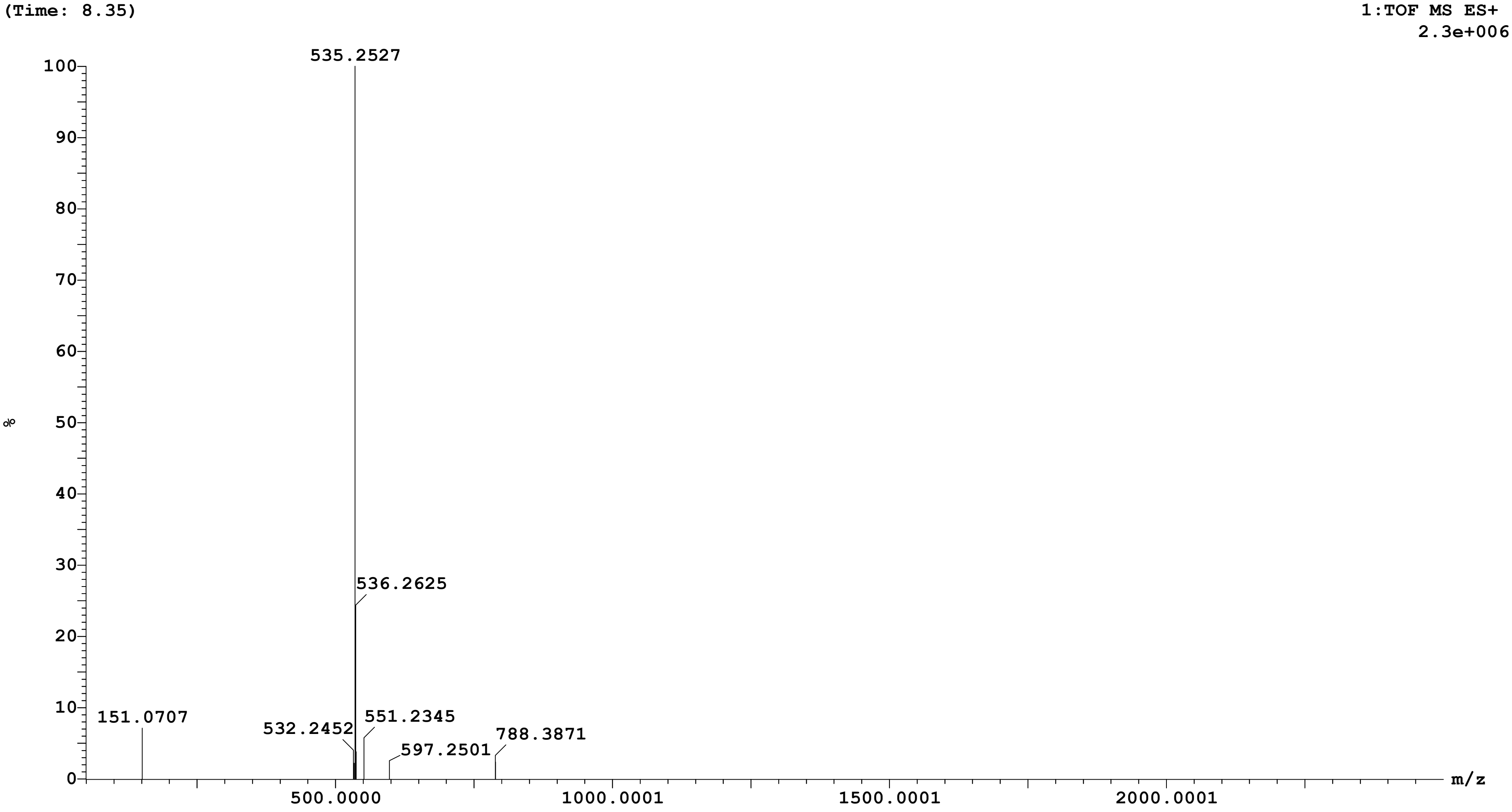

Sample: 34 Vial:1:C,2 ID:
File:AMIT_SAMPLE_B Date:16-Aug-2025 Time:06:18:36
Description:

Printed: Mon Aug 18 14:21:03 2025

| Peak ID | Time |
| --- | --- |
| 8 | 12.88 |

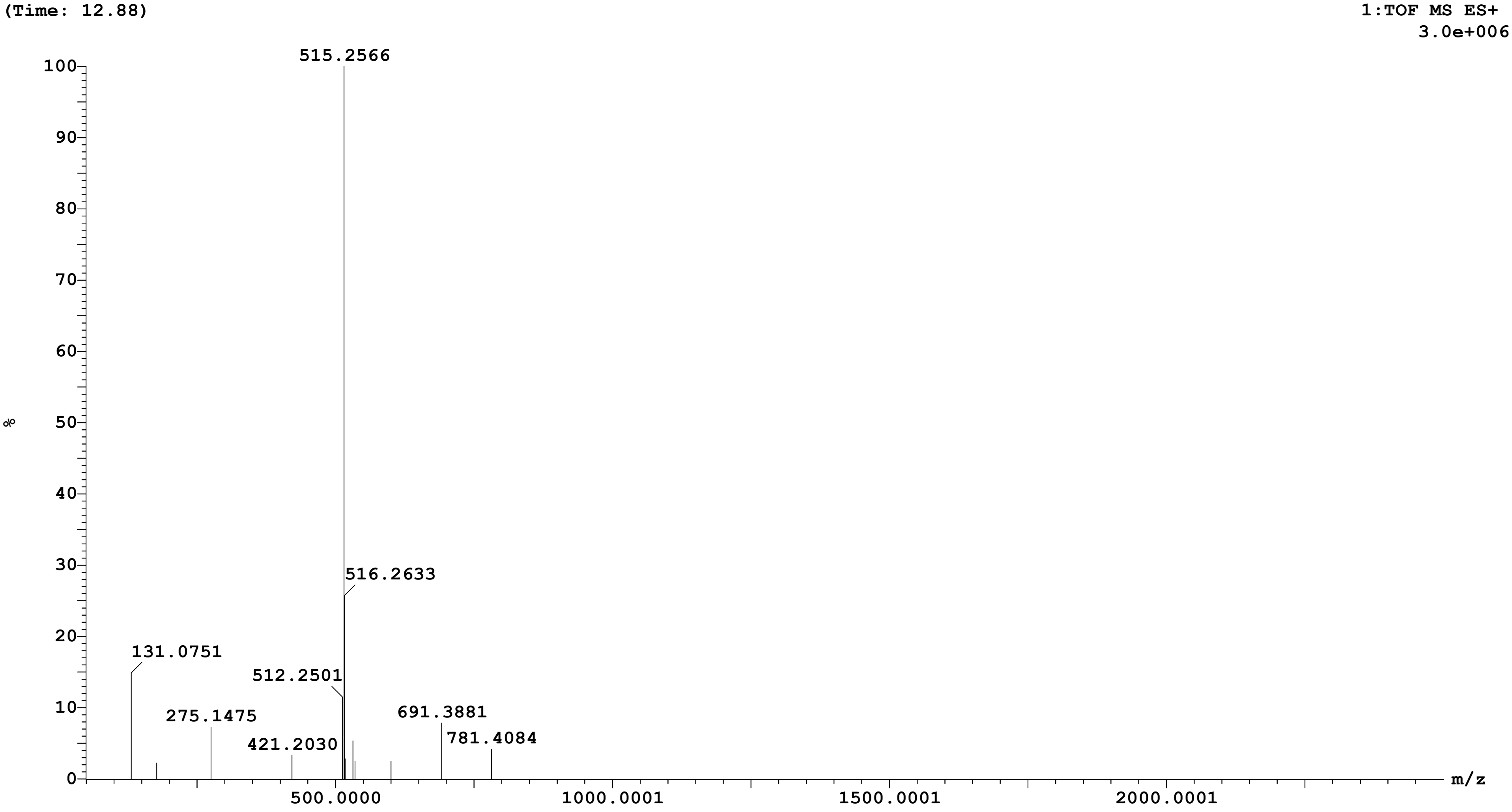

Sample: 34 Vial:1:C,2 ID:
File:AMIT_SAMPLE_B Date:16-Aug-2025 Time:06:18:36
Description:

Printed: Mon Aug 18 14:21:03 2025

| Peak ID | Time |
|---|---|
| 9 | 15.97 |

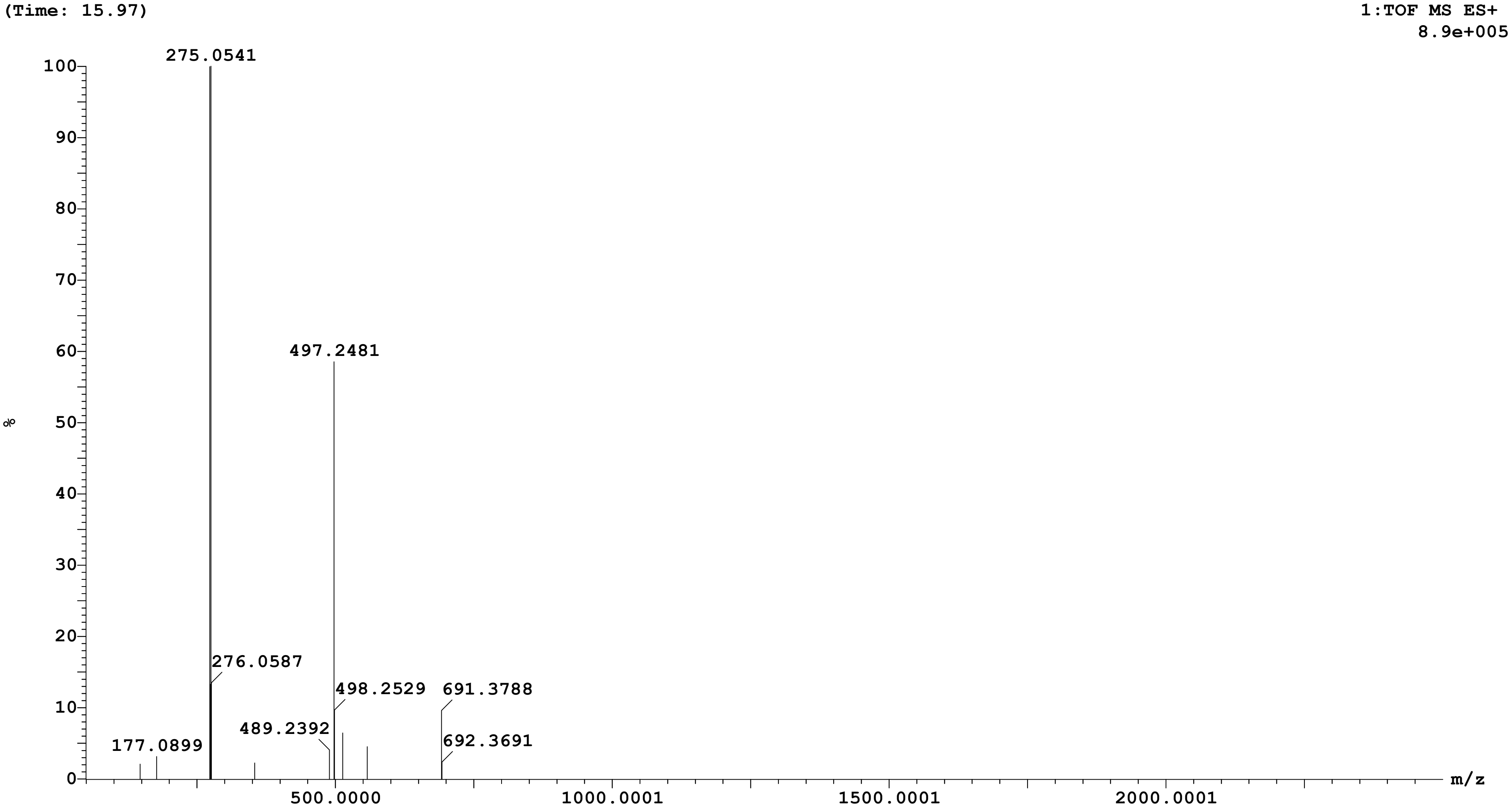

Sample: 34 Vial:1:C,2 ID:
File:AMIT_SAMPLE_B Date:16-Aug-2025 Time:06:18:36
Description:

Printed: Mon Aug 18 14:21:03 2025

| **Peak ID** | **Time** |
|---|---|
| 11 | 17.94 |

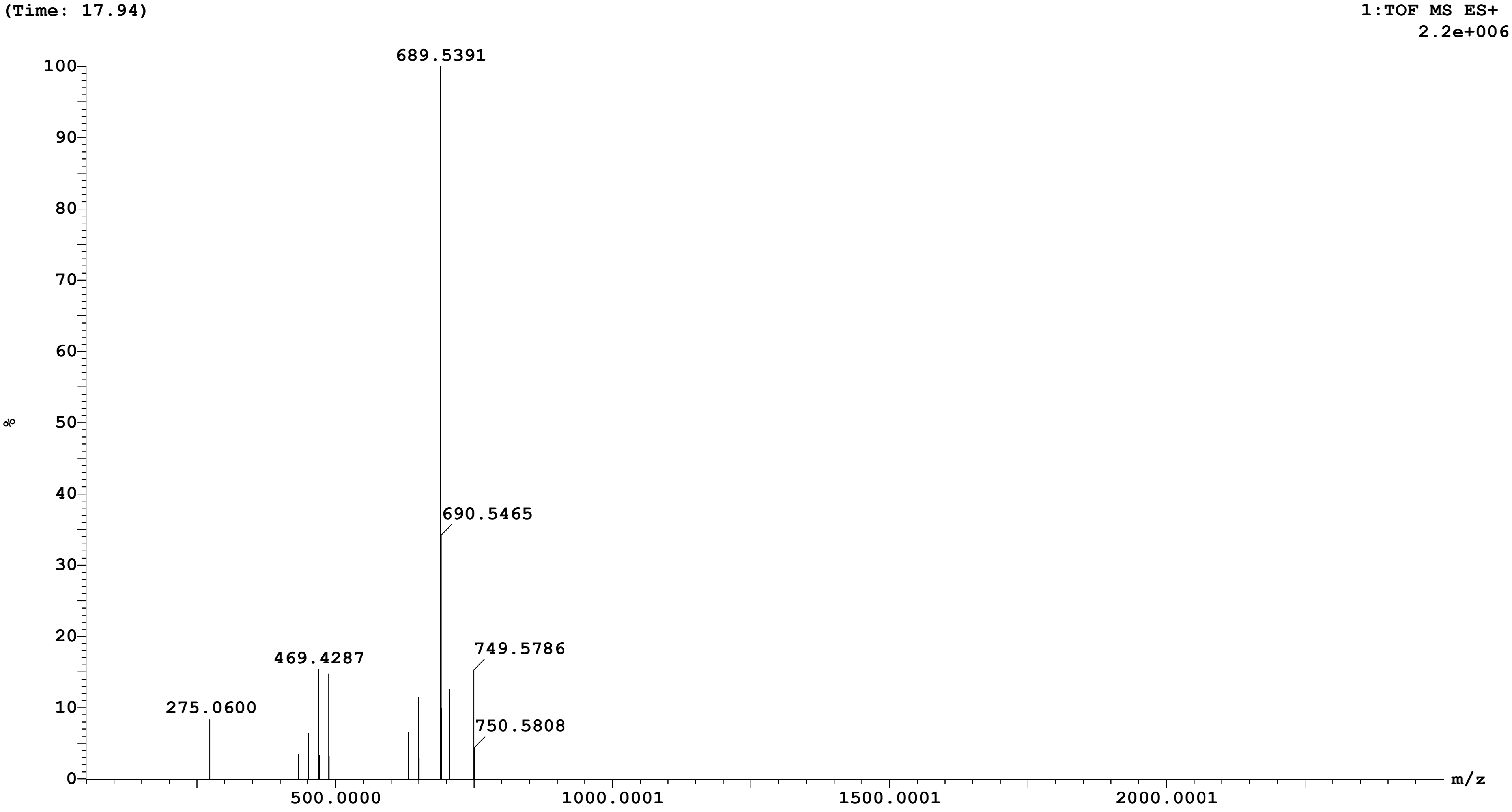

Sample: 34 Vial:1:C,2 ID:
File:AMIT_SAMPLE_B Date:16-Aug-2025 Time:06:18:36
Description:

Printed: Mon Aug 18 14:21:03 2025

| Peak ID | Time |
|---|---|
| 12 | 25.12 |

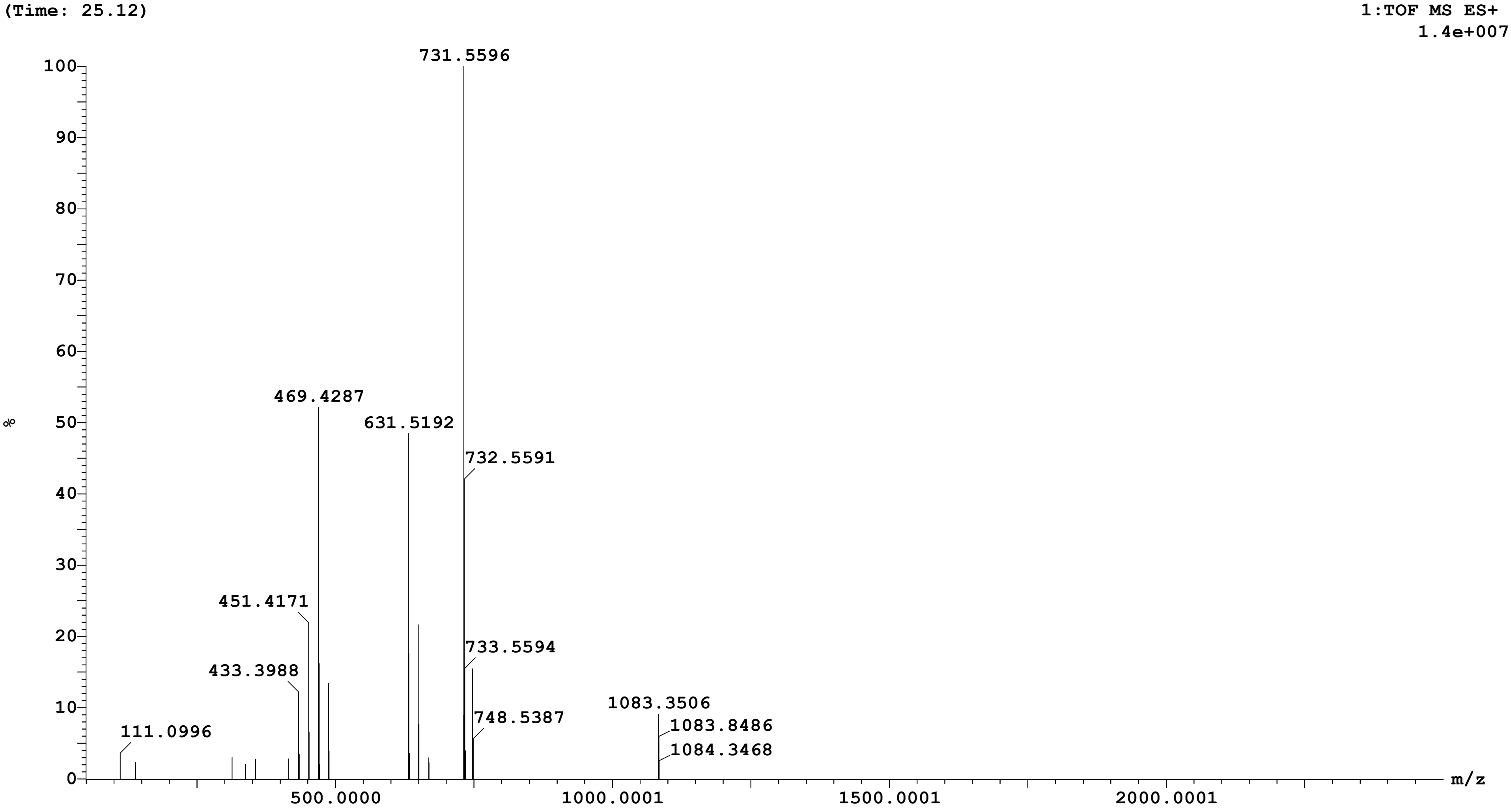

Sample: 34 Vial:1:C,2 ID:
File:AMIT_SAMPLE_B Date:16-Aug-2025 Time:06:18:36
Description:

Printed: Mon Aug 18 14:21:03 2025

| Peak ID | Time |
|---|---|
| 15 | 53.82 |

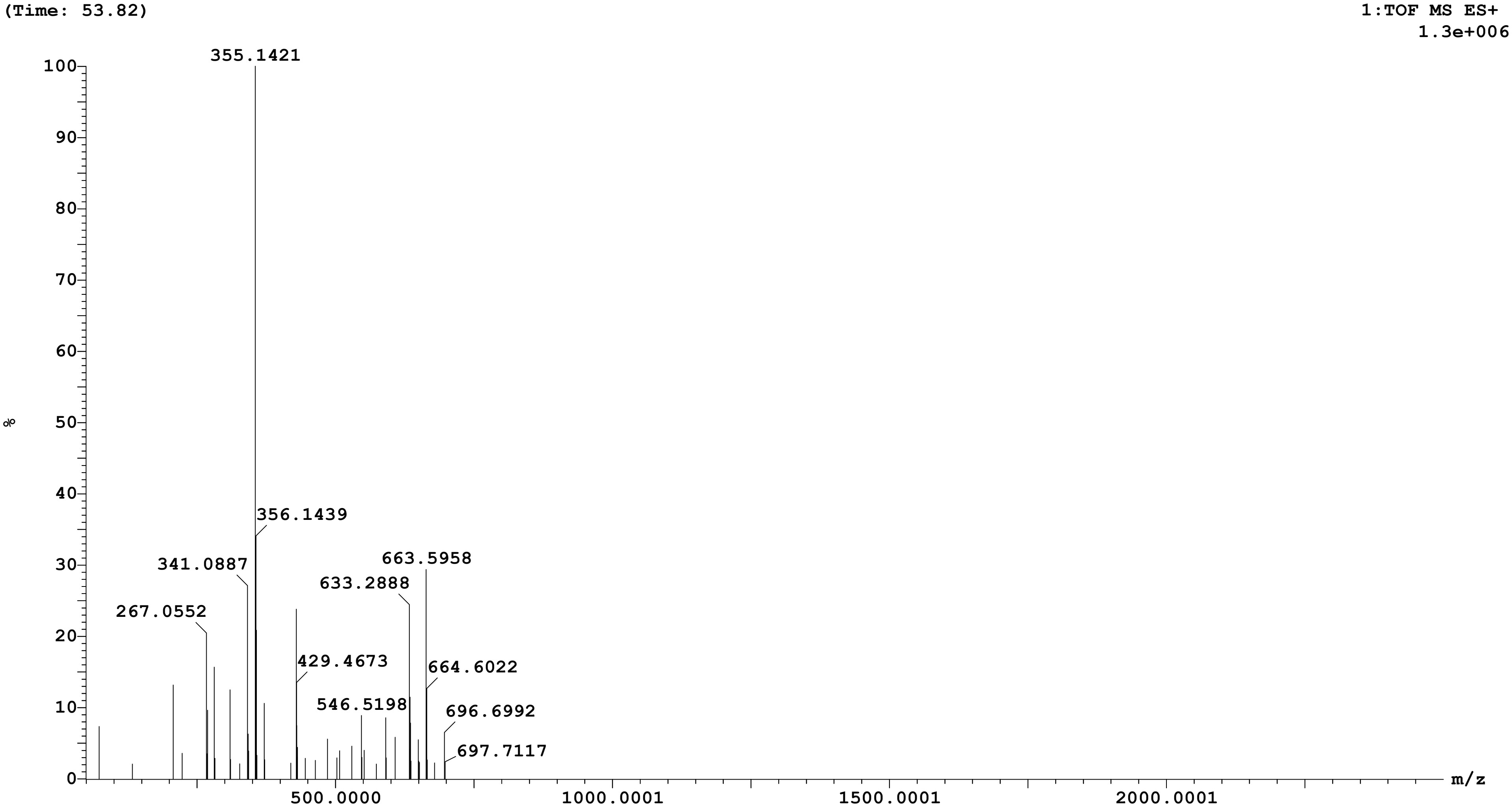

Sample: 34 Vial:1:C,2 ID:
File:AMIT_SAMPLE_B Date:16-Aug-2025 Time:06:18:36
Description:

Printed: Mon Aug 18 14:21:03 2025

| Peak ID | Time |
|---|---|
| 17 | 54.87 |

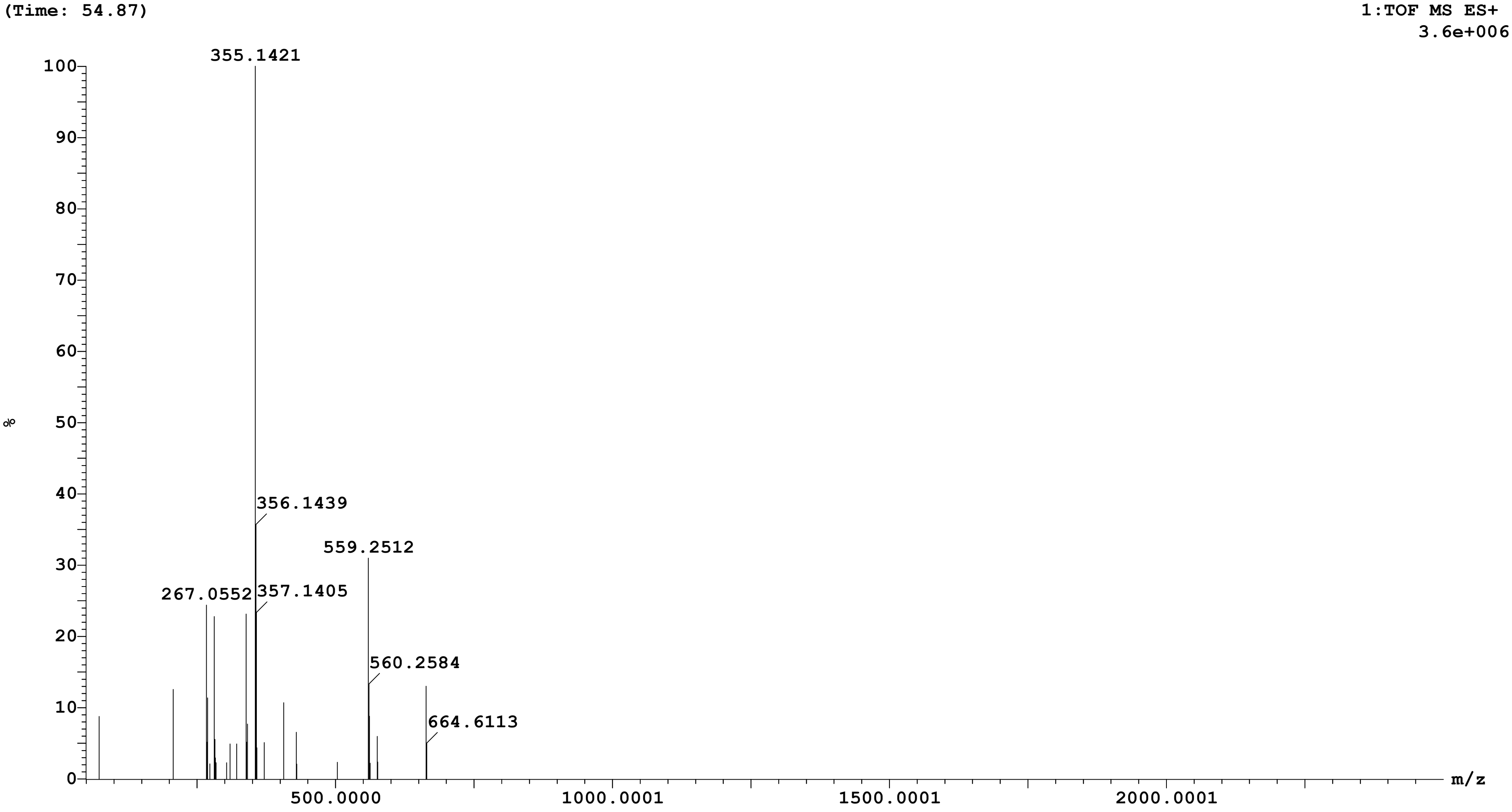

**Openlynx Report SAIF,PANJAB UNIVERSITY CHANDIGARH**

Sample: 34 Vial:1:C,2 ID:
File:AMIT_SAMPLE_B Date:16-Aug-2025 Time:06:18:36
Description:

Printed: Mon Aug 18 14:21:03 2025

| Peak ID | Time |
|---|---|
| 1 | 2.16 |

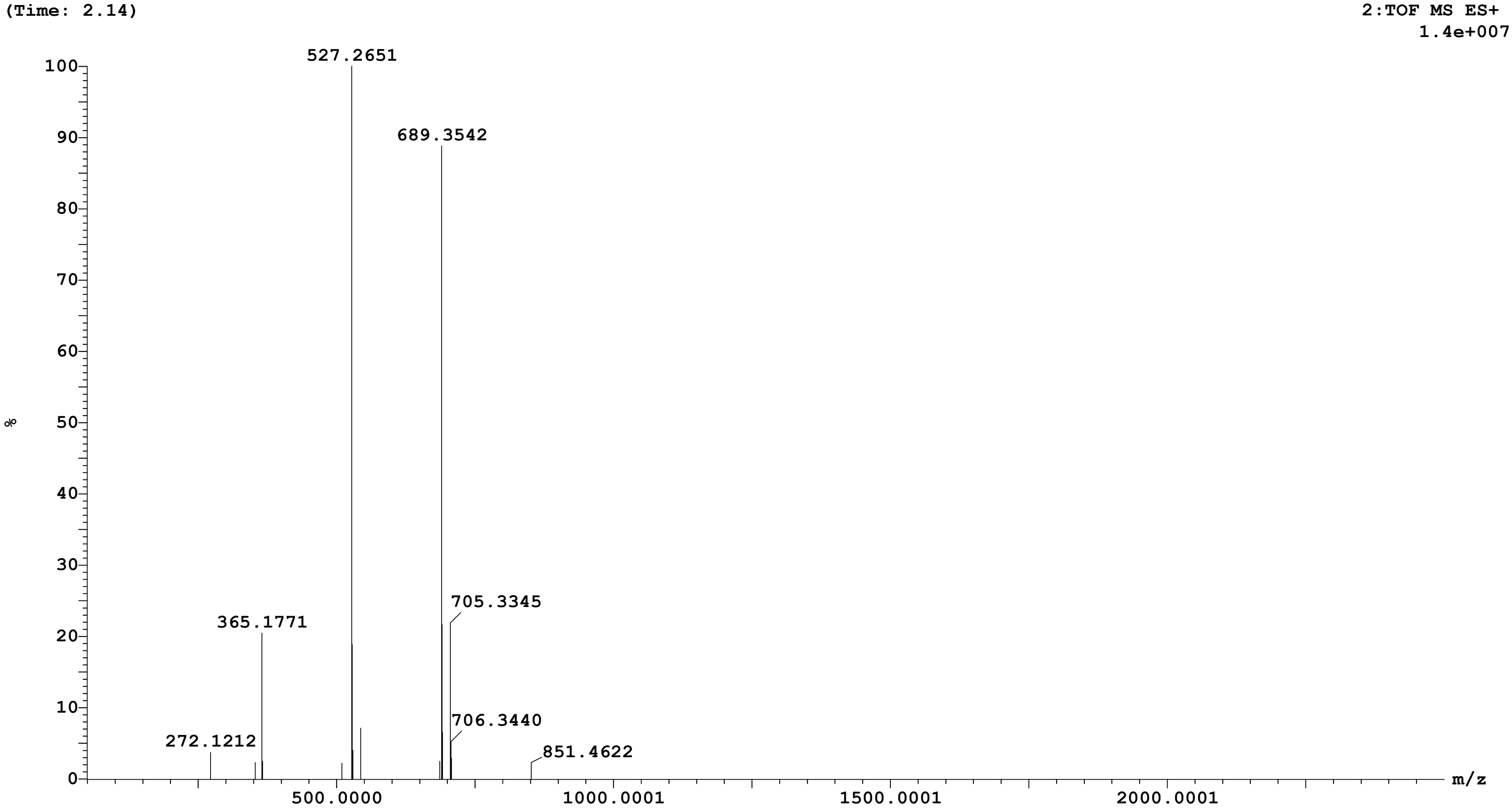

Sample: 34 Vial:1:C,2 ID:
File:AMIT_SAMPLE_B Date:16-Aug-2025 Time:06:18:36
Description:

Printed: Mon Aug 18 14:21:03 2025

| **Peak ID** | **Time** |
|---|---|
| 2 | 2.52 |

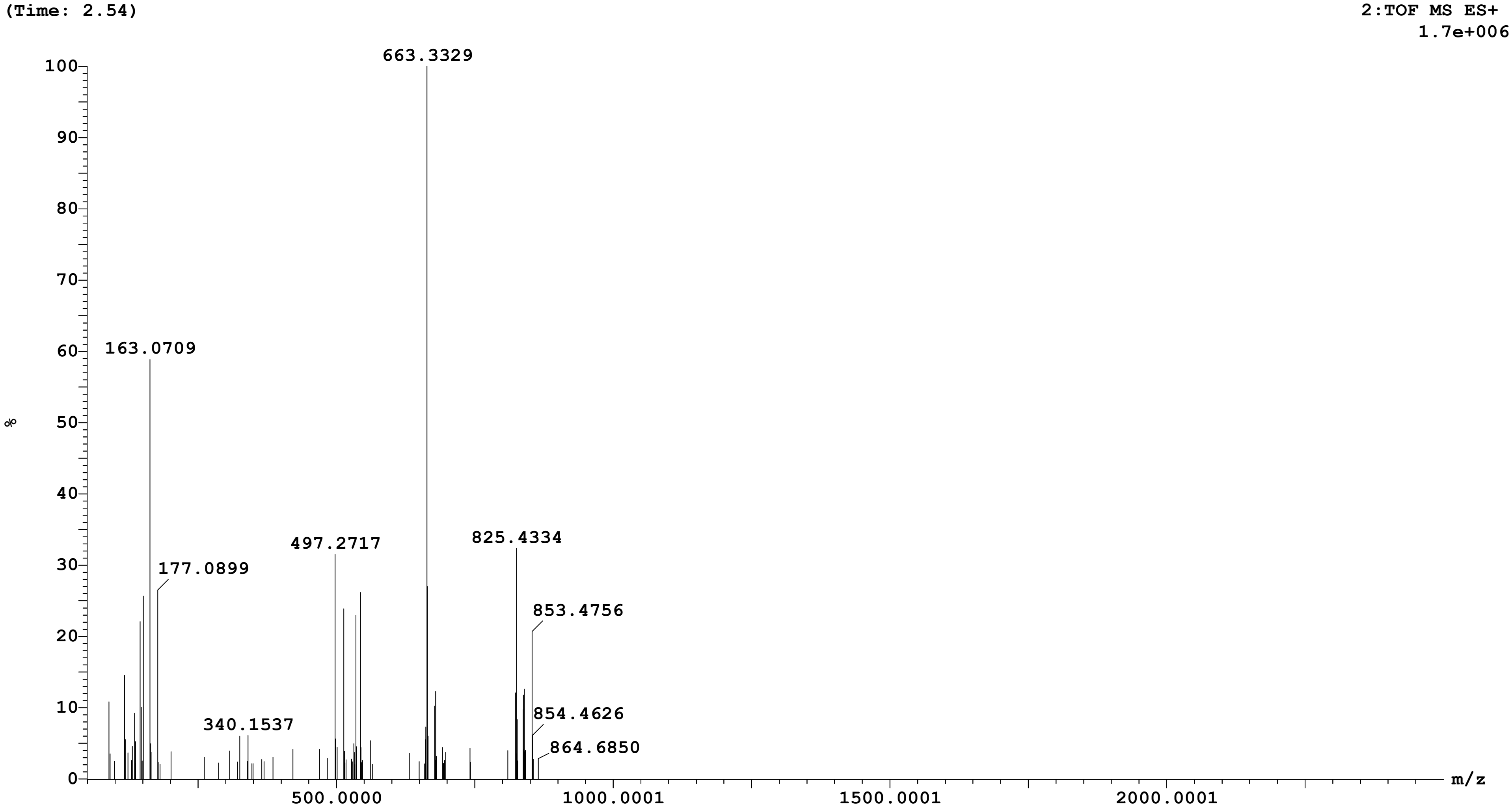

Sample: 34 Vial:1:C,2 ID:
File:AMIT_SAMPLE_B Date:16-Aug-2025 Time:06:18:36
Description:

Printed: Mon Aug 18 14:21:03 2025

| **Peak ID** | **Time** |
|---|---|
| 3 | 2.93 |

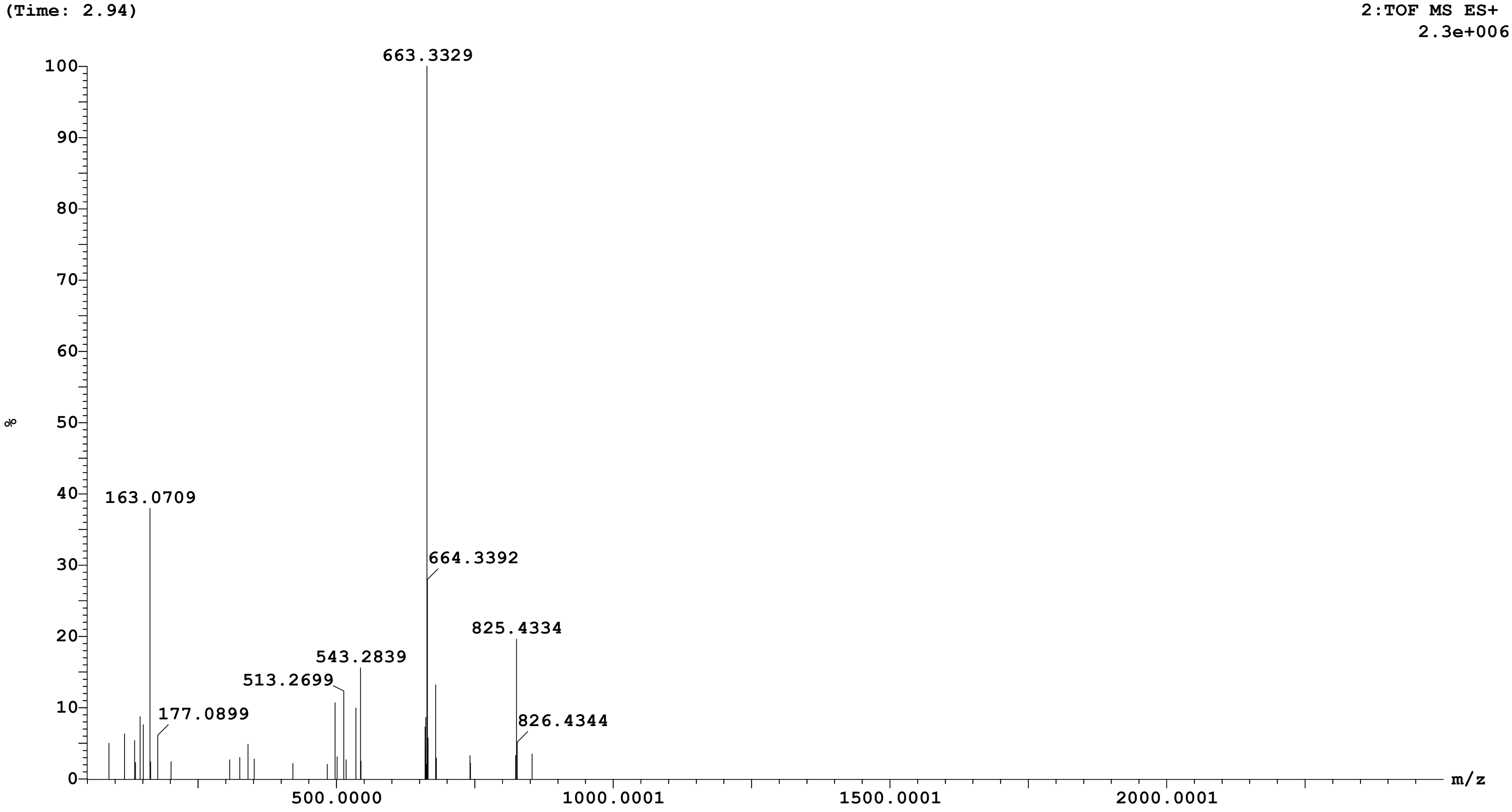

Sample: 34 Vial:1:C,2 ID:
File:AMIT_SAMPLE_B Date:16-Aug-2025 Time:06:18:36
Description:

Printed: Mon Aug 18 14:21:03 2025

| **Peak ID** | **Time** |
|---|---|
| 4 | 6.84 |

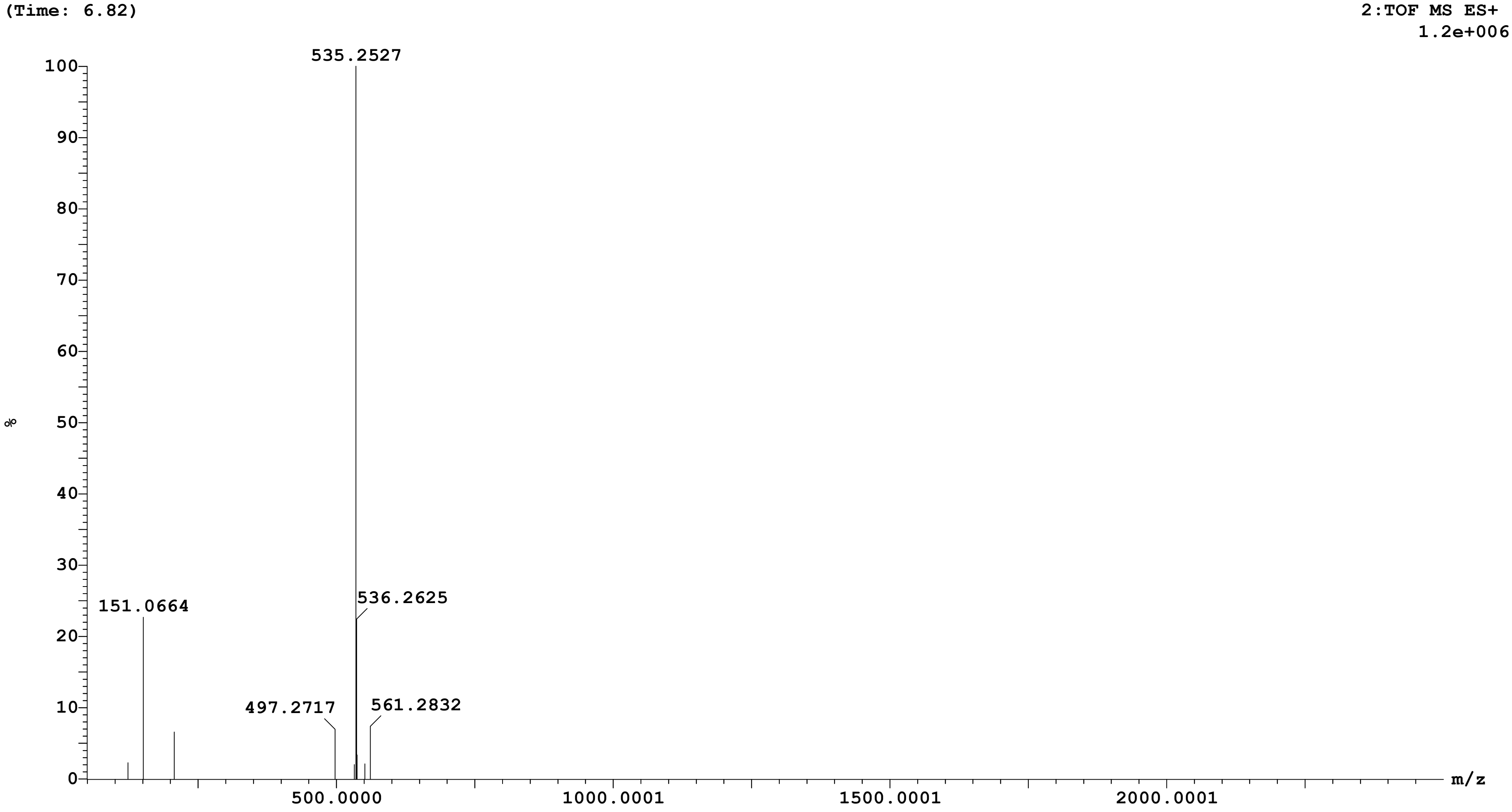

Sample: 34 Vial:1:C,2 ID:
File:AMIT_SAMPLE_B Date:16-Aug-2025 Time:06:18:36
Description:

Printed: Mon Aug 18 14:21:03 2025

| Peak ID | Time |
|---|---|
| 6 | 8.35 |

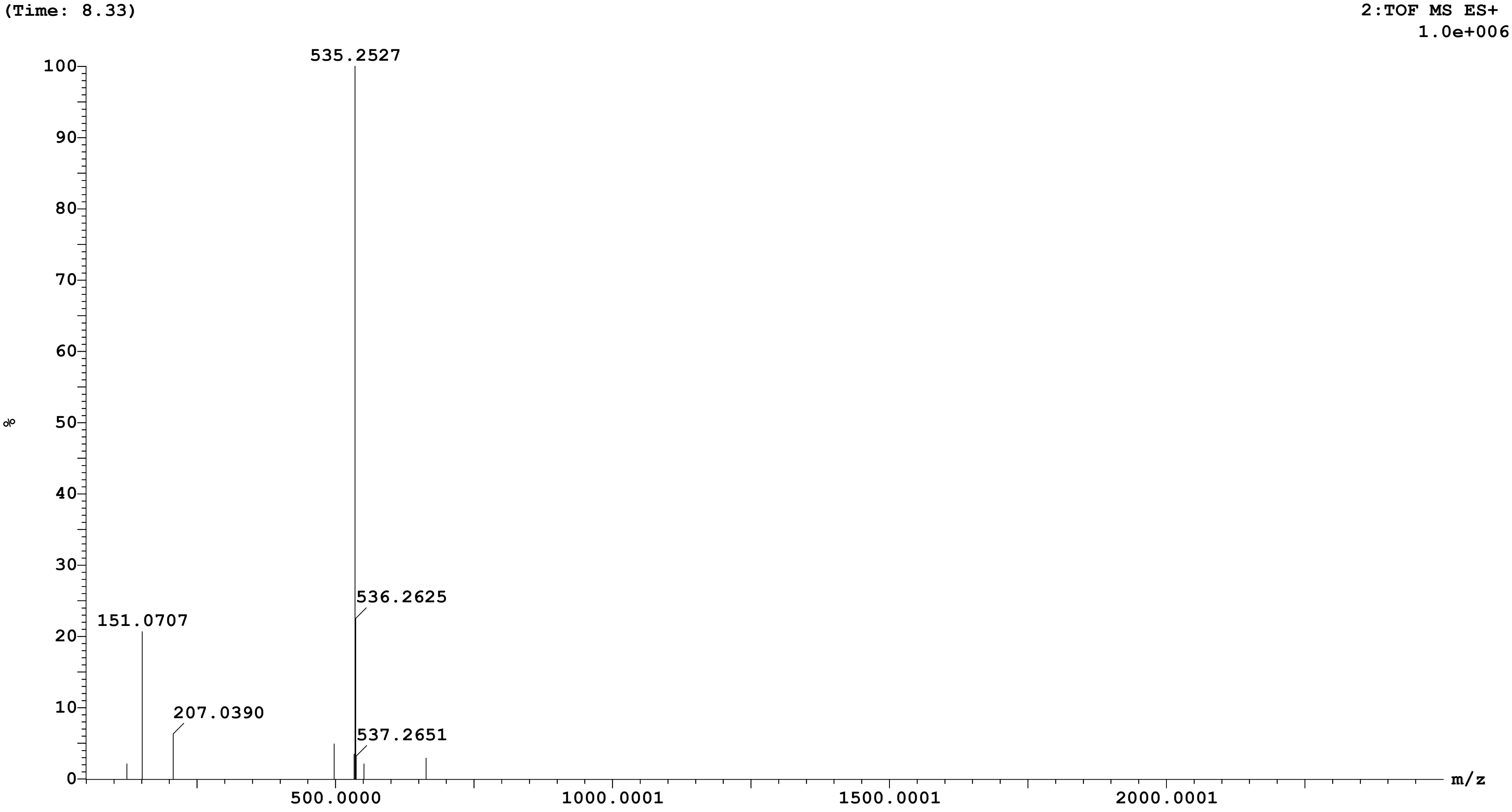

Sample: 34 Vial:1:C,2 ID:
File:AMIT_SAMPLE_B Date:16-Aug-2025 Time:06:18:36
Description:

Printed: Mon Aug 18 14:21:03 2025

| Peak ID | Time |
|---|---|
| 8 | 12.88 |

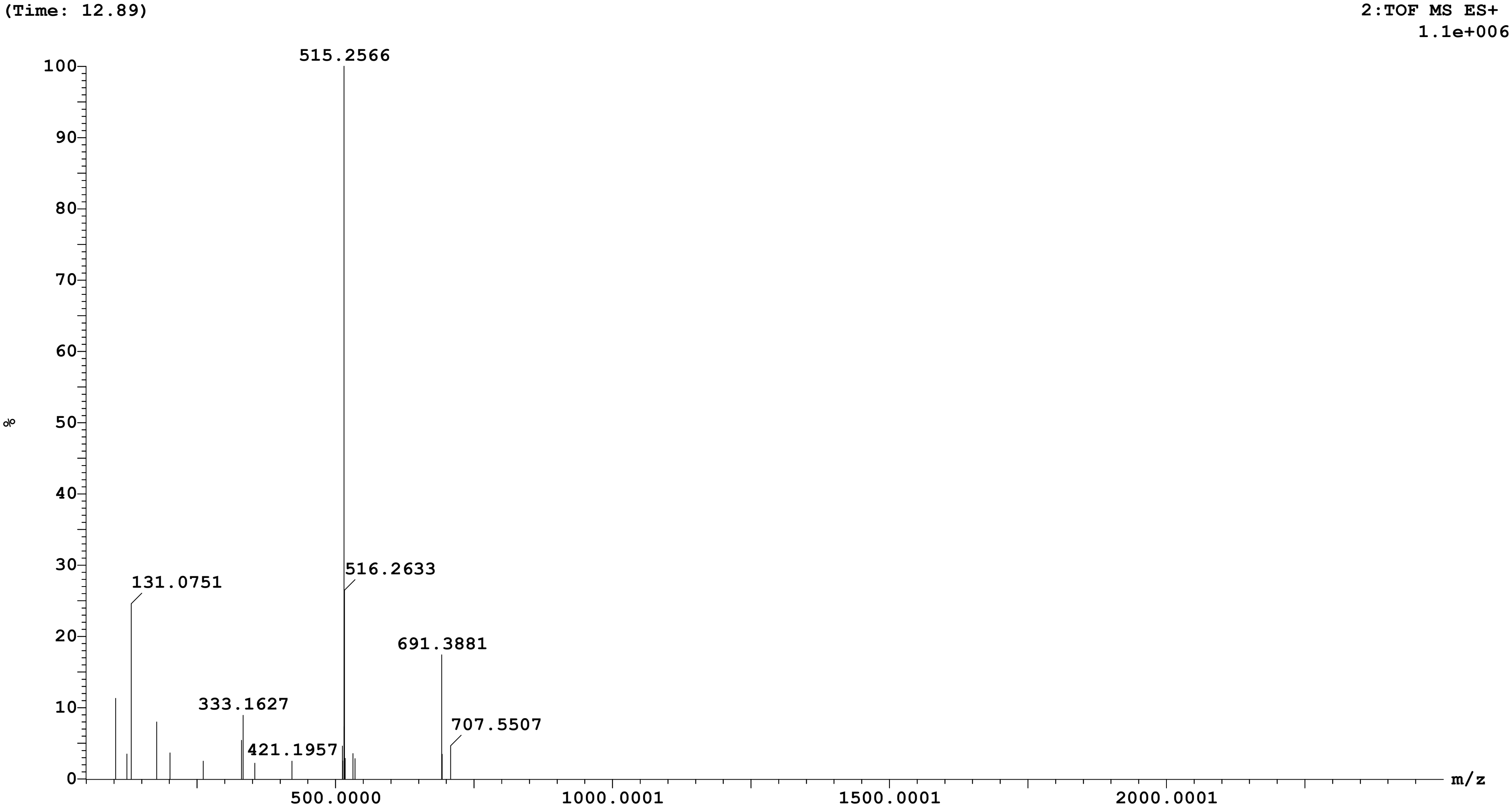

Sample: 34 Vial:1:C,2 ID:
File:AMIT_SAMPLE_B Date:16-Aug-2025 Time:06:18:36
Description:

Printed: Mon Aug 18 14:21:03 2025

| **Peak ID** | **Time** |
|---|---|
| 9 | 15.97 |

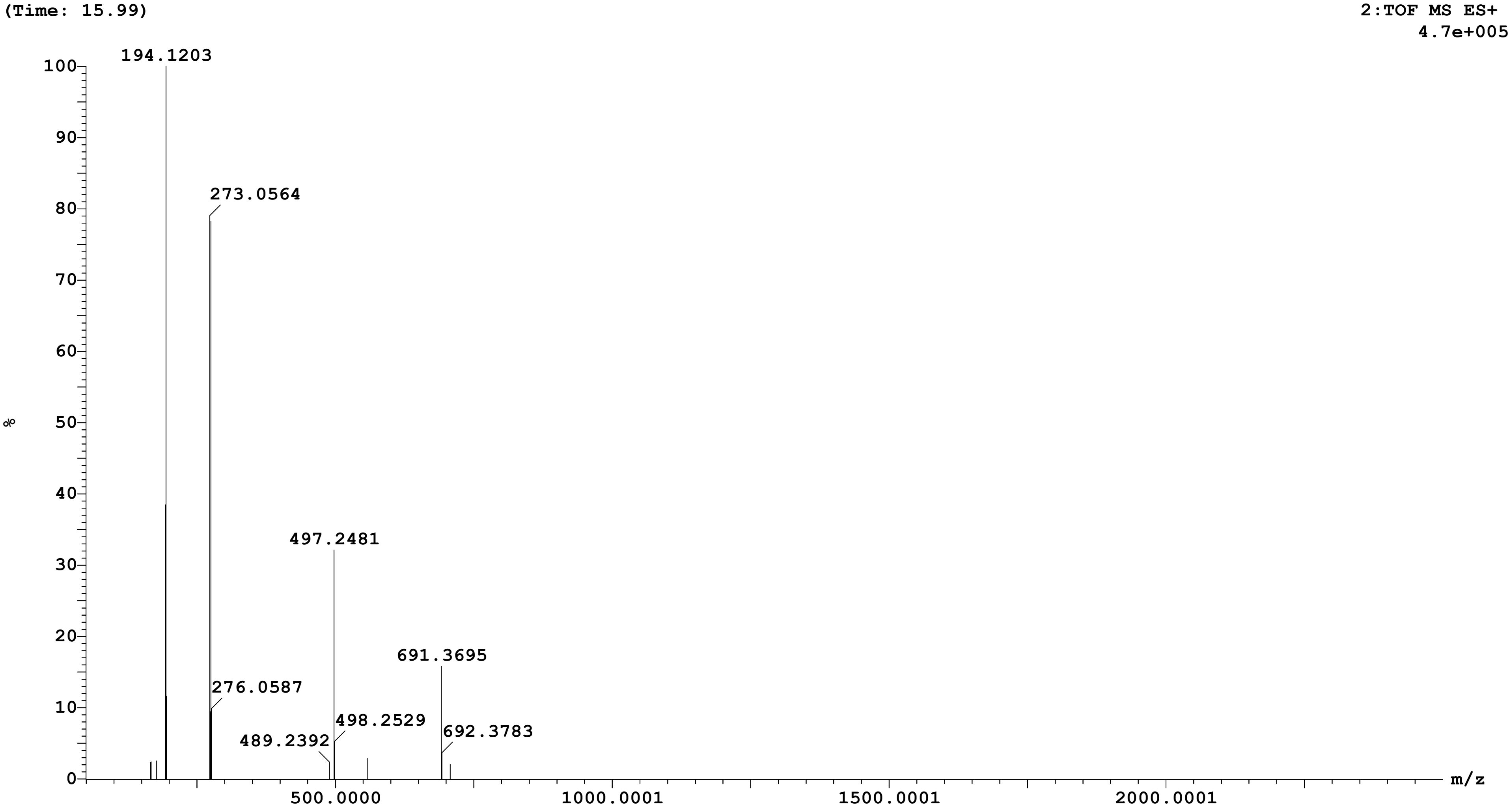

Sample: 34 Vial:1:C,2 ID:
File:AMIT_SAMPLE_B Date:16-Aug-2025 Time:06:18:36
Description:

Printed: Mon Aug 18 14:21:03 2025

| Peak ID | Time |
|---|---|
| 11 | 17.94 |

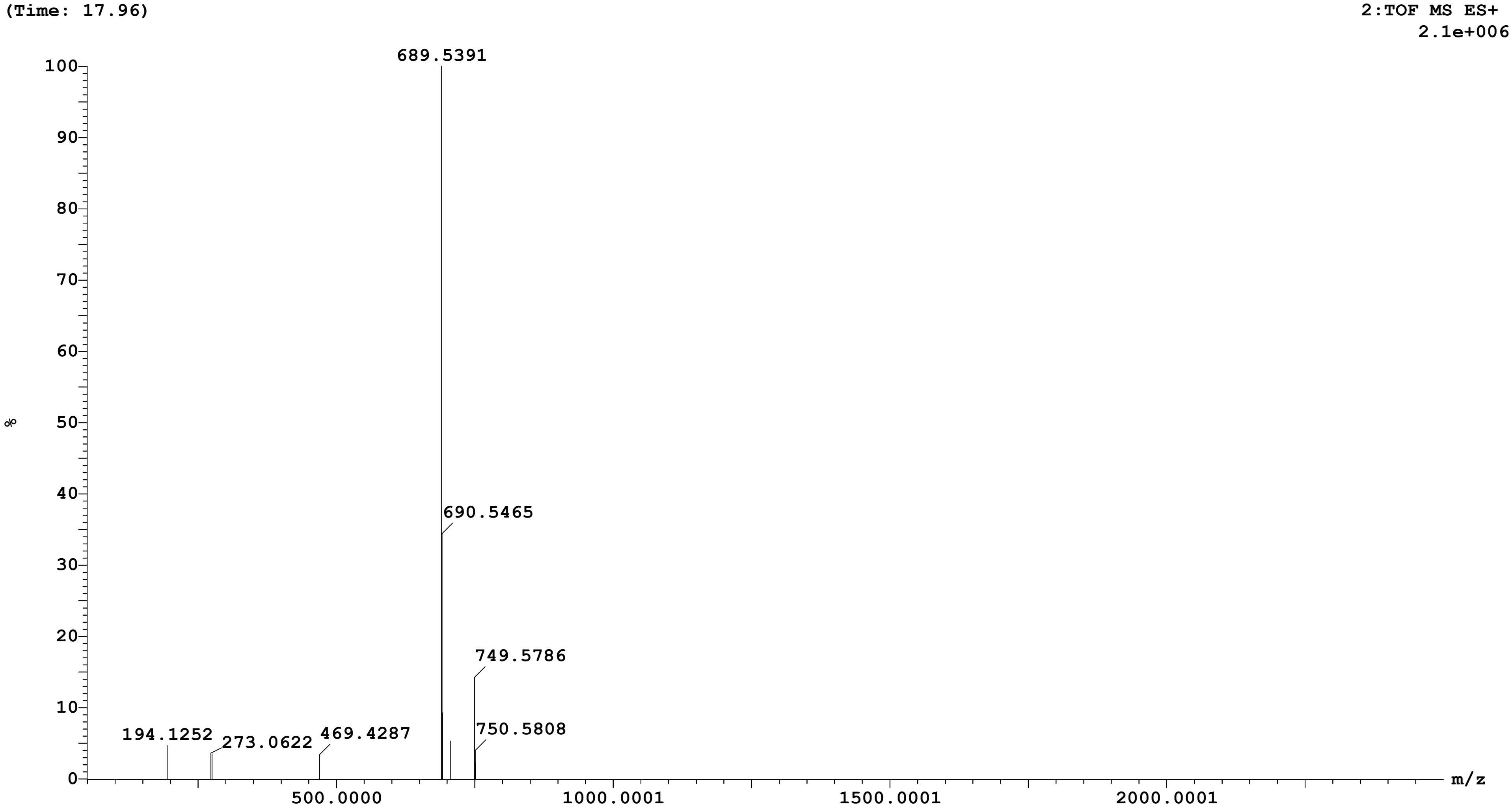

Sample: 34 Vial:1:C,2 ID:
File:AMIT_SAMPLE_B Date:16-Aug-2025 Time:06:18:36
Description:

Printed: Mon Aug 18 14:21:03 2025

| **Peak ID** | **Time** |
|---|---|
| 12 | 25.12 |

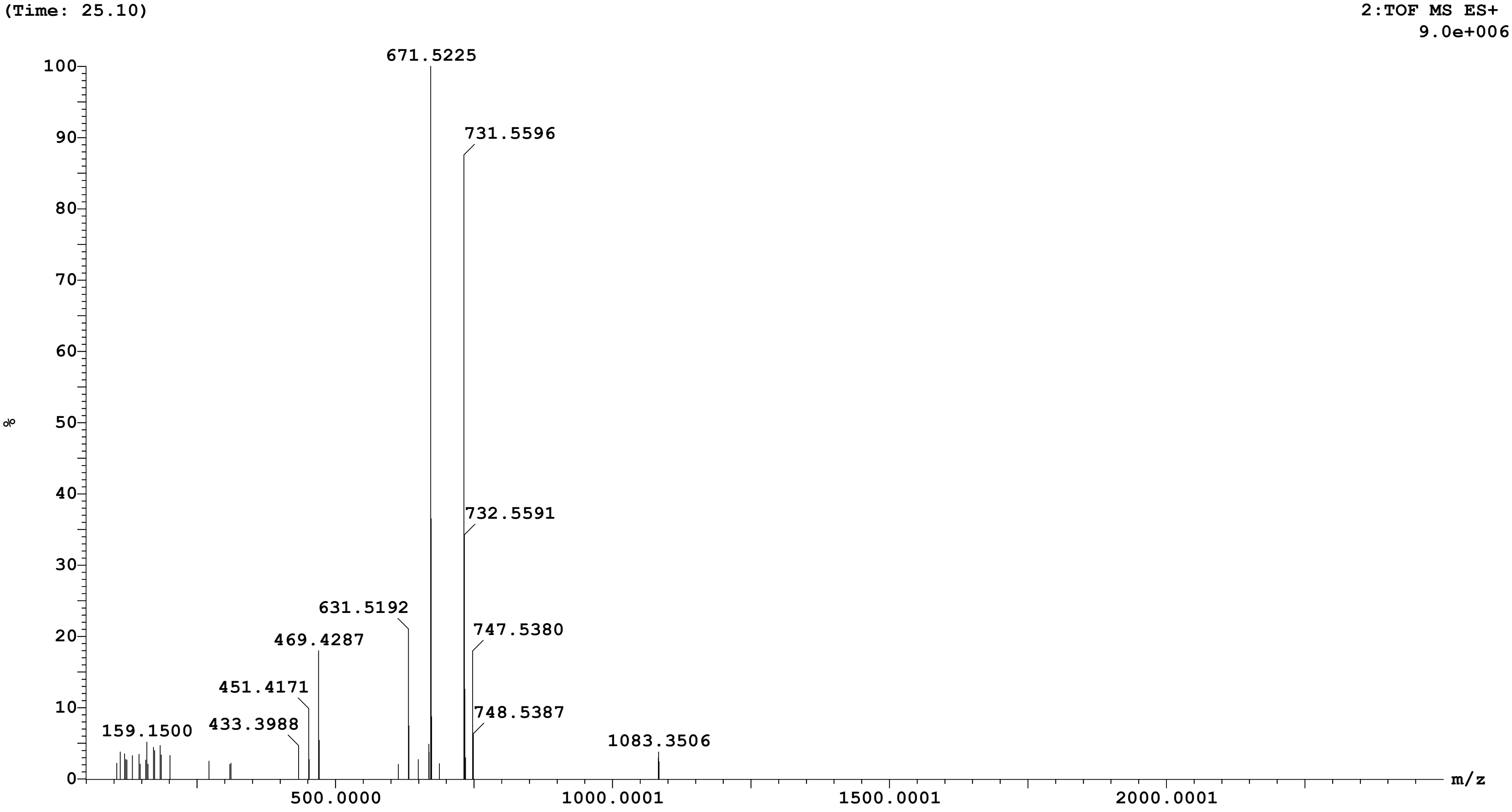

Sample: 34 Vial:1:C,2 ID:
File:AMIT_SAMPLE_B Date:16-Aug-2025 Time:06:18:36
Description:

Printed: Mon Aug 18 14:21:03 2025

| Peak ID | Time |
|---|---|
| 16 | 53.90 |

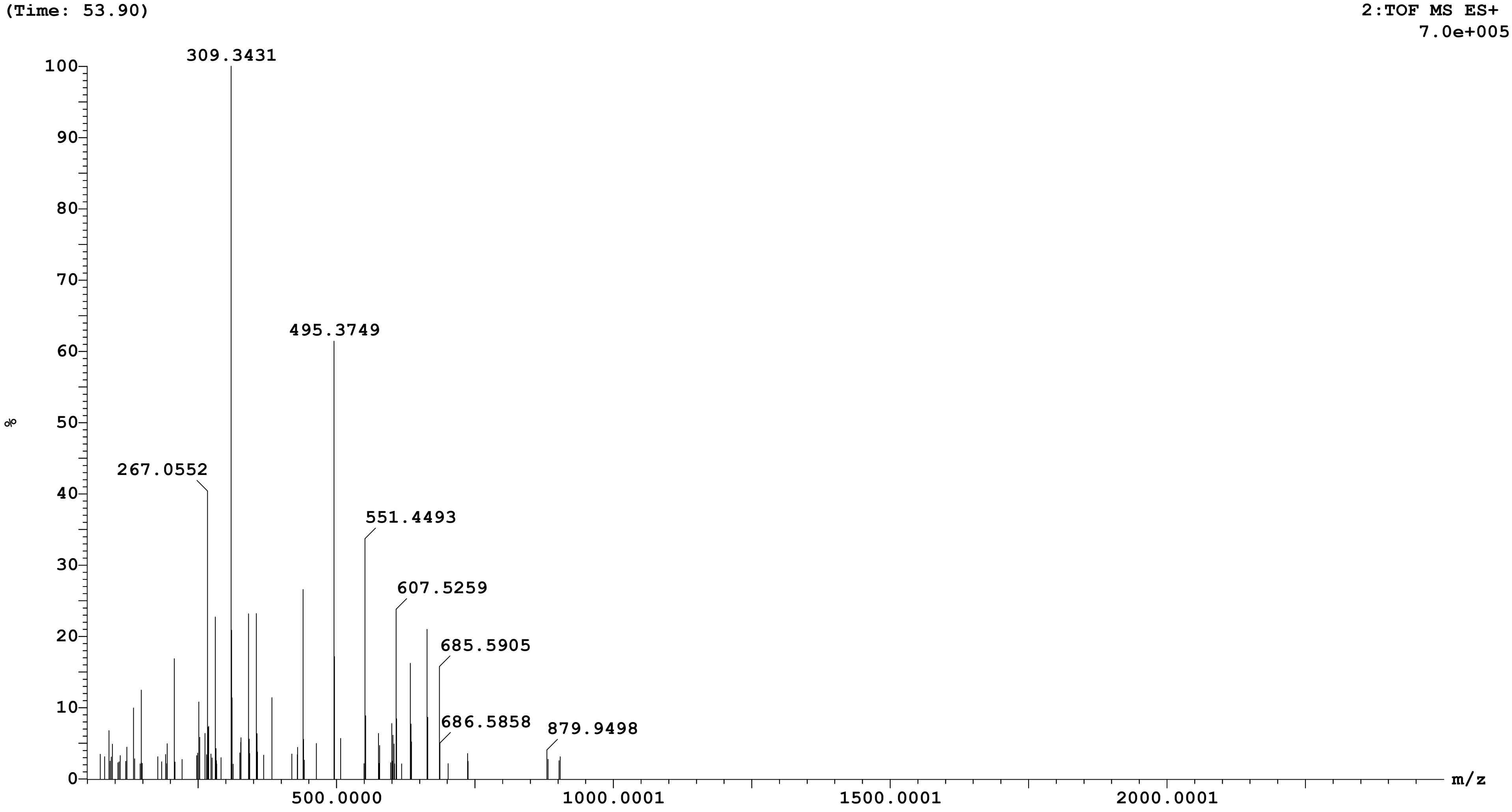

Sample: 34 Vial:1:C,2 ID:
File:AMIT_SAMPLE_B Date:16-Aug-2025 Time:06:18:36
Description:

Printed: Mon Aug 18 14:21:03 2025

| **Peak ID** | **Time** |
|---|---|
| 18 | 55.38 |

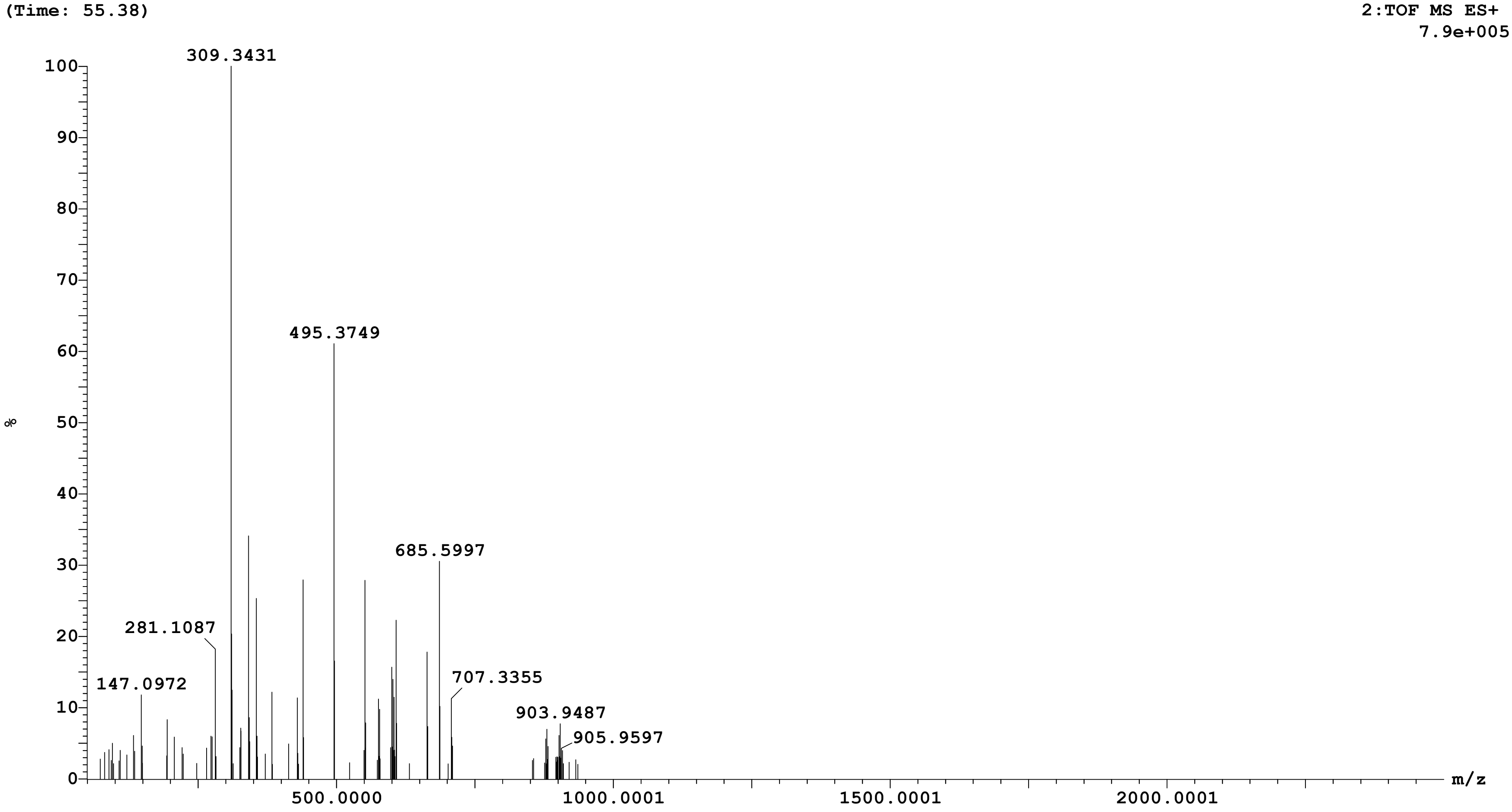
